# Developing High-Affinity Decoy Receptor Optimized for Multiple Myeloma and Diffuse Large B Cell Lymphoma Treatment

**DOI:** 10.1101/2021.05.03.442500

**Authors:** Yu Rebecca Miao, Kaushik Thakker, Can Cenik, Dadi Jiang, Kazue Mizuno, Chenjun Jia, Caiyun Grace Li, Hongjuan Zhao, Anh Diep, Jimmy Yu Xu, Xin Eric Zhang, Teddy Tat Chi Yang, Michaela Liedtke, Parveen Abidi, Wing-sze Leung, Albert C. Koong, Amato J. Giaccia

**Author notes:** Corresponding Authors: Amato J. Giaccia, Ph.D. Oxford Institute for Radiation Oncology University of Oxford Old Road Campus Research Building (ORCRB) Roosevelt Drive Oxford OX3 7DQ Yu Rebecca Miao, Ph.D. Department of Radiation Oncology Stanford University 269 Campus Drive, CCSR South Bldg. Rm 1250 Stanford CA 94305 USA. Yu Rebecca Miao Ph.D. and Kaushik Thakker Ph.D. Contributed Equally to This Study.

## Abstract

Recent T Cell therapies have been effective in the treatment of hematological cancers. However, immunotoxicity and treatment relapse pose significant clinical challenges. Here, we revealed distinctive requirement for neutralizing TNF receptor ligands APRIL and BAFF in MM and DLBCL. Furthermore, we investigated the use of BCMA decoy receptor (sBCMA-Fc) as a therapeutic inhibitor of ARPIL and BAFF. While wild-type sBCMA-Fc successfully blocked APRIL signaling with picomolar binding affinity, inhibiting tumor growth in MM models, it lacked efficacy in inhibiting DLBCL progression due to its weak binding for BAFF. To expand the therapeutic utility of sBCMA-Fc, using a protein engineering approach, we generated an affinity enhanced mutant sBCMA-Fc fusion molecule (sBCMA-Fc V3) with 4-folds and 500-folds enhancement in binding to APRIL and BAFF respectively. The sBCMA-Fc V3 clone significantly enhanced antitumor activity against both MM and DLBCL. Importantly, sBCMA-Fc V3 was proven to be a viable clinical candidate by showing adequate toxicity profile and on-target mechanism of action in nonhuman primates.

**SUMMARY:** This study demonstrates the dichotomous function of APRIL and BAFF in MM and DLBCL, that can be safely targeted by an engineered fusion protein designed to trap APRIL and BAFF with ultra-high binding affinity.

## INTRODUCTION

B cell malignancies, in particular, multiple myeloma (MM) and diffuse large B Cell Lymphoma (DLBCL) represent some of the most common hematological cancers worldwide (Chim et al., 2018; Durer et al., 2020; Swerdlow et al., 2016; Weber and Schmitz, 2022). While the use of treatment regimen combining chemo-cytotoxic drugs and targeted therapies has significantly improved overall survival, patients suffering from relapse/refractory diseases following standard-of-care (SOC) treatment still face a poor outcome (Caimi et al., 2021; Lonial et al., 2020; Raje et al., 2019). Recent approval of Chimeric Antigen Receptor (CAR)-T cells has provided a more effective treatment for patients who failed to respond to conventional therapies with some patients, achieving complete remission. Nonetheless, restrictive patient eligibility, immune-related adverse toxicities and treatment relapse are challenges remain to overcome. Therefore, safe and effective targeted therapies for patients who exhaust currently available treatment options is still a need.

Two ligands of the Tumor Necrosis Factor Superfamily (TNFSF) known as A Proliferation-Inducing Ligand (APRIL, TNFSF13) and B Lymphocyte Stimulator (BAFF, BLys and TNFSF13B) have been well documented as critical regulators of B Cell maturation and differentiation (Bolkun et al., 2014; Chiu et al., 2007). APRIL and BAFF facilitate their diverse functions on B lymphocytes through binding to three TNF receptors TACI (Transmembrane Activator and Ca^2+^ Modulator Interactor, TNFRSF13C), B Cell Maturation Antigen (BCMA, TNFRSF17) and B Cell Activating Factor Receptor (BAFF-R, TNFRSF13C) with different affinities (Wu et al., 2000; Yu et al., 2000). TACI and BAFF-R are primarily expressed on mature B Cells while BCMA expression is almost exclusively confined to plasma cells (O’Connor et al., 2004). To add another layer of complexity to this sophisticated molecular interplay, the binding affinity of APRIL and BAFF towards TACI, BCMA and BAFF-R differ so that each receptor-ligand interaction promotes unique biological outcomes. APRIL binds to BCMA with 1000-fold higher affinity compared to BAFF, whereas the reverse is observed with BAFF-TACI and BAFF-BAFF-R interactions (Marsters et al., 2000; Schuepbach-Mallepell et al., 2015; Yu et al., 2000). This distinctive receptor-ligand binding relationship further supports the differentiated roles of APRIL and BAFF in B Cell biology: APRIL-BCMA signaling is an exclusive regulator of plasma cells and BAFF-TACI/BAFF-R activation is required for the maturation of B lymphocytes (Moreaux et al., 2007; Pelletier et al., 2003). Pathologically, aberrant expression of APRIL and BAFF support disease progression and is associated with a poor treatment outcome in MM and DLBCL. Persistent APRIL and BAFF activation promotes survival advantages in MM and DLBCL, facilitating disease progression and treatment resistance (Kuo et al., 2008; Moreaux et al., 2004). Similar to the dichotomous relationship between APRIL and BAFF in normal B Cell development, APRIL-BCMA signaling play a primary role in MM pathology, while persistent activation of BAFF-TACI/BAFF-R is critical for promoting malignant B Cell growth and survival (Pham et al., 2011). Therefore, neutralizing APRIL and BAFF in B Cell malignancies may offer a new treatment option for MM and DLBCL.

In this study, we first investigated the therapeutic strategy of using soluble BCMA as a ligand trap for blocking APRIL mediated signaling in MM. While this approach has been explored with monoclonal antibodies to April (Tai et al., 2016), clinical testing did not demonstrate sufficient efficacy, most probably due to the high natural binding affinity between April and BCMA. Using a yeast surface display-based engineering approach, we developed an affinity-enhanced, soluble BCMA based mutant Fc fusion protein which effectively traps both APRIL and BAFF with enhanced binding affinities. This high affinity fusion protein showed superior blockade of both APRIL/BCMA signaling and BAFF-TACI/BAFF-R signaling in MM and DLBCL models, while demonstrating little toxicity and an on-target mechanism of action in nonhuman primate studies.

## RESULTS

### BCMA Signaling Activation is Required for Multiple Myeloma Progression and Promotes Protein Translation Machinery

Since the signaling interaction between APRIL and BCMA has being previously reported in MM, we further explored the biological significance of BCMA signaling in MM using genetic and biochemical approaches. Consistent with previous reports, BCMA mRNA was found to be uniquely elevated on MM and B Cell Lymphoma, and not detectable in cancer cells other than those of a B Cell origin (Fig. 1A, Sup. Fig. 1A). In addition, higher serum levels of APRIL (Fig. 1B left panel) and BAFF (Fig. 1B right panel) were also found in MM patients, which is consistent with previous reports (Bolkun et al., 2014; Fragioudaki et al., 2012).

**Figure 1:**
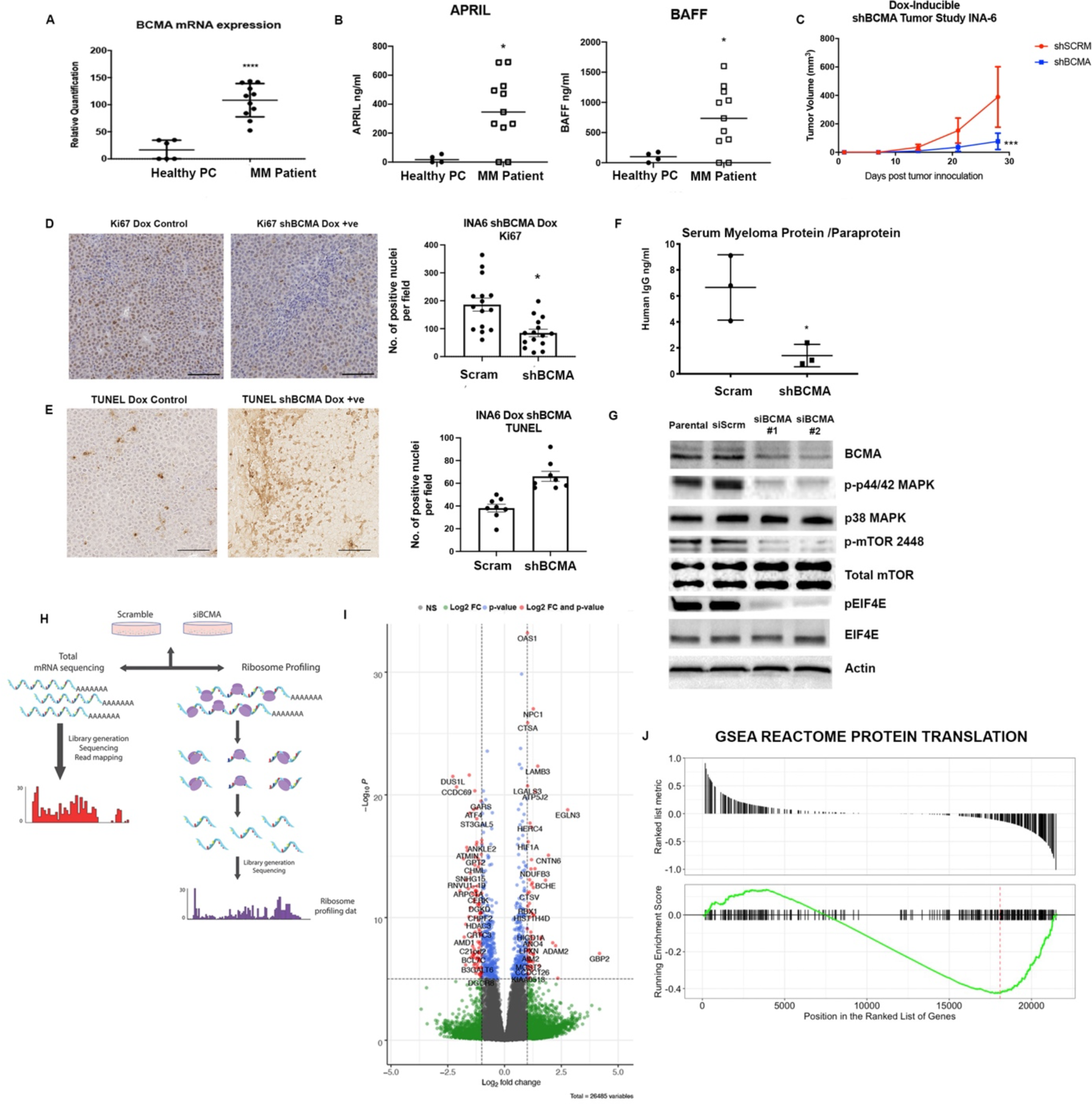
BCMA Signaling Activation is Required for Multiple Myeloma Progression and Promotes Protein Translation Machinery. **(A)** BCMA mRNA transcript validated in patient myeloma cells (N=11) and plasma cells harvested from healthy donors (N=6). (**B, left panel**) Serum APRIL level detected in MM patients (N=11) compared to healthy donors (N=4) through ELISA. (**B, right panel**) Serum BAFF level detected in MM patients (N=11) compared to healthy donors (N=4) through ELISA. **(C)** Subcutaneous tumor growth of INA-6 MM cells with stably transfected doxycycline-inducible BCMA KO shRNA (N=8) or scramble shRNA (N=7) in 6 weeks old female NOD-SCID Gamma (NSG) mice. **(D)** Representative images **(left)** and quantitation **(right)**, presented as the average number of positive nuclei per image field of Ki67 positive cells, in tumors of doxycycline-inducible shSCRM and shBCMA analyzed by IHC staining. Scale bar 50μm. **(E)** Representative images **(left)** and quantitation **(right)**, presented as the average number of positive nuclei per image field of TUNEL positive cells, in tumors of doxycycline-inducible shSCRM and shBCMA analyzed by IHC staining. Scale bar 50μm. **(F)** Total Human M protein (Paraprotein) detected in the serum of NSG mice inoculated with doxycycline-inducible shSCRM and shBCMA human INA-6 MM tumor cells harvested terminally (N=3). **(G)** Western blotting analysis of changes in protein expression associated with protein translation upon genetic knockdown using siBCMA. **(H)** Schematic illustration of ribosome profiling workflow. **(I)** Volcano plot analysis of global changes in protein translation upon loss of BCMA expression in MM. Not significant (gray), significant changes in Log_2_FC value (green), significant changes in p-value (red) and significant changes in both p-value and log_2_FC (red) are presented according to color coding. Left of the volcano plots shows reduction in target expression upon loss of BCMA, the right side shows negative correlation upon loss of BCMA. **(J)** GSEA REACTOME enrichment analysis showing enriched signatures in associated with protein translation. Statistical analysis was conducted using T test and one way ANOVA for comparing between treatment groups. Repeated ANOVA used for changes in tumor growth over time. P value *=<0.05, **=<0.01. ***=<0.001.

The dependency of MM proliferation on BCMA signaling was tested by introducing a Tet-off doxycycline-controlled BCMA stable knockdown system (dox shBCMA) into MM cell lines INA-6 and MM1.R (Sup. Fig. 1B). INA-6 MM cell has chromosomal translocation at t(11;14) with *NRAS* and *TP53* mutations and is dependent on IL-6 for growth and survival (Burger et al., 2001; Keats et al., 2007). MM1.R has chromosomal translocations at t(14;16) and t(8;14) with *KRAS* and *TRAF3* mutations (Greenstein et al., 2003). Genetic inhibition of BCMA with dox inducible shBCMA led to a significant decrease in established MM tumor size in both models (Fig. 1C. Sup. Fig 1C, D). Further demonstrating the important role of BCMA signaling despite the cytogenetic heterogeneity that occurs within MM. Tumors harvested from both models showed decreased proliferation (Fig. 1D, Sup. Fig. 2A, B) and increased apoptosis (Fig. 1E, Sup. Fig. 2C, D) upon genetic inhibition of BCMA. Furthermore, the level of human myeloma protein (paraprotein) secreted by human MM tumor cells was significantly reduced in the mice bearing shBCMA MM tumors, also indicative of decreased tumor burden (Fig. 1F) (Collier and Jackson, 1953).

**Figure 2:**
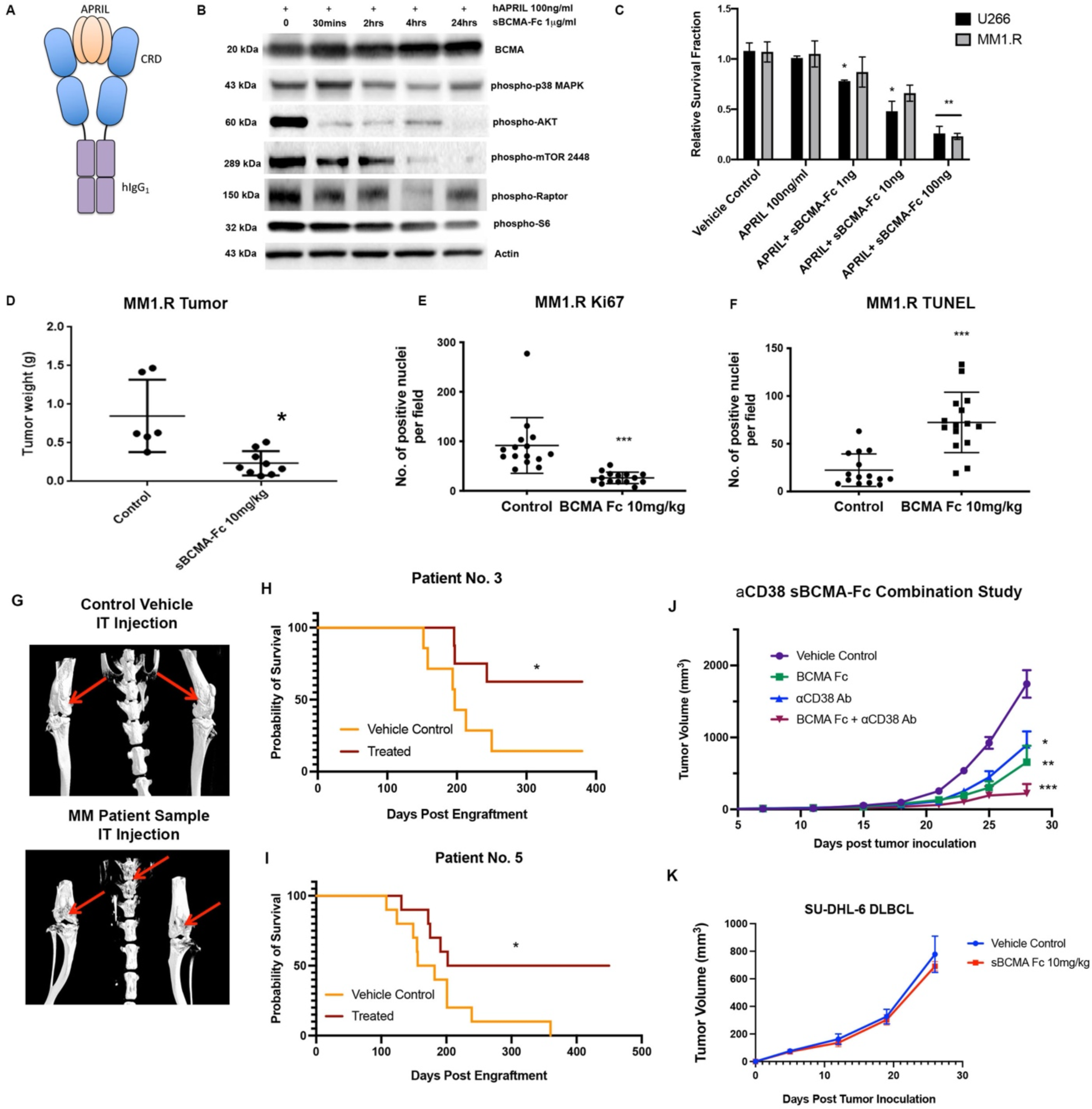
Wildtype sBCMA Decoy Receptor inhibits Multiple Myeloma Growth Through APRIL/BCMA Signaling but Lacks Efficacy in BAFF Driven DLBCL Model. **(A)** Schematic illustrations of recombinant human sBCMA-Fc binding to human ARPIL. **(B)** Analysis of BCMA downstream protein expression in U266 MM cells upon sBCMA-Fc treatment at multiple time point. **(C)** sBCMA-Fc dose-dependent, cytotoxicity assay validating the *in vitro* cell survival in the presence of increasing doses of sBCMA-Fc and hAPRIL (100ng/ml) in U266 and MM1.R MM cells. Cells were maintained in a low 3% FCS to reduce possible growth stimulation mediated through other growth factors present in FCS. Each sample was performed in triplicates. **(D)** Terminal tumor weight of mice inoculated with MM1.R MM tumors and treated with vehicle control or 10mg/kg of sBCMA-Fc. **(E)** Quantification of Ki67 staining in MM1.R MM tumors and treated with vehicle control or 10mg/kg of sBCMA-Fc. **(F)** Quantification of TUNEL staining in MM1.R MM tumors and treated with vehicle control or 10mg/kg of sBCMA-Fc. **(G)** Representative CT scans of mice tibias, femurs and vertebrates inoculated with control (top) or MM PDX tumor cells (bottom). Osteolytic bone degradation was observed in MM PDX injected animal (bottom image) but not in the control injected animals (top image). **(H)** Kaplan Meier survival analysis of animals engrafted with MM cells from patient no. 3 showing prolonged overall survival in sBCMA-Fc treated group (N=8) compared to the vehicle control (N=7). Medium survival (in days) shown below. **(I)** Kaplan Meier survival analysis of animals engrafted with MM cells from patient no. 5 showing prolonged overall survival in sBCMA-Fc treated group (N=10) compared to vehicle control (N=10). Medium survival (in days) shown below. **(J)** Subcutaneous tumor growth of MM1.R MM tumors in 6 weeks old female NSG mice dosed with sBCMA-Fc 10 mg/kg every 48 hours (N=7), *α*CD38 10mg/kg weekly (N=7), sBCMA-Fc and *α*CD38 combination (N=8) and vehicle control (N=8). **(K)** Subcutaneous tumor growth of SU-DHL-6 DLBCL tumors in mice dosed with vehicle control or sBCMA-Fc 10 mg/kg every 48 hours (N=5). Statistical analysis was conducted using T test and one way ANOVA for comparing between treatment groups. Repeated ANOVA used for changes in tumor growth over time. P value *=<0.05, **=<0.01. ***=<0.001.

Upon analyzing downstream biological changes associated with BCMA signaling, we found robust changes in the key regulators of protein translation and synthesis such as EIF family member eIF4E, suggesting a potential link between BCMA signaling and protein translation that has not been described previously (Fig 1G, Sup. Fig. 3A). To investigate whether BCMA acts as a regulator of protein translation globally or selectively, we performed ribosome profiling on U266 MM cells with genetic BCMA knockdown using siRNA. U266 harbors some of the most frequently found genetic alternations in MM patients and is a reasonable representative of MM tumor cells. Ribosome profiling is a specialized analysis method that provides a quantitative measure of gene-specific translation efficiency. Technically, mRNA fragments protected by ribosomes were isolated and sequenced, and in the same population of cells total RNA abundance was quantified by RNA-Seq for normalization (Fig. 1H) (Ingolia et al., 2011). We performed extensive quality control analysis of the ribosome profiling data to ensure accurate ribosome bound RNA readouts (Sup Fig. 3B to F). First, we analyzed total mRNA obtained from RNA-Seq data which serves as a reference for quantitating ribosome bound mRNA associated changes in translation efficiency (Sup. Fig 4A, B). Calculation of translation efficiency was conducted by analyzing the ribosome profiling data obtained from ribosome bound mRNA and normalized against total mRNA (Fig. 1I) (Cenik et al., 2015). Most notably, we found enriched gene signatures associated with initiation and elongation of protein translation (Fig. 1J, Sup. Fig. 5A, C) Furthermore, pathways that are responsible for the synthesis of both large and small ribosome subunits were also affected in BCMA inhibited MM cells, suggesting a potential role of BCMA signaling in ribosome biogenesis (Sup. Fig. 5D). This finding is consistent with our reverse phase protein array (RPPA), which shows a decreased protein translation signature upon loss of BCMA (Sup. Fig. 5B). Collectively, these data suggest a previously unknown role of BCMA signaling as a regulator of protein translation machinery in MM growth.

**Figure 3:**
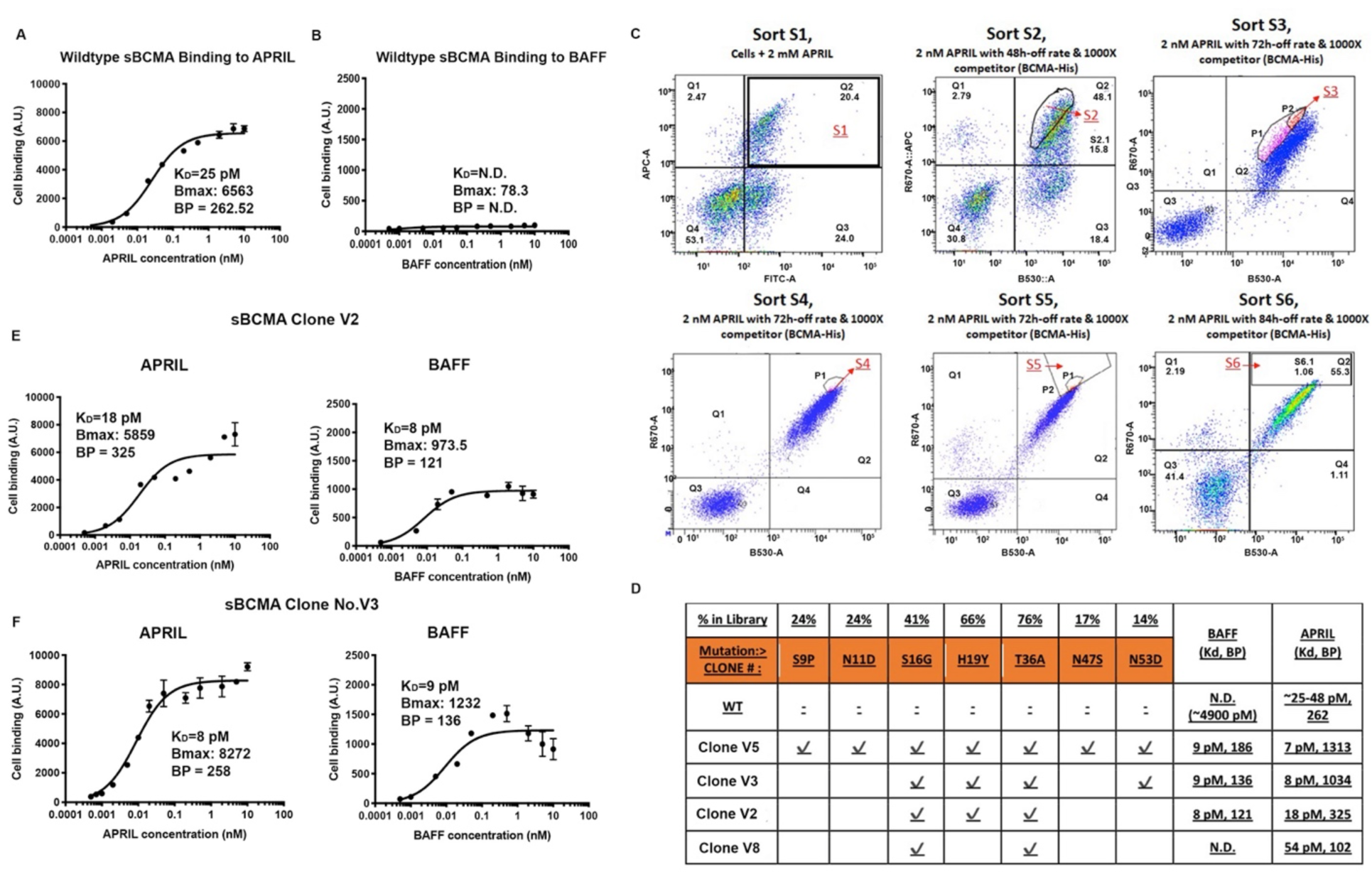
Engineering High-Affinity Decoy Receptor Fusion Protein against APRIL and BAFF. **(A)** Flow cytometry based binding curve showing yeast-displayed wildtype sBCMA binding to increasing concentration of ARPIL. Calculated K_D_, B_max_ and BP is also shown. A.U., arbitrary units. **(B)** Flow cytometry based binding curve showing yeast-displayed wildtype sBCMA binding to increasing concentration of BAFF. Calculated K_D_, B_max_ and BP is shown. **(C)** Overlaid flow cytometry dot plots representing sorting strategies of yeast-displayed sBCMA library binding to 2nM APRIL in 6 consecutive sorts with 48-84 hours off rate binding and 1000x competitor. Gated populations are collected from each sort and propagated for the next sorting round. **(D)** Binding affinities to APRIL and BAFF of wildtype sBCMA and selected mutant sBCMA clones. Conserved amino acid mutations were identified in mutant clones and the frequency of occurring mutation is also listed. **(E)** Binding curve of high-affinity mutant sBCMA clone V2 binding to increasing concentration of ARPIL (left) and BAFF (right). Calculated K_D_, B_max_ and BP is shown. **(F)** Binding curve of high-affinity mutant sBCMA clone V3 binding to increasing concentration of ARPIL (left) and BAFF (right). Calculated K_D_, B_max_ and BP is shown.

**Figure 4:**
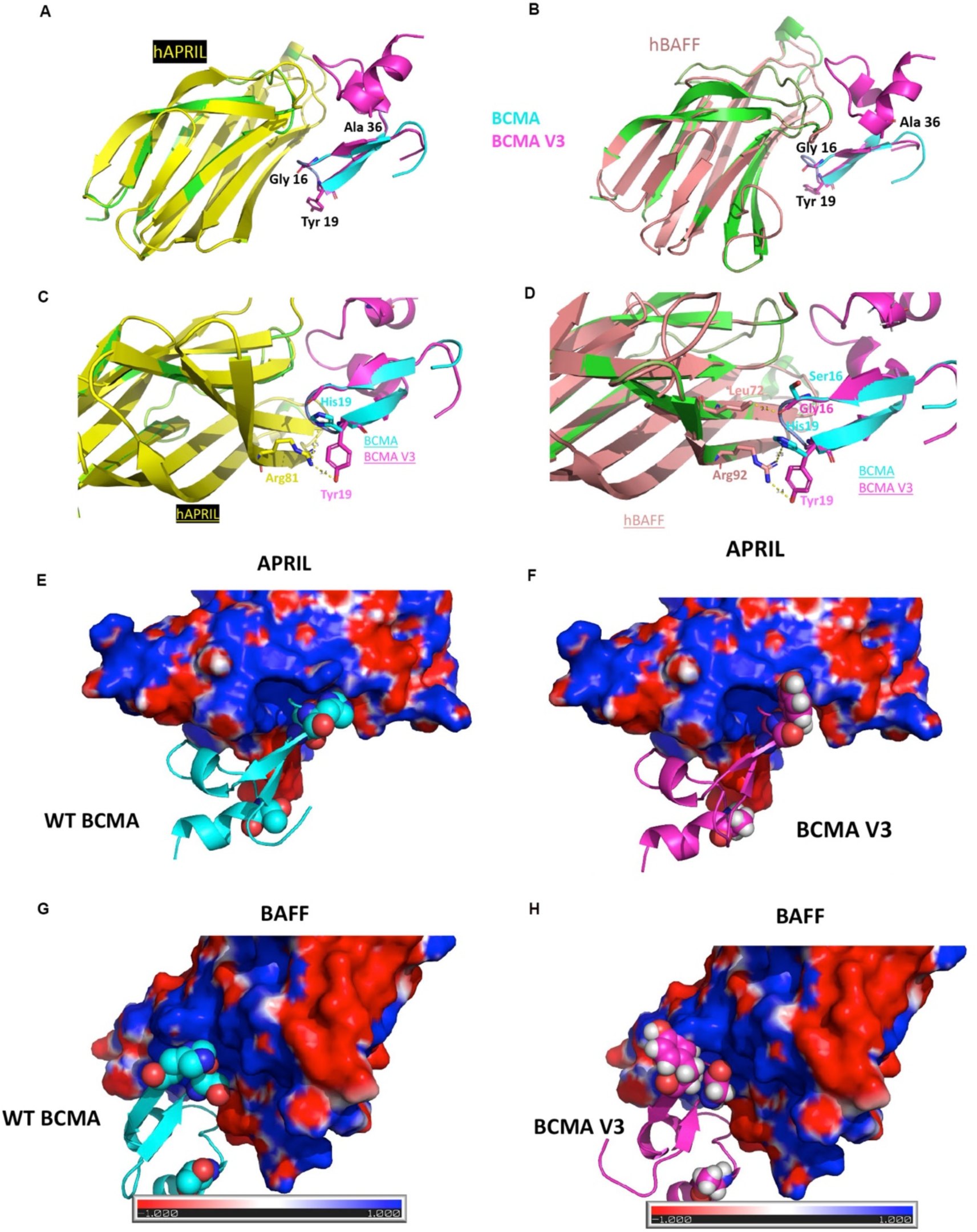
**(A)** Computational modeling of sBCMA V3 mutant clone (magenta) in co-complex with hAPRIL (yellow) overlaying on PDB structure 1XU2 consisting of human BCMA (cyan) in co-complex mAPRIL (Green). Mutation sites Gly16, Tyr19 and Ala36 were showed in stick. **(B)** Computational modeling of sBCMA V3 mutant clone (magenta) in co-complex with hBAFF (salmon) overlaying on PDB structure 1XU2 consisting of human BCMA (cyan) in co-complex mAPRIL (Green). Mutation sites Gly16, Tyr19 and Ala36 were showed in stick. **(C)** Predicted binding interaction between hAPRIL (yellow) and sBCMA V3 (cyan) showing Arg81 on hAPRIL interact with H19Y on sBCMA V3. (D) Predicted binding interaction between hBAFF (salmon) and sBCMA V3 (cyan) showing Arg93 on hBAFF interact with H19Y on sBCMA V3. Comparison of surface complementarity of wildtype BCMA with hAPRIL **(E)** and hBAFF **(G)**, sBCMA V3 with hAPRIL **(F)** and hBAFF **(H).** Electrostatic surface of hAPRIL and hBAFF are shown. Red indicates negative electrostatic potential; blue indicates positive electrostatic potential and gray indicates hydrophobic regions. The mutant sites of BCMA were represented in ball structure.

**Figure 5:**
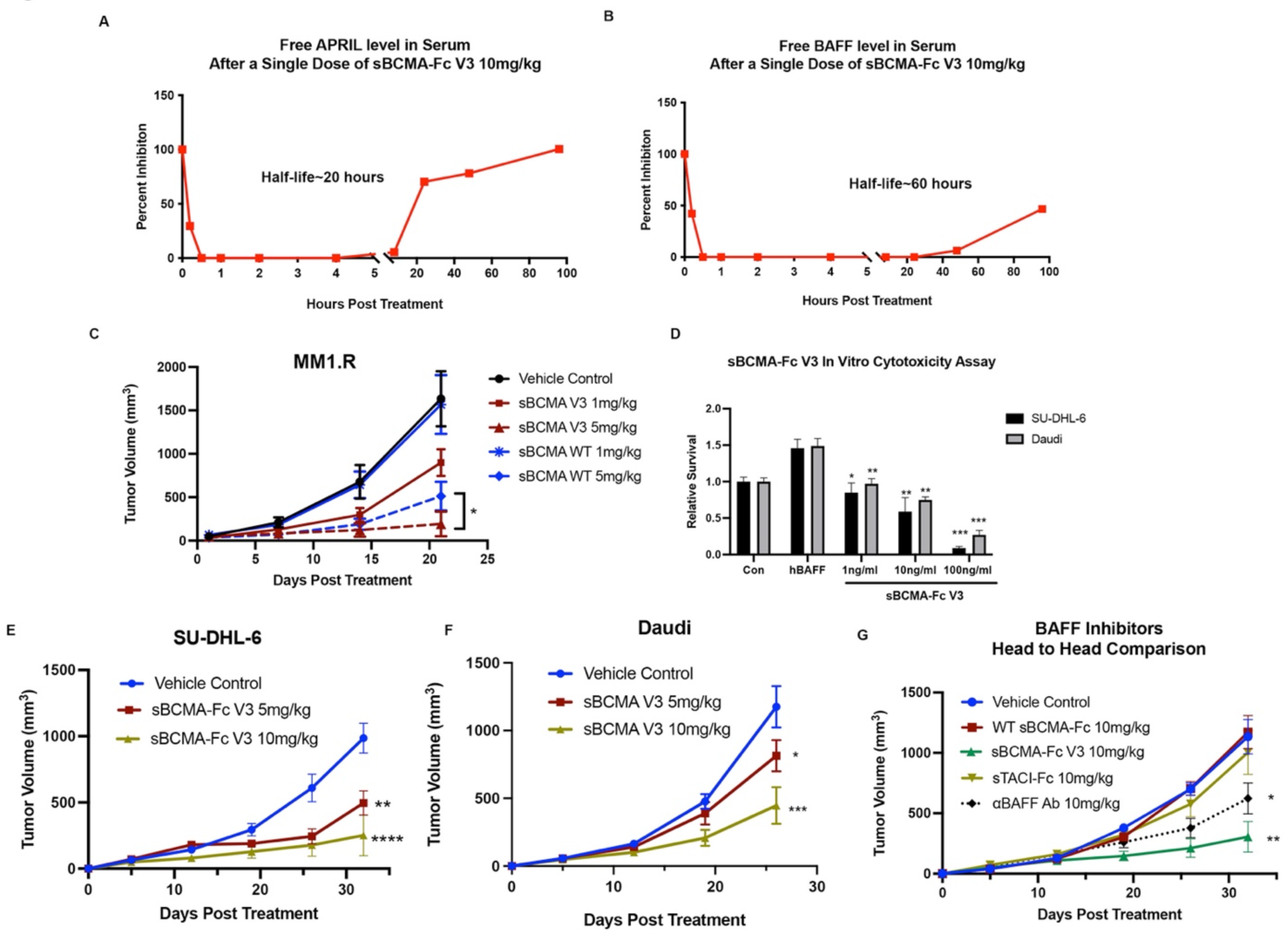
**(A)** Serum level of APRIL in mouse serum after a single dose of sBCMA-Fc V3 at 10mg/kg. Each data point represents duplicate repeats collected at each time point. **(B)** Serum level of BAFF in mouse serum after a single dose of sBCMA-Fc V3 at 10mg/kg. Each data point represents duplicated repeats collected at each time point. **(C)** Subcutaneous tumor growth of MM1.R MM tumors in 6 weeks old female NSG mice dosed with wildtype sBCMA-Fc at 1 and 5 mg/kg every 48 hours (N=5), sBCMA-Fc V3 at 1 and 5 mg/kg every 48 hours (N=5) and vehicle control (N=5). **(D)** sBCMA-Fc V3 dose-dependent, cytotoxicity assay validating the *in vitro* cell survival in the presence of increasing doses of sBCMA-Fc V3 and hBAFF (100ng/ml) in SU-DHL-6 and Daudi DLBCL cells. Cells were maintained in a low 3% FCS to reduce possible growth stimulation mediated through other growth factors present in FCS. Each sample was performed in triplicates. **(D)** Subcutaneous tumor growth of SU-DHL-6 DLBCL tumors in 6 weeks old female *NOD-scid* mice dosed with 5 mg/kg, 10 mg/kg sBCMA-Fc V3 every 48 hours (N=5) and vehicle control (N=5). **(F)** Subcutaneous tumor growth of Daudi DLBCL tumors in 6 weeks old female *NOD-scid* mice dosed with 5 mg/kg, 10 mg/kg sBCMA-Fc V3 every 48 hours (N=5) and vehicle control (N=5). **(G)** *In vivo* head-to-head comparison of anti-tumor efficacies in subcutaneous SU-DHL-6 DLBCL tumors treated with 10 mg/kg wildtype sBCMA-Fc every 48 hours (N=5), 10 mg/kg sBCMA-Fc V3 every 48 hours (N=5), 10 mg/kg sTACI-Fc every 48 hours (N=5) and 10 mg/kg *α*BAFF Antibody twice a week (N=5). Subcutaneous tumor growth was monitored throughout the study. Statistical analysis was conducted using T test and one way ANOVA for comparing between treatment groups. Repeated ANOVA used for changes in tumor growth over time. P value *=<0.05, **=<0.01. ***=<0.001.

### Wild-type sBCMA Decoy Receptor inhibits Multiple Myeloma Growth Through APRIL/BCMA Signaling but Lacks Efficacy in BAFF Driven DLBCL Models

Since gain of functional mutations in BCMA are rarely reported, BCMA signaling activity is almost entirely regulated by its ligands APRIL and BAFF. Therefore, we investigated a ligand blocking approach, utilizing a soluble decoy receptor comprised of the BCMA extracellular domain (ECD) fused to the human IgG1 Fc domain (sBCMA-Fc) to trap and neutralize APRIL and BAFF (Fig. 2A).

Treatment with sBCMA-Fc decreased downstream protein expression of similar canonical signaling regulated by the BCMA that we found through genetic inhibition of BCMA (Fig. 2B). To determine the inhibitory effect of sBCMA-Fc on the viability of MM cells, we performed *in vitro* cytotoxicity assays using U266 and MM1.R cells. A concentration dependent decrease in MM cell viability was observed when cells treated with sBCMA-Fc and cultured in exogenous APRIL with reduced serum conditions, supporting the importance for APRIL as a growth stimulus in MM cells (Fig 2C). Thus, sBCMA-Fc effectively neutralized ligand mediated activation of BCMA signaling pathway *in vitro,* resulting in decreased MM cell growth.

We next examined the efficacy of sBCMA-Fc *in vivo.* Treatment with sBCMA-Fc resulted in a significant tumor reduction in both MM1.R (Fig. 2D) and INA-6 MM tumors (Sup. Fig. 6A) and was associated with decreased tumor cell proliferation (Fig. 2E, Sup. Fig. 6B) and increased apoptosis (Fig. 2F, Sup. Fig. 6C). Collectively, these findings provide supporting evidence that sBCMA-Fc is efficacious as a monotherapy in xenograft models of MM. To further validate the therapeutic potential of sBCMA-Fc in human MM specimens, we established human PDXs MM models by engrafting freshly isolated human MM cells into the tibia of an immunodeficient murine host. We screened eight male and three female patients, of whom six were untreated and five were treated. We found that the majority of patients had IgG kappa (IgG*κ*) or IgG lambda (IgG*λ*) myeloma type (Sup. Fig. 7A). MM cells from patient no. three and five (both IgG*κ* myeloma) were successfully engrafted and propagated for *in vivo* testing. After inoculation of PDX derived MM cells, we confirmed engraftment by assessing the increase in levels of human M protein (IgG*κ*) in mouse serum (Sup. Fig. 4B) and then initiated sBCMA-Fc treatment. CT scans of animals with successful engraftment showed macroscopic osteolytic lesions that are consistent with osteopenia found in MM patients (Fig. 2G). Engrafted mice were dosed with 10 mg/kg doses of sBCMA-Fc every 48 hours for 28 days and tumor growth was evaluated by detecting human IgG*κ* in blood until mice reached a terminal endpoint. Treatment with sBCMA-Fc led to a significant reduction in tumor progression based on IgG*κ* signals and prolonged overall survival in both PDX lines that were successfully propagated, demonstrating the ligand dependency of BCMA signaling in human MM cells (Fig. 2H, I, Sup. Fig. 7C, D). We also evaluated the therapeutic relevance of sBCMA-Fc in combination with other MM targeted therapies such as *α*CD38 antibodies. The combination of sBCMA-Fc with a CD38 therapeutic antibody (*α*CD38) showed superior antitumor activity compared to either monotherapy alone (Fig. 2J, Sup. Fig. 8A). There were no differences in bodyweight among the treatment groups (Sup Fig. 8B).

**Figure 6:**
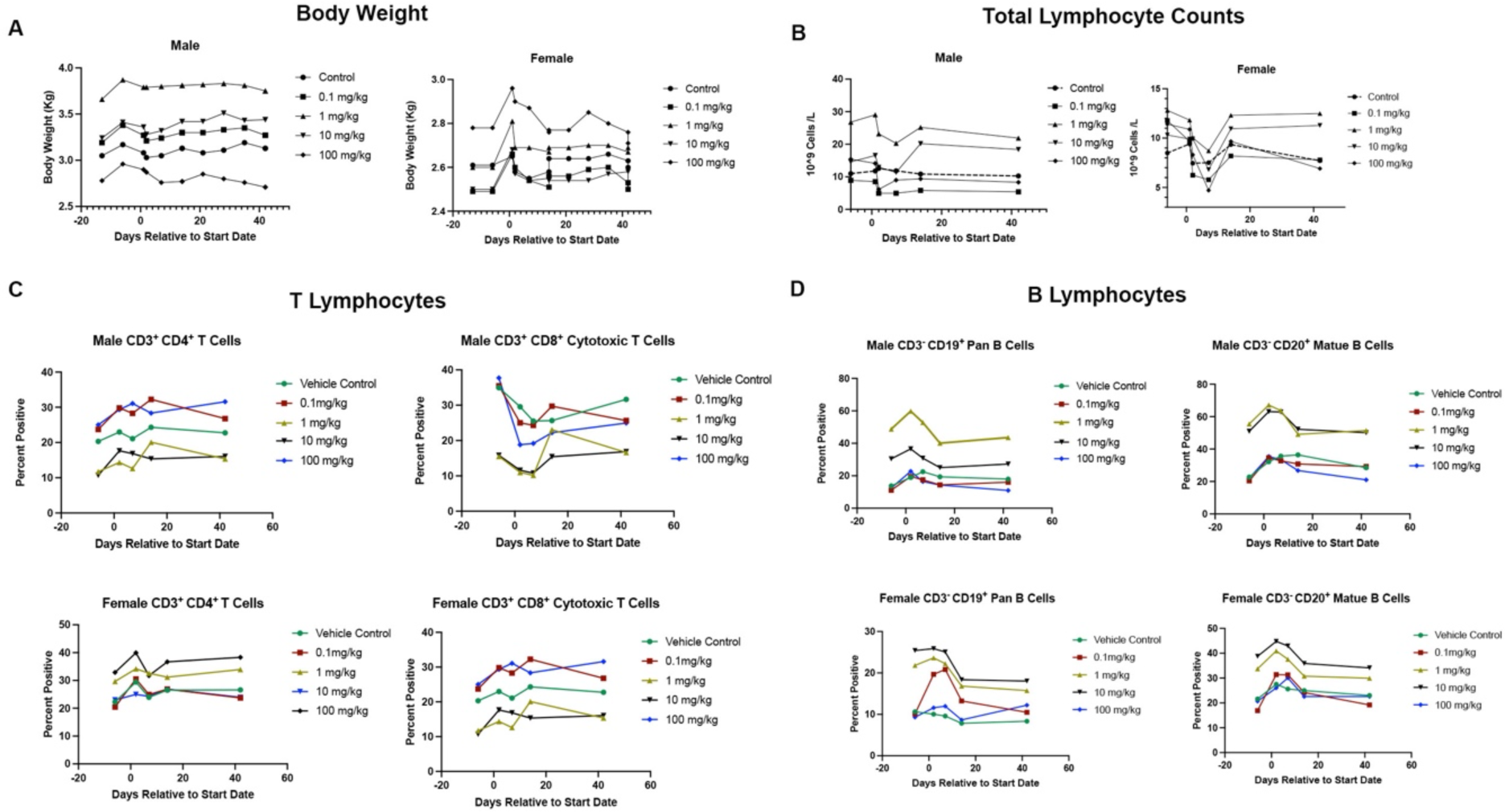
**(A)** Changes in total body weight in kilograms of male (left) and female (right) cynomolgus monkey test subjects throughout the experiment and observation period. (B) Total lymphocyte counts expressed as 10^9 cells/Liter in the blood of male (left) and female (right) cynomolgus monkey test subjects. **(C)** Counts of CD3^+^CD4^+^ T lymphocytes and CD3^+^CD8^+^ cytotoxic T lymphocytes by Flow Cytometry in the blood of male (top) and female (bottom) cynomolgus monkey test subjects pre and post dosing as well as during observation period. **(D)** Counts of CD3^-^ CD19^+^ Pan B lymphocytes and CD3^-^CD20^+^ Mature B lymphocytes by Flow Cytometry in the blood of male (top) and female (bottom) cynomolgus monkey test subjects pre and post dosing as well as during observation period. Statistical analysis not applicable.

Based on published studies, BAFF is also capable of binding to BCMA receptor at a lower affinity (Bossen and Schneider, 2006), we wanted to investigate the therapeutic efficacy of sBCMA-Fc in B Cell malignancy models such as DLBCL, which in contrast to MM, is BAFF dependent (Fu et al., 2021; Lyu et al., 2010). BAFF sensitive SU-DHL-6 and Daudi DLBCL cells were treated with different concentration of sBCMA-Fc to assess *in vitro* cytotoxicity. Interestingly, we only observed cytotoxicity in DLBCL cell lines when they were treated with highest dose of 100ng/ml of sBCMA-Fc (Sup. Fig. 8C, D). To determine the efficacy of sBCMA-FC *in vivo*, mouse bearing subcutaneous SU-DHL-6 tumors were treated with sBCMA-Fc at 10mg/kg every two days, which is the efficacious dosing schedule used in the MM tumor models. In contrast to MM, treatment with sBCMA-Fc at 10mg/kg didn’t provide any significant antitumor benefit in BAFF sensitive SU-DHL-6 lymphomas (Fig. 2K). Since MM is primarily an APRIL driven disease and DLBCL is BAFF dependent, we hypothesized that sBCMA-Fc lacks sufficient binding affinity to BAFF inhibit BAFF mediated signaling.

### Engineering High-Affinity Decoy Receptor Fusion Protein Against APRIL and BAFF

To directly determine whether the lack of antitumor activity demonstrated by sBCMA-Fc against DLBCL is primarily due to weak binding affinity to BAFF, we evaluated the binding kinetics of sBCMA to hAPRIL and hBAFF. While the K_D_ between sBCMA and hAPRIL has been reported to be between 25-48 pM (Hymowitz et al., 2005; Yu et al., 2000), the K_D_ between sBCMA and hBAFF was weak and was experimentally estimated to be approximately 4900 pM (Fig. 3A, B). This finding supported our hypothesis that the low binding between sBCMA-Fc and BAFF is likely to be a contributing factor to poor therapeutic efficacy in BAFF driven DLBCL models.

To address this shortcoming, we engineered a sBCMA mutant with the capability of neutralizing both APRIL and BAFF at high affinity. Using low-fidelity Taq polymerase-based error-prone PCR as previously described (Miao et al., 2021), nucleotide mutations were randomly generated into the extracellular domain of BCMA gene from amino acid 1 (Methionine) to 54 (Alanine). Purified cDNA Products from the PCR reactions were electroporated into EBY100 yeast cells for library assembly, the library size was estimated to be 2x10^8^ by dilution plating. The resulting library displayed on yeast surface was sorted by fluorescence activated cell sorting (FACS) to isolate clones with desired binding characteristics. We carried out initial affinity screening using hAPRIL because it was previously report that the BCMA binding site to ARPIL and BAFF shares high homology. Therefore, clones identified in this screening process will likely have high binding affinity to both APRIL and BAFF (Liu et al., 2003). Six rounds of FACS sorting were performed sequentially. Equilibrium sorting in Round 1 was performed at room temperature with 2nM of hAPRIL. Subsequent kinetic off-rate sorts were conducted in the presence of 2 nM APRIL for 3 hours at room temperature, washed and were incubated with approximately 1000x molar excess of sBCMA to render unbinding events irreversible. The incubation time for each consecutive dissociation step increases from 48 to 84 hours and 1-3% clones with the highest binding to APRIL were enriched from each sort and propagated for the subsequent sorting round (Fig. 3C).

The top 118 clones selected from sort rounds 3 to 6 were sequenced and analyzed for consensus mutations (Sup. Tables 1 to 4). Overall, 7 consensus mutations with high frequencies were identified (Fig. 3D). These mutations are: 1) position 9 Serine to Proline (S9P), 2) position 11 Asparagine to Aspartic Acid (N11D), 3) position 16 Serine to Glycine (S16G), 4) position 19 Histidine to Tyrosine (H19Y), 5) position 36 Tyrosine to Alanine (T36A), 6) position Asparagine to Serine (N47S) and 7) Asparagine to Aspartic Acid (N53D). Based on binding analysis of conserved mutant clones to APRIL and BAFF, we identified 4 clones with high binding affinity based on K_D_ and B_max_ measured for each ligand of interest (Fig. 3D). Interestingly, while APRIL was used for both equilibrium and kinetics off-rate sorting, we observed a dramatic improvement in the binding affinities between candidate mutant clones and BAFF. sBCMA V3 carrying mutations S16G, H19Y, T36A and N53D was selected as our top candidate because it possessed fewer mutations while still retained high binding affinity towards APRIL and BAFF (Fig. 3E, F).

### Computational Analysis Identifying Structural Evidence for Affinity Enhancement

To determine the changes associated with both structural and biophysical properties of our top candidate clone sBCMA V3 that result in its affinity enhancement, we performed computational model simulations of sBCMA-Fc V3 in co-complex with APRIL and BAFF. First, we performed sequence alignment and subsequent modeling based on the wild-type hBCMA and hAPRIL complex structures obtained from Protein Data Bank PDB No. 1XU2 (Hymowitz et al., 2005). This structure served as the backbone for mapping amino acid mutations. Additional structural information on hAPRIL and hBAFF was derived from PDB No. 4ZCH which reported the single-chain human APRIL-BAFF-BAFF heterotrimer structure (Schuepbach-Mallepell et al., 2015). Sequence alignment between hAPRIL, hBAFF and mAPRIL are shown in Sup. Fig. 9A. As expected, the sequence identity between hAPRIL and mAPRIL (1XU2) shared a high identity of 86%, compared to hBAFF and mAPRIL which only had 40% sequence homology (Sup. Fig. 9B). We also performed structural alignment comparing hAPRIL, hBAFF and mAPRIL (1XU2). Structural alignment between hAPRIL and mAPRIL revealed high structural homology with a Root-Mean-Square-Deviation (RMSD) score of 0.893Å. Interestingly, while hBAFF and mAPRIL were not highly conserved in their primary sequence, they shared high similarity on tertiary structure with RMSD score of 1.248Å (Sup. Fig. 9C). This finding is consistent with a previous report that hAPRIL and hBAFF share a similar binding site at the DxL motif of BCMA (Gordon et al., 2010). Based on information we obtained from structural alignment, we constructed models of BCMA/hAPRIL (Fig. 4A) and BCMA/hBAFF (Fig. 4B) co-complexes using Prime from Schrödinger Suites. Currently, only partial BCMA structure between amino acid residues 8 to 43 have been reported and publicly available. This is because amino acid residue 1 to 7 forms the cleavable signal peptide which subsequently detached from the mature protein. Using multiple modeling software, we predict that residue 44 to 54 are likely to be a highly flexible and there is no reported electron density information available. Since predicting the biophysical properties of a flexible region is not possible, we were not able to calculate the affinity and stability attribution of the fourth mutation (N53D) on sBCMA V3 clone (Sup. Fig. 9D).

To analyze the structural and biophysical changes associated with first three amino acid mutations in sBCMA V3, we performed modeling using AlphaFold2, trRosetta, RoseTTAFold and RosettaRemodel (Fig. 4C, D). Using PDB structure 1XU2 as a reference, we found all modeling analysis with the exception of RoseTTAFold produced structural prediction with high confidence. RoseTTAFold had low structural simulation confidence and the predicted structure failed to be mapped onto 1XU2 (Sup. Fig. 9D). It is worth noting that out of the four mutations, mutations S16G, H19Y are located within the BCMA binding DxL motif and T36A, N53D are located outside of the binding motif. Mutation driven changes in binding affinity and overall protein stability between BCMA V3/APRIL or BAFF was calculated by the Residue Scanning Calculation module from Bioluminate, Schrodinger. Mutations S16G, H19Y and T36A individually or collectively led to improvement in the binding affinity, overall stability and thermodynamics of sBCMA V3/hAPRIL co-complex (Table 1). Similarly, these mutations contributed towards enhanced affinity, stability and thermodynamics of sBCMA V3/hBAFF co-complex (Table 2). Based on protein interaction analysis, the H19Y mutation is likely to be the most important contributor responsible for improved binding affinity to both hAPRIL and hBAFF. While residue 19 has been reported as a key residue associated with interaction between wild-type BCMA and BAFF (Bossen and Schneider, 2006), there has been no previous report of its role in APRIL binding. Here, we found amino acid substitution from Histidine to Tyrosine on BCMA residue no. 19 is a critical mutation responsible for enhanced binding to both APRIL and BAFF. Surface complementarity analysis between wild-type sBCMA, sBCMA V3, hAPRIL and hBAFF was also performed (Fig. 4E to H). Again, H19Y stood out as the key mutation which resulted in the improvement of surface complementarity between sBCMA V3/hAPRIL/hBAFF co-complexes (Table 3). In its co-complex form, H19Y mutation shortened the intramolecular distance from 3.7 Å to 2.1 Å between sBCMA V3 Tyr19 and Arg81 on hAPRIL. Similarly, the intramolecular distance between sBCMA V3 Tyr19 and Arg92 on hBAFF was reduced from 3.7 Å to 2.0 Å, corresponding to a tighter binding co-complex structure. Mutation S16G on sBCMA V3 also improved surface complementarity to both hAPRIL and hBAFF to a lesser extent compared to H19Y mutation but did not alter intramolecular distance. Mutation T36A did not affect these parameters because this mutation is away from the DxL binding motif on BCMA.

**Table 1:** Calculated Bind Affinity and Protein Stability (in kcal/mol for Each BCMA Mutation in Co-Complex with APRIL.

| Residue | Original | Mutated | d Affinity | d Stability (solvated) | Total dG Value Original/Mutated |
| --- | --- | --- | --- | --- | --- |
| R:16 | SER | ALA | -0.13 | -1.78 | N/A |
| R:19 | HIS | TYR | -3.95 | -3.59 | N/A |
| R:36 | THR | ALA | -0.22 | 4.91 | N/A |
| R:16<br>R:19<br>R:36 | SER<br>HIS<br>THR | ALA<br>TYR<br>ALA | -5.85 | -1.19 | -4.333/-33.041 |

**Table 2:** Calculated Bind Affinity and Protein Stability (in kcal/mol for Each BCMA Mutation in Co-Complex with BAFF.

| Residue | Original | Mutated | d Affinity | d Stability (solvated) | Total dG Value Original/Mutated |
| --- | --- | --- | --- | --- | --- |
| R:36 | THR | ALA | -0.14 | 4.98 | N/A |
| R:19 | HIS | TYR | -7.73 | -7.52 | N/A |
| R:16 | SER | GLY | 1.38 | 1.03 | N/A |
| R:16<br>R:19<br>R:36 | SER<br>HIS<br>THR | GLY<br>TYR<br>ALA | -3.55 | 2.8 | -17.462/-19.858 |

**Table 3:**
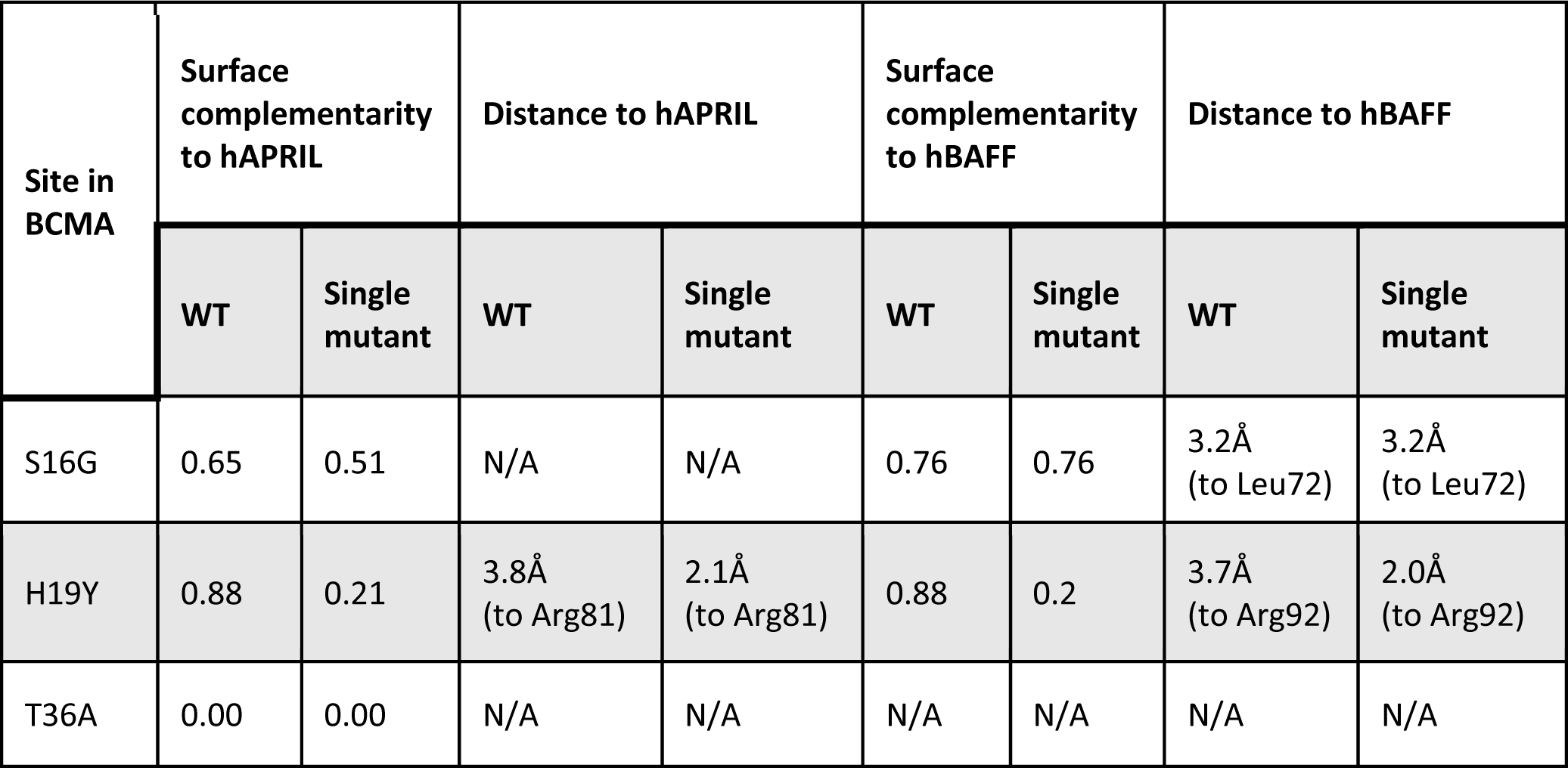
Calculated Surface Complementarity and Molecular Distance Between BCMA V3 and APRIL/BAFF.

| Site in BCMA | Surface complementarity to hAPRIL |  | Distance to hAPRIL |  | Surface complementarity to hBAFF |  | Distance to hBAFF |  |
| --- | --- | --- | --- | --- | --- | --- | --- | --- |
|  | WT | Single mutant | WT | Single mutant | WT | Single mutant | WT | Single mutant |
| S16G | 0.65 | 0.51 | N/A | N/A | 0.76 | 0.76 | 3.2Å<br>(to Leu72) | 3.2Å<br>(to Leu72) |
| H19Y | 0.88 | 0.21 | 3.8Å<br>(to Arg81) | 2.1Å<br>(to Arg81) | 0.88 | 0.2 | 3.7Å<br>(to Arg92) | 2.0Å<br>(to Arg92) |
| T36A | 0.00 | 0.00 | N/A | N/A | N/A | N/A | N/A | N/A |

### Affinity-Enhanced sBCMA-Fc V3 Promotes Antitumor Efficacies in MM and DLBCL Xenograft Models

To establish the therapeutic potency of affinity-enhanced mutant clone sBCMA V3 in models of Multiple Myeloma and B cell malignancies, we generated a sBMCA V3+hIgG1 fusion protein with the addition of a human IgG1 Fc domain (sBCMA-Fc V3) for improved stability and bioavailability. The pharmacokinetics of sBCMA-Fc V3 was validated in *NOD-scid* immunocompromised non-tumor bearing mice. A single dose of 10mg/kg sBCMA-Fc V3 was injected into non-tumor bearing animal completely suppressed the level of ARPIL and BAFF, respectively, within 30 minutes (Fig 5A, B). Over time, sBCMA-Fc V3 showed an inhibitory half-life of approximately 20 hours with APRIL suppression (Fig. 5A) and 60 hours with BAFF inhibition (Fig. 5B). This finding is consistent with the half-life of human decoy receptors in mice reported previously (Kariolis et al., 2017; Miao et al., 2021).

To evaluate whether the affinity enhanced sBCMA-Fc V3 increases the antitumor activity in MM models previously tested with wild-type sBMCA-Fc, we evaluated the effect of 1, 5 and 10 mg/kg of wild-type sBMCA-Fc or sBCMA-Fc V3 on MM tumor growth. A significant improvement in antitumor effects was observed for both concentrations of sBCMA-Fc V3 compared to the wild-type sBCMA-Fc treated at the same doses (Fig 5C). Interestingly, the antitumor affects are comparable when animals were treated with both molecules at a higher dose of 10 mg/kg, suggesting both treatments have reached plateau in achieving ligand/receptor inhibition (Sup. Fig. 10A. Since MM progression is primarily APRIL driven, the enhanced APRIL binding in sBCMA-Fc V3 treated groups resulted in more efficient APRIL neutralization at lower concentration.

Next, we investigated whether sBCMA-Fc V3 with a 500-fold improvement in its binding affinity to BAFF (9 pM) would increase antitumor activity in BAFF driven DLBCL models. sBCMA-V3 Fc mediated inhibition of BAFF signaling pathway was evaluated *in vitro* in BAFF dependent SU-DHL-6 and Daudi DLBCL cell lines. sBCMA-Fc V3 treatment effectively reduced both DLBCL cell line growth in a dose-dependent manner, suggesting these cells are rely on BAFF for growth and survival (Fig. 5D). Treatment with sBCMA-Fc V3 also led to decreased expression of pAKT, p38-MAPK and pNF-*κ*B p52, all of which are critical molecular components of the BCMA/TACI/BAFF-R canonical signaling pathways that governs malignant B cell growth (Hatzoglou et al., 2000) (Sup. Fig. 10B). To determine the therapeutic efficacy of sBCMA-V3 Fc *in vivo*, SU-DHL-6 and Daudi tumor models were established by subcutaneous engraftment in *NOD-scid* mice. Animals were treated with 5 mg/kg or 10 mg/kg of sBCMA-Fc V3 and compared to the vehicle control. In both DLBCL tumor models, sBCMA-Fc V3 treatment resulted in significant dose-dependent reduction in tumor size (Fig. 5E, F). To further study the therapeutic potential of wild-type sBCMA-Fc compared to sBCMA-Fc 3V in DLBCL treatment, we added additional experimental groups treating SU-DHL-6 tumors with high doses of wild-type sBCMA-Fc at 10mg/kg and 20mg/kg. Similar to what we observed in previous studies (Fig. 2K), 10mg/kg wild-type sBCMA-Fc had little effect on tumor growth, but tumor reduction was observed with the 20mg/kg treatment group (Sup. Fig. 10C). This data provides evidence that wild-type sBCMA-Fc has suboptimal BAFF neutralizing capabilities and requires higher concentrations for BAFF inhibition of DLBCL tumor growth.

To further investigate the therapeutic the therapeutic efficacy of sBCMA-Fc V3, we compared it to other therapeutic agents known to inhibit BAFF in a head-to-head study. These other agents were a recombinant soluble TACI-Fc (sTACI-Fc) decoy receptor that binds APRIL and BAFF at 6.4 nM and 160 pM respectively (Wu et al., 2000), as well as an antibody against BAFF (*α*BAFF Ab) with a K_D_ of 0.995 nM and no reported binding to APRIL (Shin et al., 2018). The clinical versions of both molecules are currently used for the treatment of various B cell related autoimmune diseases (Bag-Ozbek and Hui-Yuen, 2021; Barratt et al., 2020; Lee and Amengual, 2020). We performed an *in vivo* head-to-head comparison study evaluating the antitumor efficacy of sBCMA-Fc V3, sTACI-Fc, *α*BAFF Ab and wild-type sBCMA-Fc in SU-DHL-6 DLBCL tumors. All therapeutic groups were treated with a dose of 10mg/kg, 3 times per week for decoy receptors and 2 times per week for antibody. sBCMA-Fc V3 treated tumors resulted in significant and robust tumor reduction compared to the vehicle control alone. sTACI-Fc treated groups showed a trend towards reduced tumor growth but didn’t reach statistical significance. In contrast, *α*BAFF Ab treated animals showed a modest but statistically significant reduction in tumor growth. Consistent with our previous observation, wild-type sBCMA-Fc did not show significant therapeutic benefit due to insufficient BAFF binding (Fig. 2G).

In summary, sBCMA-Fc V3 with enhanced binding affinity to APRIL and BAFF resulted in better antitumor activity in both APRIL driven MM and BAFF driven DLBCL models. In particular, when compared to wild-type sBCMA-Fc, sTACI-Fc and *α*BAFF Ab, sBCMA-Fc V3 treatment resulted in superior antitumor efficacy in DLBCL model, most likely to be driven by its engineered affinity binding towards BAFF. This observation further supported the therapeutic potential of sBCMA-Fc V3 as a treatment for MM and DLBCL.

### sBCMA-Fc V3 Demonstrates Adequate Toxicity Profile and On-Target Mechanism of Action in Nonhuman Primates

Nonhuman primate toxicity studies are particularly useful because they often respond in a similar physiologic manner as humans. To evaluate the translational potential of sBCMA-Fc V3, we conducted a single dose toxicity study in cynomolgus monkeys to investigate drug mediated acute toxicity after intravenous infusion of sBCMA-Fc V3.

One animal from each sex was assigned to 5 groups and given a single intravenous infusion of sBCMA-Fc V3 (0.1, 1, 10 and 100 mg/kg) or vehicle (Table 4). Animals were observed for 2 weeks before and 6 weeks after the treatment. Parameters evaluated include body weight, food consumption, hematology, coagulation, plasma chemistry, lymphocyte immunophenotype, immunoglobulin production, cytokines and gross pathology. Blood samples were collected from each animal during days -13, -6, -3 and 1 before dosing, then on days 2, 7, 14 and 42 during dosing (Table 4). Overall, there was no unscheduled deaths in the study. In addition, no drug-related abnormalities of body weight or food consumption were observed for the animals in each group during the observation period (Fig. 6A). Hematology analysis showed modest declines of RBC, HGB, and HCT in female and male monkeys of each group on Day 2, Day 7, and/or Day 14 (Sup. Table 5 to 8). However, when factor in the total volume of blood sampled during the experiment, it was likely that the decreases in RBC, HGB, and HCT may be related to blood sampling. No abnormalities were observed in coagulation (Sup. Table 9, 10) and plasma chemistry (Sup. Table 11 to 16). However, a dose-dependent, transient reduction in total lymphocyte numbers in female monkeys on day 7 post-dose followed by a rebound was observed. A similar but less significant trend was observed in male monkeys (Fig. 6B). We further investigated specific T cell and B cell subpopulation including CD4^+^ T cells and CD8^+^ T cells (Fig. 6C), as well as CD19+ pan B cells and CD20^+^ mature B cells in the test subjects at different time points (Fig. 6D). We found no significant differences upon treatment with a single dose of sBCMA-Fc V3 up to 100mg/kg while changes associated with multiple dosing require further study. No gross abnormal tissue pathology was present in animal tissues examined during necropsy.

**Table 4:** sBCMA-Fc V3 Single Dose Toxicology Study Design.

| Group | Number of animals |  | Animal number |  | Treatment | Dose level | Concentration | Dose volume |
| --- | --- | --- | --- | --- | --- | --- | --- | --- |
|  | Male | Female | Male | Female |  | mg/kg | mg/mL | mL/kg |
| 1 | 1 | 1 | 1101 | 2102 | Vehicle | 0 | 0 | 10 |
| 2 | 1 | 1 | 1203 | 2204 | sBCMA V3 | 0.1 | 0.01 | 10 |
| 3 | 1 | 1 | 1305 | 2306 | sBCMA V3 | 1 | 0.1 | 10 |
| 4 | 1 | 1 | 1407 | 2408 | sBCMA V3 | 10 | 1 | 10 |
| 5 | 1 | 1 | 1509 | 2510 | sBCMA V3 | 100 | 10 | 10 |

All dose levels/concentrations in the table are nominal.

**Table 5:** Changes of Immunoglobulin in the Females Dosed From 1 to 100 mg/kg.

| Parameters | Animal Number | Dose Level (mg/kg) | Day -6 | Day 2 | Day 7 | Day 14 | Day 42 |
| --- | --- | --- | --- | --- | --- | --- | --- |
| **IgA (g/L)** | 2306 | 1 | 1.33 | / | -20% | -35% | / |
|  | 2408 | 10 | 0.9 | / | -30% | -50% | -61% |
|  | 2510 | 100 | 0.98 | / | -27% | -52% | -87% |
| **IgG (g/L)** | 2408 | 10 | 9.12 | / | / | -22% | / |
|  | 2510 | 100 | 12.24 | / | / | -25% | -37% |
| **IgM (g/L)** | 2306 | 1 | 1.49 | / | -24% | -40% | / |
|  | 2408 | 10 | 0.96 | / | -32% | -43% | -51% |
|  | 2510 | 100 | 0.93 | / | -20% | -42% | -74% |

Formula: (Parameter Day n - Parameter Day -6)/ Parameter Day -6 ×100%. /: No abnormality.

**Table 6:** Changes of Immunoglobulin in the Males at 1 to 100 mg/kg.

| Parameters | Animal Number | Dose Level (mg/kg) | Day -6 | Day 2 | Day 7 | Day 14 | Day 42 |
| --- | --- | --- | --- | --- | --- | --- | --- |
| IgA (g/L) | 1305 | 1 | 1 | / | -19% | -44% | / |
|  | 1407 | 10 | 0.84 | / | -18% | -31% | / |
|  | 1509 | 100 | 0.95 | -20% | -37% | -54% | -75% |
| IgG (g/L) | 1509 | 100 | 16.38 | / | -34% | -44% | -56% |
| IgM (g/L) | 1305 | 1 | 1.83 | / | -24% | -38% | / |
|  | 1407 | 10 | 0.85 | / | / | -25% | -29% |
|  | 1509 | 100 | 2.75 | -38% | -56% | -70% | -82% |

Formula: (Parameter Day n - Parameter Day -6)/ Parameter Day -6 ×100%. /: No abnormality.

A panel of cytokines including IL-2, IL-4, IL-5, IL-6, IL-10, IL-13, IL-17*α*, IL-21 TNF-*α* and IFN-*γ* were examined in the blood pre and post treatment (Sup. Table 17 to 20). At 100mg/kg, we observed a transient increase in IL-10, IL-17*α*, and IFN-*γ* only in male monkeys on day 1, 2 and 3 post-dosing comparing to pre-dosing measurements and vehicle treated animals, that appeared to be transient. However, this change was not observed in the female monkeys treated with the same dose. No further change in cytokine levels was observed in other dosing groups.

Most importantly, a dose and time dependent reduction in immunoglobulin levels were observed in female and male monkeys treated. sBCMA-Fc V3 lead to a reduction in IgA immunoglobulin in both male and female monkeys over time starting at day 7 post-dose with gradual decrease recorded up to study termination (42 days post-dosing). A similar result was observed with IgM levels and IgG but to a much lesser extent (Table 4 and 5). A dose-dependent reduction in immunoglobulin production confirms the on-target mechanism of sBCMA-Fc V3 on APRIL and BAFF mediated immunoglobulin production and class switching and is consistent with our previous observation that the inhibition of APRIL/BAFF mediated signaling leads to decreased immunoglobulin production (Fig. 1F). This finding supports our hypothesis that sBCMA-Fc V3 treatment inhibits APRIL and BAFF mediated biological activity in a dose-dependent manner and will be helpful in designing a dosing schedule for clinical trials.

## DISCUSSION

The activation of APRIL and BAFF, lead to the activation of downstream pro-survival signals that promote the growth of abnormal B cell malignancies and other pathologies (Bolkun et al., 2014; Briones et al., 2002). The roles of APRIL and BAFF have been well studied in MM and DLBCL and are attractive therapeutic targets (Carpenter et al., 2013; Chiu et al., 2007). However, the multifaceted ligands/receptors interaction between APRIL/BAFF and their receptors BCMA/BAFF-R/TACI in B cell malignancy is challenging to overcome. Past approaches have used a ligand neutralization approach for APRIL or BAFF using monoclonal antibodies or decoy receptors for BCMA/BAFF-R/TACI in a limited number of disease applications (Rossi et al., 2009; Shrestha et al., 2021; Tai and Anderson, 2019). By using a soluble BCMA decoy receptor fusion protein sBCMA-Fc with natural high-affinity to APRIL, we can successfully inhibit MM tumors that rely on APRIL stimulation, despite differences in the cytogenetic profile of MM. However, this approach was not effective for targeting BAFF driven DLBCL due to low binding affinity between BAFF and BCMA. To therapeutically target BAFF, we used a yeast surface display-based protein engineering approach to generate mutant sBCMA clones with ultra-high binding affinity against both APRIL and BAFF. The resultant therapeutic candidate sBCMA-Fc V3 led to superior antitumor activities in both APRIL driven MM and BAFF driven DLBCL as well as desirable safety profile.

While the biological dependence of BCMA signaling on MM cell has been well studied, BCMA has only been exploited as a pan-MM surface marker for targeted delivery of cytotoxic therapeutics (Tai and Anderson, 2019). By using both *in vivo* analysis and ribosome based deep-sequencing approach, we demonstrated the strong dependence of MM for BCMA signaling, which acts as a critical regulator of protein translation that govern a subset of cellular targets. Ribosome profiling analysis provided insight into the global translational landscape of MM cells upon the loss of BCMA and revealed distinct molecular targets that were post-transcriptionally controlled which otherwise would not be detected by measuring total mRNA transcripts (Cenik et al., 2015; Ingolia et al., 2019). Specifically, we found enrichment in pathways that govern protein translation, including translation initiation, elongation, and ribosome integrity. While this discovery will be the subject of an independent investigation, there is little doubt that protein translation plays an essential role in plasma cells, which serves as a power house of immunoglobulin production (Perini et al., 2021). The inhibition of BCMA results in the reduction of Ig immunoglobulins, which at high levels, is associated with infections, peripheral neuropathy and bone damage in MM patients (Collier and Jackson, 1953; Pawlyn and Jackson, 2019).

Owning to the low binding affinity between wild-type BCMA and BAFF, we engineered a high-affinity clone sBCMA V3 with single digit picomolar binding to both APRIL and BAFF. Computational based structural analysis on sBCMA V3 indicated that the primary driver of affinity enhancement in this clone is the mutation H19Y. Although H19 was previously reported as an important residue for binding between BCMA and BAFF, its interaction with APRIL was not described. Supporting this statement, the H19Y mutation was isolated from APRIL based affinity screening and also resulted in the enhancement of BAFF binding. Thus, H19Y mutation is a highly desirable mutation on BCMA for better binding to both APRIL and BAFF. Interestingly, it is also worth noting that three other mutations S16G, T36A and N53D were not previously characterized as residues associated with BCMA binding to APRIL or BAFF while clearly contributing to the binding and stability of the BCMA/APRIL and BCMA/BAFF co-complexes. These findings provide the impetus to solve the crystal structures of sBCMA V3 in co-complex with APRIL and BAFF, which may provide additional biophysical evidence to explain how these mutations driven affinity enhancement. Therapeutically, enhancement in BAFF binding and its associated structural modifications shifted the treatment paradigm of wild-type sBCMA-Fc from a treatment selective for MM to a therapeutic candidate suited for targeting multiple B cell pathological indications. The biological function of BAFF and APRIL is not limited to B cell malignancy but extends beyond to autoimmune disorders and diseases triggered by pathological B cells such as IgA nephropathy, a much broader clinical indication for sBCMA-Fc V3 is possible (Samy et al., 2017).

sBCMA-Fc V3 is found to be well tolerated in single dose toxicity study conducted in cynomolgus monkeys. More importantly, a dose dependent reduction in immunoglobulin production, possibly as a result of APRIL and BAFF driven Ig class switching was observed in both male and female without affecting the mature B cell population. This data may partly explain our findings in MM tumor bearing mice, where genetic loss of BCMA signaling led to reduced abnormal immunoglobulin production, which is a key feature of MM pathogenesis. Overall, sBCMA-Fc V3 is found to be safe and well tolerated in all treated animals for up to 42 days with no significant clinical or pathological abnormality observed in dose up to 100mg/kg. Hence the maximal tolerated dose (MTD) is determined to be higher than 100mg/kg at single dose.

In summary, here we further explored the dichotomous function of APRIL and BAFF in MM and DLBCL, and engineered a high-affinity sBCMA-Fc V3 fusion protein capable of neutralizing APRIL and BAFF at low picomolar affinity. In so doing, we have transformed wild-type sBCMA-Fc decoy receptor to a high-affinity APRIL and BAFF dual ligand trap with superior antitumor activities in models of MM and DLBCL. More importantly, sBCMA-Fc V3 has a favorable safety profile and on-target mechanism of action in both murine and nonhuman primate models. Collectively, these data support sBCMA-Fc V3 as a clinically viable candidate for the treatment of APRIL and BAFF driven B cell malignancies and beyond.

## MATERIAL AND METHOD

### Study Approval

This study was designed to characterize the therapeutic efficacy and biological functionality of engineered soluble BCMA decoy receptor as a treatment for APRIL and BAFF driven Multiple Myeloma (MM) and Diffuse Large B Cell Lymphoma (DLBCL). For MM and DLBCL, patient specimens were collected from patients treated at Stanford Cancer Center under the approval of Stanford IRB No.13535. Healthy blood specimens were obtained from Stanford Blood Center under the same IRB protocol. *In vivo* animal studies were conducted under the approval of AAAPLAC at Stanford University. Sample sizes for animal were determined based on power calculations previously conducted *in vivo* studies. All animals were randomly assigned to treatment groups. Samples were not excluded from studies except for animals that required early termination due to unforeseeable illness that are unrelated to the study. Endpoints of experiments were defined in advance for each experiment. Tumor growth curves were presented for studies where tumor growth was measurable, serum levels of Myeloma protein levels were used as a marker of tumor progression in MM orthotopic PDX model and Kaplan-Meier analysis were used to define survival advantages in the PDX study whereas all other studies with measurable subcutaneous tumor uses final tumor growth as study endpoint. Appropriate statistical analysis was used for each study.

### Primate Toxicology Studies

The purposes of this study were to evaluate the acute toxicity after single administration of sBCMA-V3 via intravenous infusion in cynomolgus monkeys, to provide the maximum tolerated dose (MTD) as reference for the design of subsequent toxicity studies and clinical trials, andto characterize the toxicokinetics and immunogenicity. This study was contracted to Center for Drug Safety Evaluation and Research (CDSER), Shanghai Institute of Materia Medica (SIMM), Chinese Academy of Sciences (CAS) Address: 501 Haike Road, Zhangjiang Hi-Tech Park,Pudong New Area, Shanghai, P.R.China. This study was approved by CDSER-SIMM ethics committee for experimental usage and performed under the guidelines of NMPA: Guideline on Single Dose Toxicity Studies for Pharmaceuticals, May, 2014. NMPA: General Guideline on Non-clinical Safety

Evaluation for Therapeutic BiologicalProducts, January 2007. ICH Guideline M3 (R2): Guideline on Nonclinical Safety Studies for the Conduct of Human Clinical Trials and Marketing Authorization for Pharmaceuticals. CPMP/ICH/286/95. June 2009. ICH S6 (R1): Guideline on Preclinical Safety Evaluation of Biotechnology – Derived Pharmaceuticals, June 2011. Justification for Selection of Animal Species, Number of Animals and Route ofAdministration is as follows; 1) The cynomolgus monkey is considered as appropriate non-rodent species for safety evaluation of sBCMA-Fc V3. 2) The minimum number of animals used in this study met the requirements of scientific evaluation of the toxicity of the test article. 3) sBCMA-Fc V3 was administered via intravenous infusion in this study because intravenous administration is the intended administration route in human.

### Cell Lines

Human MM cell lines U266, MM1.R, INA6 and RPMI-8226 cells were maintained in RPMI-1640 media supplemented with 10% fetal bovine serum (FBS) and 1% Penicillin/Streptomycin in standing flasks and a humidified 37 °C, 5% CO_2_ incubator. INA6 Cells were supplemented with 2ng/ml of human IL-6 to maintain its growth. All cell lines were generously gifted by Dr. Albert Koong and Dr. Dadi Jiang at the Department of Radiation Oncology, MD Anderson (Houston, TX). Isolation of B-cells from healthy donor and MM patients were performed using EasySep^TM^ Human B-Cell Isolation Kit according to manufacturer’s protocol (Cat. No. 17954, Stemcell Technologies, Vancouver, Canada). DLBCL cells SU-DHL-6 and Daudi were purchased from ATCC (Cat. No. CRL-2959 and CCL-213, ATCC, Manassas, VA).

### Establishing MM PDX Models Using MM Patient Specimens

Mononuclear cells were isolated from bone marrow aspirate of Multiple Myeloma patients and inoculated into the left tibia of 5-6 weeks NSG mice. Injection path into the tibia was established using an empty needle penetrating through the tibia bone guided by X-ray, then the empty needle is removed, and X-ray turned off to avoid MM cells exposing to radiation. Patient cells were inoculated using a fresh needle and syringe. MM tumor growth was monitored by serum level of human IgG (M) protein. Once the host mice showed successful engraftment marked by increase of serum human IgG level, the animal was sacrificed, bone marrow flushed, and mononuclear cells collected for intratibial injection into 2 host mice. This process is repeated until sufficient N number is reached for each study. 11 patient samples were inoculated, 2 patient samples were successfully propagated for *in vivo* studies.

### *In vivo* Studies

All animal experiments were reviewed and approved by the Institutional Animal Care and Use Committee (IACUC) at Stanford University. Female NOD-*scid* gamma mice aged 6-8 weeks were purchased from the Jackson Laboratory (Stock no. 005557) and used for all *in vivo* analysis throughout the study. Mice were housed in pathogen-free animal facility, kept under controlled environment with 12 hours light-dark cycles. For INA6 and MM1.R tumor studies including (dox-inducible BCMA KO, WT sBCMA-Fc and sBCMA-Fc V3), 1x10^7^ cells were injected subcutaneously with 50% growth factor reduced Matrigel (Cat. no. 356230, Corning, NY). Bodyweight and tumor growth were measured 3 times a week until study termination. Animals were terminated upon subcutaneous tumor reaches ethical termination point. For PDX studies, non-terminal bleeding was performed on animals every 14 days for evaluating serum M protein as markers of tumor progression. Animals were terminated once shown signs of physical distress. For DLBCL tumor studies, Female NOD-*scid* gamma mice aged 6-8 weeks were purchased from the Jackson Laboratory (Stock no. 001303). Mice were house in the same pathogen-free animal facility where mice for MM studies were kept. For SU-DHL-6 and Daudi tumor studies, 5x10^6^ cells were injected subcutaneously with 50% growth factor reduced Matrigel (Cat. no. 356230, Corning, NY). Bodyweight and tumor growth were measured once a times a week until study termination. Wild-type sBCMA-Fc, sBCMA-Fc V3 was manufactured by ChemPartner Shanghai (Shanghai, P.R. China) using HEK293 transient expression and protein G purification system. Purified material was assessed by SEC-HPLC and SDS-PAGE for quality control. *α*CD38 antibody (Cat no. A2027) was purchased from SelleckChem (Houston, Tx), recombinant mouse soluble TACI-Fc (Cat. no. 577708) was purchased from Biolegend (San Diego, CA), *α*BAFF antibody was purchased from Invitrogen (Cat. no. MA1-822774, ThermoFisher Scientific, Waltham, MA)

### Ribo-Seq RNA Library Preparation

Briefly, snap frozen cell pellets (100 million cells per sample) were lysed in polysome lysis buffer (20 mM Tris-HCl pH 7.5, 250 mM NaCl, 15mM MgCl2, 1mM DTT, 0.5% Triton X-100, 0.024 U/ml TurboDNase, 0.48 U/ml RNasin, and 0.1 mg/ml cycloheximide). Lysates were centrifuged for 10 minutes at 4C, 14,000 x g. The supernatant was used for the isolation of ribosome bound mRNA, and total mRNA sequencing. SUPERase-In (0.24U/ml) was added to the lysate used for polysome fractionation to prevent RNA degradation.

### Ribosome Sequencing Analysis

The sequencing files for ribosome profiling and RNA-Seq data were processed using RiboFlow (Ozadam et al., 2020). All source code is freely available at https://github.com/ribosomeprofiling. Briefly, 3’ adapter sequence (AAAAAAAAAA) was removed from all reads using cutadapt. The 5’ end of each read includes 3 bases from the template switching reaction and also removed before alignment. We used a sequential alignment strategy to first filter out rRNA and tRNA mapping reads followed by mapping to representative isoforms for each as defined in the APPRIS database (Rodriguez et al., 2018). Next, PCR duplicates were removed from the ribosome profiling data using the 5’end of the sequence alignment coordinates. Finally, the resulting information was compiled into a .ribo file (Ozadam et al., 2020) for downstream analyses.

### Bioinformatic Analysis

All statistical analysis were carried out using RiboR (Ozadam et al., 2020). For quantification of ribosome occupancy, footprints of length between 26 and 30 (both inclusive) nucleotides were used. Metagene plots were generated using the 5’ end of each ribosome footprint. Ribosome occupancy and RNA-Seq data was jointly analyzed, and transcript specific dispersion estimates were calculated after TMM normalization (Robinson and Oshlack, 2010). To identify genes with differential translation efficiency, we used a generalized linear model that treats RNA expression and ribosome occupancy as two experimental manipulations of the RNA pool of the cells as previously described (Cenik et al., 2015). The model was fit using edgeR (Robinson and Oshlack, 2010) and p-values were adjusted for multiple hypothesis testing using Benjamin-Hochberg correction.

We used an adjusted p-value threshold of 0.05 to define significant differences. R packages cowplot, pheatmap, EnhancedVolcano, ggpubr, ggplot2, and reshape2 were used for analyses and plotting (Kassambara and Moreaux, 2018).

### Synthesis of Yeast-displayed sBCMA Library

DNA encoding human BCMA extracellular domain, amino acids Met1 – Ala54, was cloned into the pCT yeast display plasmid using *NheI* and *BamHI* restriction sites. Sequence numbering was done to match that used in Sasaki *et al*. to facilitate comparisons to their work with the wild-type proteins. An error-prone library was created using the BCMA extracellular domain DNA as a template and mutations were introduced by using low-fidelity Taq polymerase (Invitrogen, ThermoFisher Scientific, Waltham, MA) and the nucleotide analogs 8-oxo-dGTP and dPTP (TriLink Biotech). Six separate PCR reactions were performed in which the concentration of analogs and the number of cycles were varied to obtain a range of mutation frequencies: five cycles (200 μM), ten cycles (2, 20, or 200 μM), and 20 cycles (2 or 20 μM). Products from these reactions were amplified using forward and reverse primers each with 50 bp homology to the pCT plasmid in the absence of nucleotide analogs. Amplified DNA was purified using gel electrophoresis and pCT plasmid was digested with *NheI* and *BamHI*. Purified mutant cDNA and linearized plasmid were electroporated in a 5:1 ratio by weight into EBY100 yeast where they were assembled *in vivo* through homologous recombination. Library size was estimated to be 2×10^8^ by dilution plating.

### Library Screening

Yeast displaying high affinity BCMA mutants were isolated from the library using fluorescence-activated cell sorting (FACS). For FACS round 1, equilibrium binding sorts were performed in which yeast were incubated at room temperature in phosphate buffered saline with 0.1 % BSA (PBSA) with the 2 nM APRIL (Peprotech) for 24 h. After incubation with APRIL, yeast were pelleted, washed, and resuspended in PBSA with 1:100 mixture of anti-c-Myc FITC antibody (Abcam) and anti-HA AF647 (Invitrogen) for 1 hr at 4 °C. Yeast were then washed, pelleted and resuspended using PBSA followed by FACS analysis. For FACS rounds 2 – 6, kinetic off-rate sorts were conducted in which yeast were incubated with 2 nM APRIL for 3 hours at room temperature, after which cells were washed twice to remove excess unbound APRIL, and resuspended in PBSA containing a ∼50 fold molar excess of BCMA to render unbinding events irreversible. The length of the unbinding step was as follows: sort 2) 48 h; sort 3, 4 & 5) 72 h; sort 6) 84 h, with all unbinding reactions performed at room temperature. During the last hour of the dissociation reaction, cells were mixed with 1:100 mixture of anti-c-Myc FITC antibody (Abcam) and anti-HA AF647 (Invitrogen) for 1 hr at 4 °C. Yeast were pelleted, washed, and resuspended in 0.1% BSA. Labeled yeast were sorted by FACS using a Vantage SE flow cytometer (Stanford FACS Core Facility) and CellQuest software (Becton Dickinson). Sorts were conducted such that the 1–3% of clones with the highest APRIL binding/c-Myc expression ratio were selected, enriching the library for clones with the highest binding affinity to APRIL. In sort 1, 10^8^ cells were screened and subsequent rounds analyzed a minimum of ten-fold the number of clones collected in the prior sort round to ensure adequate sampling of the library diversity. Selected clones were propagated and subjected to further rounds of FACS. Following sorts 3, 4,5 and 6, plasmid DNA was recovered using a Zymoprep kit (Zymo Research Corp.), transformed into DH5a supercompetent cells, and isolated using plasmid miniprep kit (Qiagen). Sequencing was performed by MCLAB.

Analysis of yeast-displayed sort products was performed using the same reagents and protocols and described for the library sorts. Samples were analyzed on a FACS Calibur (BD Biosciences) and data was analyzed using FlowJo software (Treestar Inc.).

### Binding Affinity Assay

Cells were cultured in standard tissue culture condition. Cells were harvested and the supernatant discarded then dispensed onto a staining plate at 3x10^5^ cells per well. The plate was centrifuged at 300g at 4°C for 5 minutes. Various concentrations of sBCMA mutants and negative control were diluted in FACS buffer containing 2% FBS, 100µL/well was added. Cells were incubated for 1 hour at 4°C and washed twice with 200µL FACS buffer and centrifuged at 300g for 5 minutes. The supernatant was discarded before and after each wash. Cells were re-suspended at 100µL/well with 1:1000 diluent with anti-human IgG-Alexa 488 (Cat. no. A28175, ThermoFisher, Waltham, MA). Plates were incubated for 1 hour at 4°C. Cells were washed twice with FACS buffer and centrifuged at 300g for 5 minutes. Supernatant was discarded and cells were re-suspended in 100µL cold PBS. The cells were kept in the dark and FACS analysis carried out on FACS CantoII, (BD Biosciences, San Jose, CA). Geometric mean (measure of binding affinity) of double positive population was determined by using FlowJo software. In order to determine the K_d_ (ligand concentration that binds to half the receptor sites at equilibrium) of the binding reaction, binding affinity was plotted against ligand concentration and the graph was analyzed using one site – specific binding in Graphpad Prism to get the Kd value.

### Computational Structural Simulation

Computation based structural simulation was carried out using a number of structural prediction software. sBCMA V3 in co-compled with hAPRIL and hBAFF was modeled using a combination of Prime from Schrödinger Suites 2021-2, Alphafold2, RoseTTAFold, trRosetta and RosettaRemodel based on sequences and structural alignment mapped to Protein Data Bank ID 1XU2. Mutation mediated changes within sBCMA V3 binding to hAPRIL and BAFF was calculated by Residue Scanning Calculation Module from Bioluminate, Schrödinger. Surface complementarity and protein-protein interaction between sBCMA V3 and hAPRIL/hBAFF was calculated by Protein Interaction Analysis module from Bioluminate, Schrödinger.

### In Vitro Cell Based Viability Assays

Cell viability was determined with Cell counting hemocytometer or Beckman coulter counter depending on the study. Cells were plated in 96-well plates at a density of 2500 cells (U266, SU-DHL-6 and Daudi) or 3000 cells (MM1.R and INA-6). For wild-type sBCMA-Fc and sBCMA-Fc V3 treatment, cells were cultured in 1% FCS RPMI media overnight followed by 1 hour of 100ng recombinant APRIL or BAFF stimulation, increasing doses of sBCMA were added to designated wells. Both APRIL and sBCMA-Fc treatment are replenished every 48 hours until experiment ends on day 7.

### Mice Computerized Tomography (CT) Scan to Confirm MM Induced Bone Degradation

High-resolution micro-CT images were acquired using an *in vivo* micro-CT scanner SkyScan 1276 (Bruker, Billerica, Massachusetts, US) under isoflurane anesthesia. The scanning mode was set as 360°, step-and-shoot scanning without average framing. After each scan, the projection images were reconstructed using the software (NRecon with GPU acceleration, Bruker), followed by converting the set of reconstructed slices to DICOM files (DICOM converter, Bruker).

### Enzyme-Linked Immunosorbent Assay (ELISA)

Serum and cell lysate expression of Human APRIL (Cat. no. DY884B, R&D Systems Minneapolis, MN), human BAFF (Cat. no. DBLYS0B, &D Systems Minneapolis, MN), mouse APRIL (Cat. no. MBS738004 My Biosource, San Diego, CA), mouse BAFF (Cat. no. MBLYS0, R&D Systems Minneapolis, MN), Human total IgG (M) protein (Cat. no. BMS2091, ThermoFisher, Waltham, MA), mouse total IgG (M) protein (Cat. no. 88-50400-88, ThermoFisher, Waltham, MA), Human IL-6 (Cat. no. EH2IL6, Invitrogen/ ThermoFisher, Waltham, MA) expressions were detected using commercial ELISA kit listed above according to manufacturer’s protocol.

### Immunohistochemistry

Tissues were fixed in 10% neutral buffered formalin and embedded in paraffin blocks for cutting and mounting on glass slides. For Ki67 staining, slides were de-paraffined and antigen retrieval carried out using 10mM Citric Acid Buffer, 0.05% Tween 20, pH 6 Slides were removed from buffer and cooled at room temperature for 15 minutes. Quenched endogenous peroxidase with 1:10 dilution of 34% hydrogen peroxide and water for 15 minutes. Avidin and Biotin blocker were added for 15 minutes each. Protein block using 2% fetal calf serum was added for 20 minutes. The serum and antibody were diluted in PBT (1XPBS, 0.1% BSA, 0.2%, 0.01% Tween 20). Anti-Human Ki67 antibody (Cat. no. sc-23900, Santa Cruz Biotechnologies, Dallas, Tx) incubated overnight at 4°C. Biotinylated anti-mouse secondary antibody 1:2500 (Cat. no. BA92001, Vector Laboratories, Burlingame, CA) was added on each slide and incubated at 37°C for 30 minutes then incubated with STREP-HRP for 30 minutes at 37°C. Signals developed using DAB substrate kit (#34002, ThermoFisher Scientific, Waltham, MA). TUNEL apoptosis assay was carried out using ApopTag ®Peroxidase *In Situ* Apoptosis Detection Kit (Cat. no. S7100, Millipore Sigma, Burlington, MA) and performed according to manufacturer’s instruction. All cases were scanned at 40x magnification using the Leica Aperio AT2 Digital Pathology Scanner (Leica Biosystem, Buffalo Grove, IL). Images taken were analyzed using NDP.view2 image analysis software developed by Hamamatsu Japan (Bridgewater, NJ).

### Immunoblotting

Cell lysates were subjected to sodium dodecyl sulfate polyacrylamide gel electrophoresis, followed by transfer to nitrocellulose membrane. The membranes were then probed with primary Abs against total BCMA (Cat. no. 27724-1-AP, Proteintech, Rosemont, IL), Pan-Akt (Cat. no. 4691 Cell Signaling Technology, Danvers, MA), pAkt (Cat. no. 4060, Cell Signaling Technology, Danvers, MA), phospho-p38 MAPK (Cat. no 4511, Cell Signaling Technology, Danvers, MA), p38 MAPK (Cat. no. 8690 Cell Signaling Technology, Danvers, MA), Phospho-mTOR ser2448 (Cat. no. 5536 Cell Signaling Technology, Danvers, MA), total mTOR (Cat. no. 2983 Cell Signaling Technology, Danvers, MA), Raptor (Cat. no. 2280 Cell Signaling Technology, Danvers, MA), Phospho-Raptor Ser792 (Cat. no. 89146 Cell Signaling Technology, Danvers, MA), Phospho-p70 S6K (Cat. no. 9204 Cell Signaling Technology, Danvers, MA), Phospho-S6 (Cat. no. 9204 Cell Signaling Technology, Danvers, MA) and b-Actin (no. sc-47778 HRP, Santa Cruz Biotechnology Inc. Dallas, Tx) at 4°C overnight. The blots were then washed and probed with HRP-conjugated anti-goat (no. sc-2020, Santa Cruz Biotechnology Inc. Dallas, Tx), or HRP-conjugated anti-rabbit (no. A16110, Thermo Fisher Scientific) as appropriate. The blots were developed with Bio-Rad Western C Developing Reagent (no. 170-5060 Bio-Rad) and visualized with Chemidoc digital imager (no. 1708280, Bio-Rad).

### *In Vitro* genetic knockdown studies

BCMA siRNA (SMARTPool Cat. No. L-011217-00-0005) and Doxycyclin inducible shRNA (SAMRTvector Cat. No. V3SH7669-230564302) constructs were purchased through GE Dharmacon Horizon (Cambridge, UK). For siRNA, transfection procedures were carried out using Lonza 4D-Nucleofactor device and kits in accordance with manufacturers protocol (Basal, Switzerland). For doxycycline inducible BCMA knockdown cells, 3 shRNA sequence were tested in according to the transfection protocols provided by the manufacturer. Sequence no. 1 and 2 showed successful knockdown of BCMA and was used for the subsequent *in vivo* testing.

### shRNA Sequences

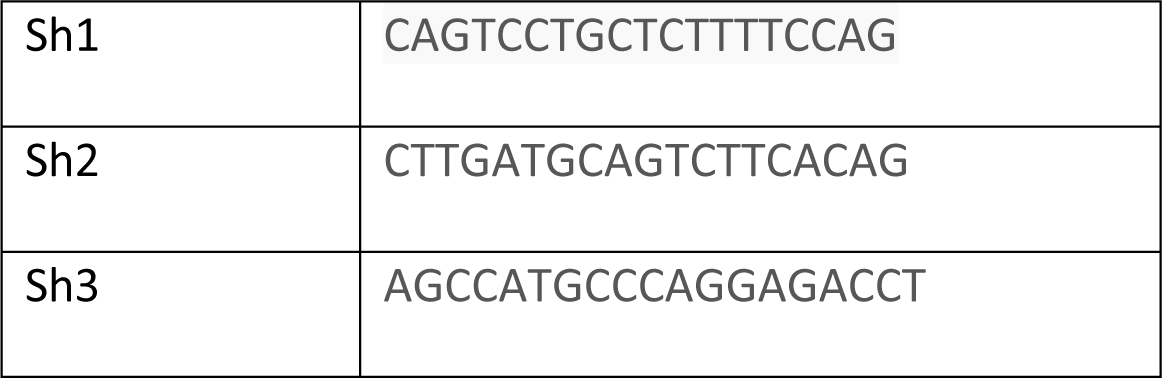

### Real-Time PCR analysis

RNA was isolated using TRIzol reagent according to the manufacturer’s instructions (Invitrogen). RNA was reverse transcribed using cDNA synthesis kit (Biorad, Hercules, CA). Real-time PCR was performed as previously described. Relative expression levels of target genes were normalized against the level of GAPDH expression. Fold difference (as relative mRNA expression) was calculated by the comparative CT method (2^Ct(GAPDH RNA–gene of interest)^).

### Primer Sequences

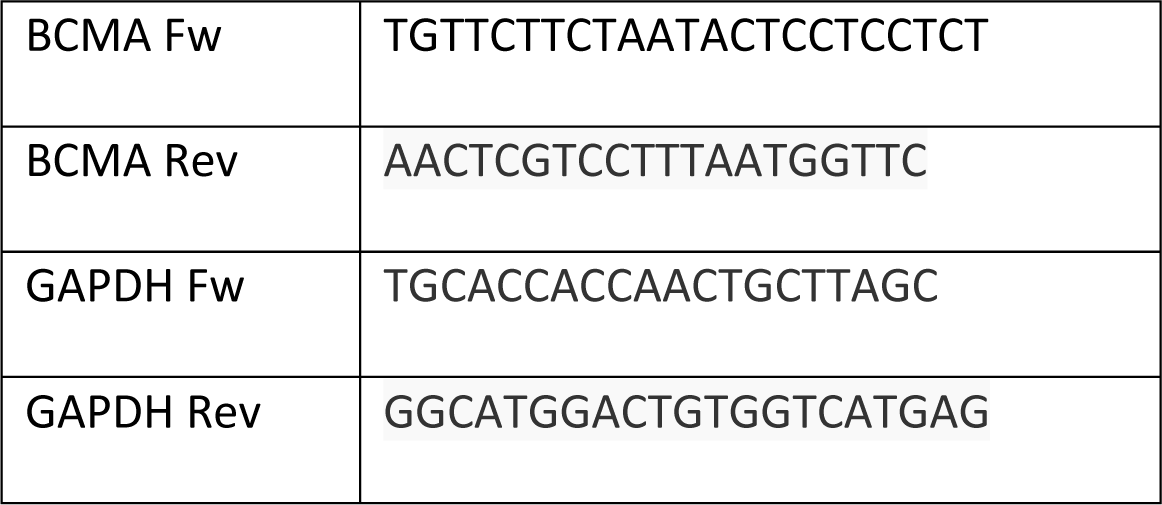

### Reverse Phase Protein Array (RPPA)

The RPPA was performed by MD Anderson RPPA core as described according to the published protocol.

### Statistical Analysis

All cell number, tumor volume, survival and quantification of *in vivo* and *in vitro* studies were conducted using GraphPad Prism software (GraphPad Software Inc, CA). ANOVA with Tukey-Kramer test was used for comparing multiple treatment groups with each other. *P* < 0.05 was considered significant. Repeated measure ANOVA was used for comparing multiple treatment groups measured over time. Statistical analysis of survival curves was conducted for the survival studies. A log-rank (Mantel-Cox) test was performed to compare mean survival among groups; *P* ≤ 0.05 was considered statistically significant.

## SUMMARY OF SUPPLEMENTAL MATERIALS

**Sup Fig. 1**: BCMA Signaling Activation is Required for Multiple Myeloma Progression.

**Sup Fig. 2**: Changes in MM proliferation and apoptosis upon genetic loss of BCMA.

**Sup Fig. 3**: Bioinformatic analysis of MM cells upon genetic loss of BCMA.

**Sup Fig. 4**: Changes in mRNA expression upon genetic loss of BCMA.

**Sup Fig. 5**: Altered protein translation machinery signaling upon loss of BCMA.

**Sup Fig. 6**: *In vivo* efficacy in MM models treated with wild-type sBCMA-Fc.

**Sup Fig. 7**: Demographic information and treatment status of MM PDX models.

**Sup Fig. 8**: *In vivo* and *in vitro* studies using wild-type sBCMA-Fc as a treatment in MM and DLBCL.

**Sup Fig. 9**: Computational structural alignment of BCMA V3/APRIL and BCMA V3/BAFF co-complex.

**Sup Fig. 10**: Therapeutic efficacy of sBCMA-V3 Fc in MM and DLBCL models.

**Sup. Table 1-4:** Top clones selected from rounds 3-6 of affinity-based flow cytometry sorting.

**Sup. Table 5-8:** Hematology analysis on male and female cynomolgus monkey dosed with sBCMA-Fc V3.

**Sup. Table 9 and 10:** Blood coagulation analysis on male and female cynomolgus monkey dosed with sBCMA-Fc V3.

**Sup. Table 11-16:** Chemistry panel analysis on male and female cynomolgus monkey dosed with sBCMA-Fc V3.

**Sup. Table 17-20:** Cytokine panel analysis on male and female cynomolgus monkey dosed with sBCMA-Fc V3.

## ACKNOWLEDGEMENTS AND CONFLICT OF INTEREST

The authors wish to thank Drs. Vignesh Viswanathan, Dhanya Nambiar, Quynh-Thu Le for sharing experimental reagents. We would like to Stanford animal facility for maintaining animal colonies, Chempartner Shanghai for the production of wild-type sBCMA-Fc and sBCMA-Fc V3, TB-seq for isolation and preparation of ribosome mRNA library and the University of Delaware sequencing facility for performing ribosome sequencing.

This work was supported by grants from The Silicon Valley Foundation (A.J.G), The Sydney Frank Foundation (A.J.G), The Kimmelman Fund (A.J.G), Medical Research Council (MRC) UK Grant (A.J.G).

This work was supported in part by the Cancer Prevention and Research Institute of Texas (RR180042 to C.C.) and Welch Foundation (F-2027-20200401 to C.C.)

YRM, AJG, KT and KM are co-inventors on the patent, “sBCMA Variants and Fc Fusion Proteins Thereof”. Provisional Application No. 63/064,880. YRM, AJG and XEZ have stock equity in AKSO Biopharmaceutical Inc.

## AUTHOR CONTRIBUTIONS

Conceptualization: YRM, KT, CC, DJ, KM, XEZ, ML, PA, ACK, AJG Methodology: YRM, KT, CC, DJ, KM, CGL, HZ, ML, PA, AL Investigation: YRM, KT, CC, DJ, KM, CJ, TTCY, CGL, HZ, AD, JX, XEZ Fund acquisition: AJG, CC Project administration: YRM, AJG Supervision: YRM, ACK, AJG Writing: YRM, KT, CC, CJ, DJ, AJG

