## Supplemental figures for "Developing High-Affinity Decoy Receptor Optimized for Multiple Myeloma and Diffuse Large B Cell Lymphoma Treatment"

#### Supplemental Figure 1

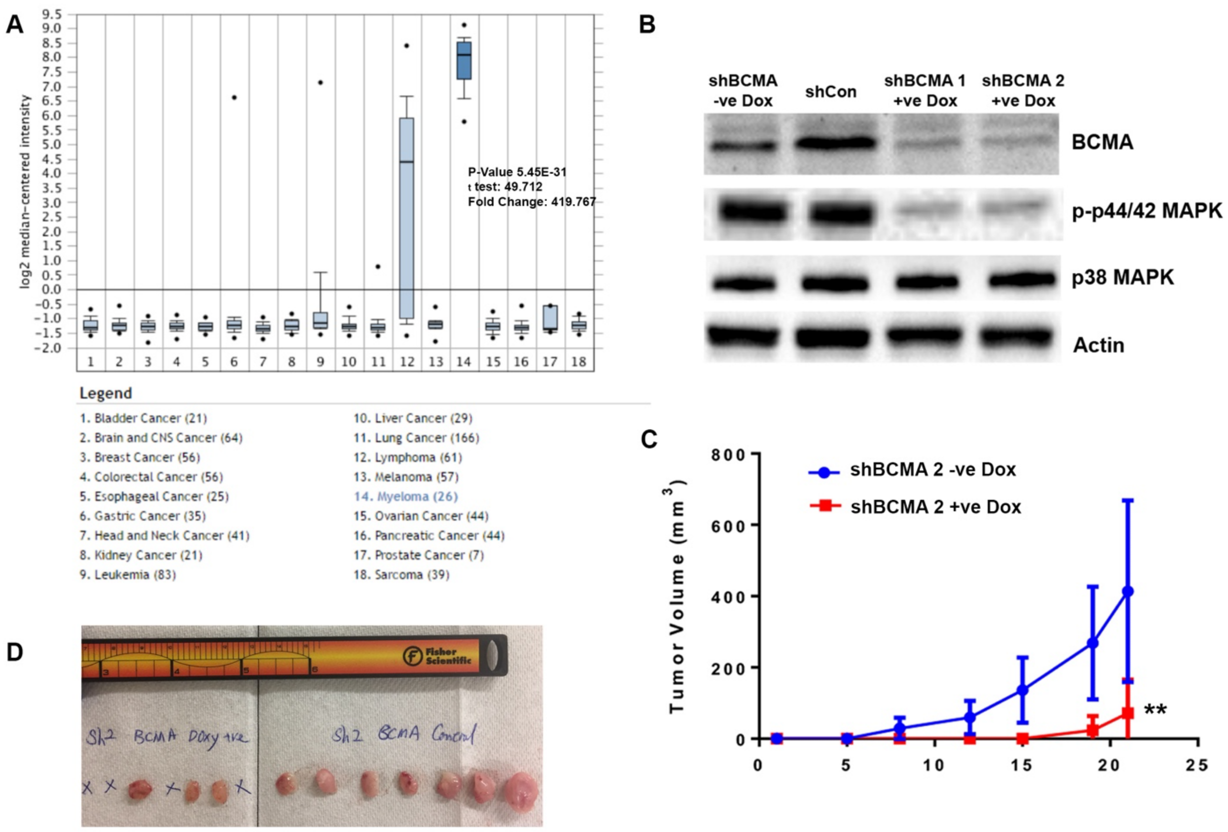


**Sup. Figure 1: (A)** BCMA mRNA expression in a panel of 19 tumor cell lines queried through Oncomine Database. **(B)** Western blotting analysis confirming knockdown of BCMA in MM cell line by Doxycycline-inducible shRNA. **(C)** Tumor growth kinetics in MM1.R MM cells transfected with inducible dox shBCMA, mice were given drinking water with or without doxycycline (5mg/ml). **(D)** Images of MM1.R tumors harvested from study described in **(C)**. Statistical analysis was conducted using One-way ANOVA for comparing between treatment groups and repeated ANOVA for changes occur over-time. P value *=<0.05, **=<0.01. ***=<0.001

**Supplemental Figure 2**

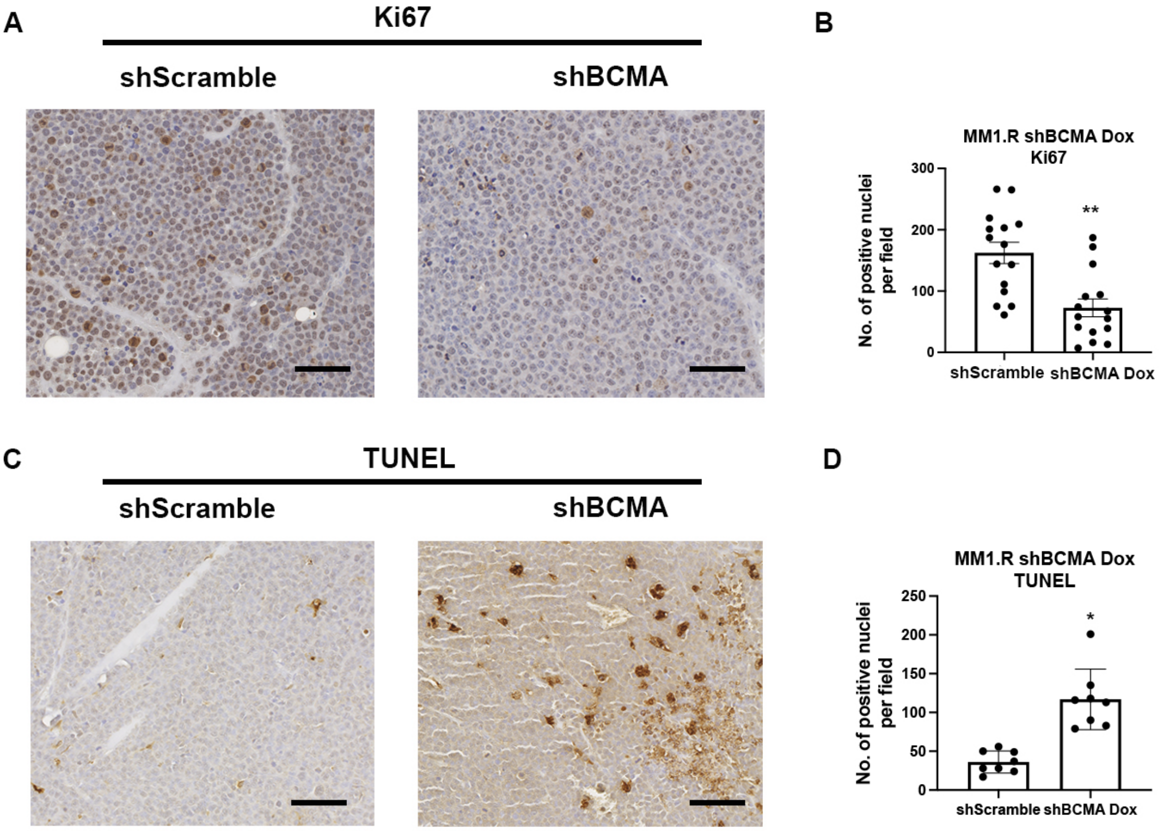


**Sup. Figure 2: (A)** Representative images of Ki67 positive cells in the harvested tumors of doxycycline-inducible shScrm and shBCMA analyzed by IHC staining. Scale bar 50μm. **(B)** Quantitative analysis of Ki67 positive cells in the harvested tumors of doxycycline-inducible shScrm Control and shBCMA, represented as the average number of positive nuclei per image field. **(C)** Representative images of TUNEL positive cells in the harvested tumors of doxycycline-inducible shScrm and shBCMA analyzed by IHC staining. Scale bar 50μm. **(D)** Quantitative analysis of TUNEL positive cells in the harvested tumors of doxycycline-inducible shScrm and shBCMA, represented as the average number of positive nuclei per image field. Statistical analysis was conducted using T test and one way ANOVA for comparing between treatment groups. Repeated ANOVA used for changes in tumor growth over time. P-value *=<0.05, **=<0.01. ***=<0.001.

**Supplemental Figure 3**

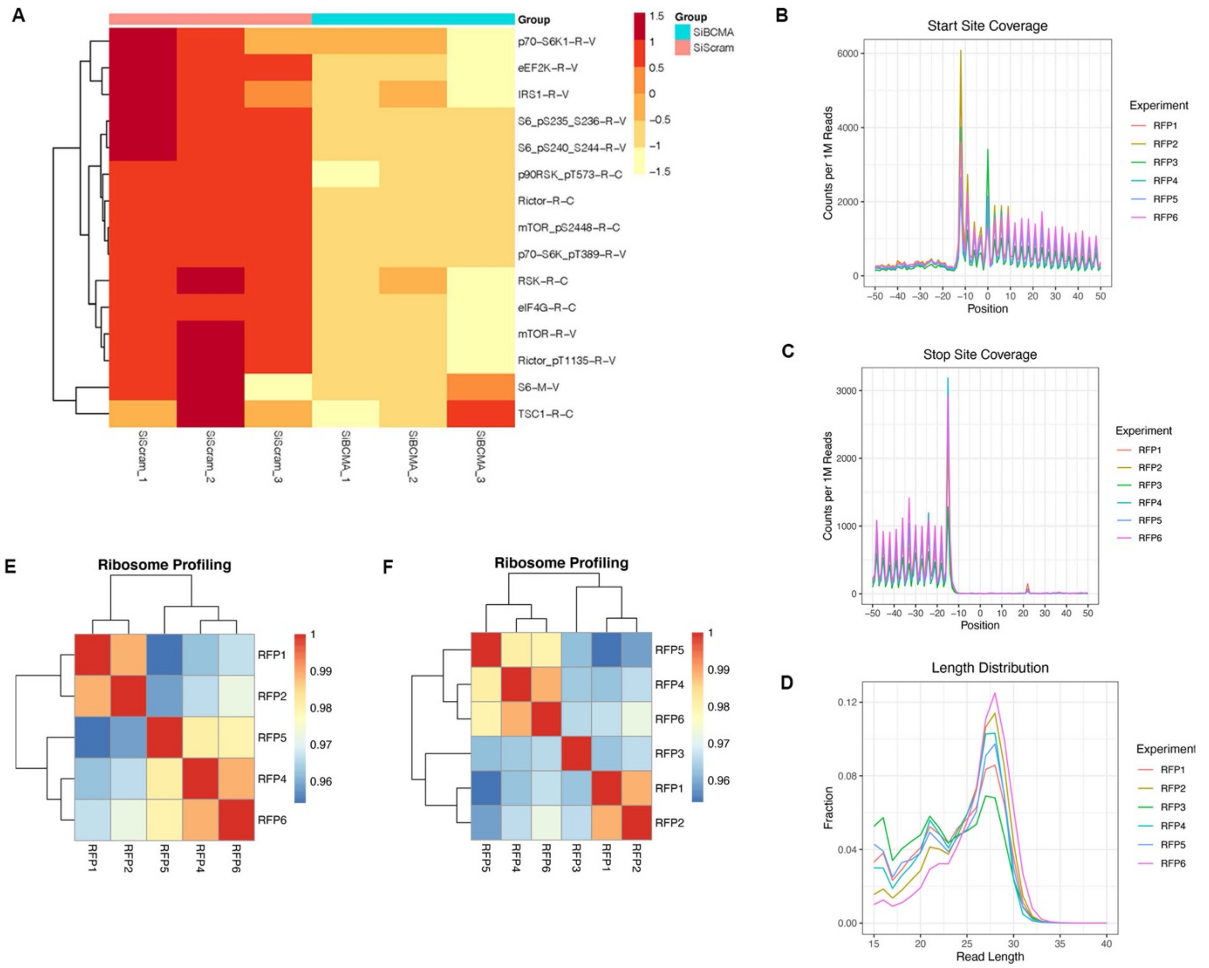


**Sup. Figure 3: (A)** Loss of BCMA induced changes in downstream targets associated protein translation analyzed through Reverse Phrase Protein Array. **(B, C, D)** Quality assessment of read length across all ribosome profiling samples sent for analysis. Three nucleotide periodicity, enrichment of footprints in coding regions and the expected signals at the translation start and stop sites. **(E)** Spearman Correlation analysis of Ribosome profiling samples showing consistent reads across all samples with the exception of RFP3. **(F)** Spearman Correlation analysis after RFP3 was removed from the dataset.

### Supplemental Figure 4

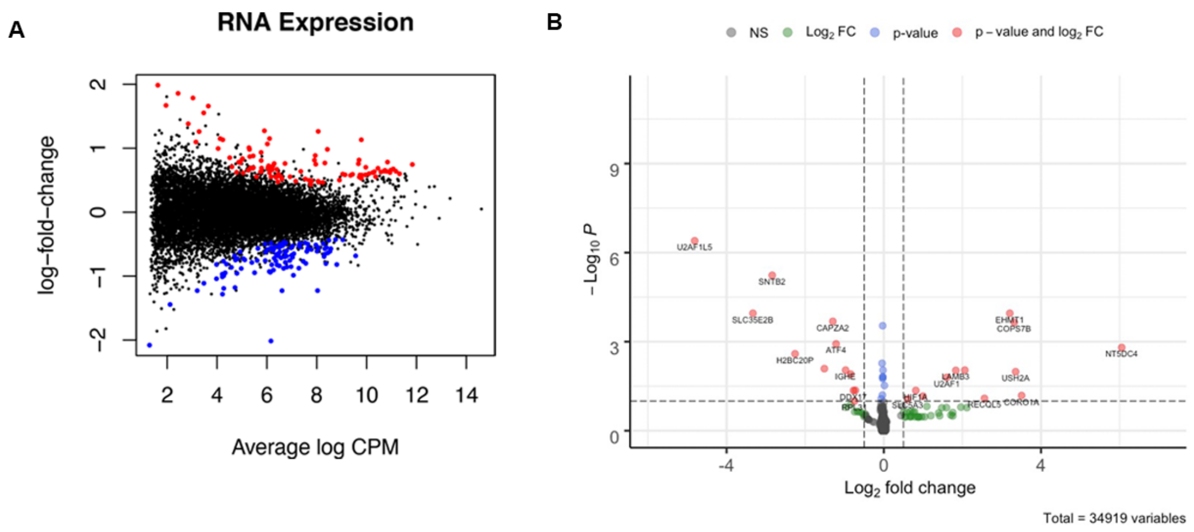


**Sup. Figure 4: (A)** Distribution plot of changes in RNA expression associated with genetic knockdown of BCMA. **(B)** Volcano plot analysis identifying downstream targets with significant changes in total RNA transcript categorized into $Log_{10}Fc$, p-value and both.

Supplemental Figure 5

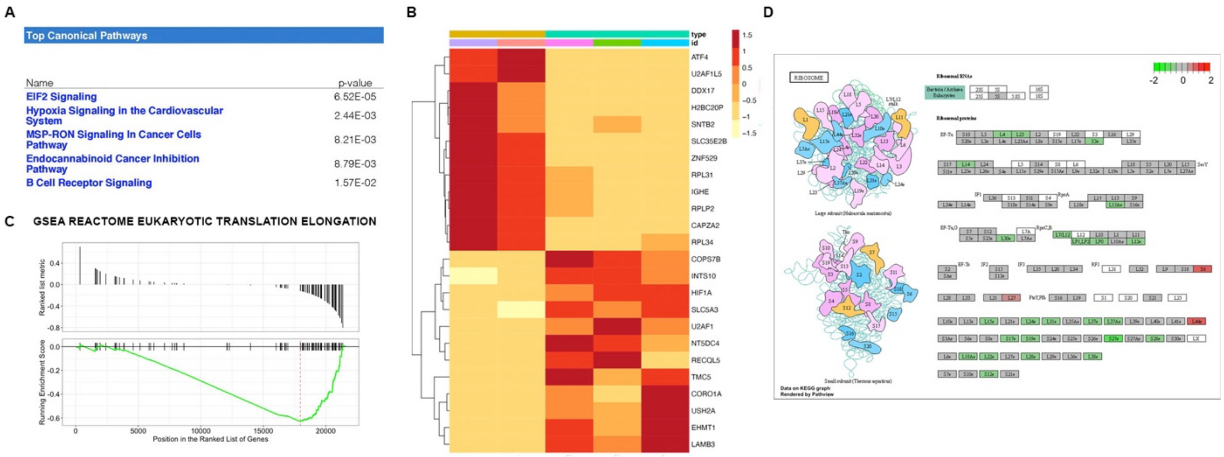


**Sup. Figure 5: (A)** Ingenuity Pathway Analysis of RNA-seq data showing top five significantly changed canonical pathways. **(B)** Representative heatmap of RPPA analysis showing subset of the significant changes in protein expression landscape associated with BCMA signaling axis. **(C)** GSEA REACTOME enrichment analysis showing enriched signatures in associated with protein translation elongation upon loss of BCMA. **(D)** Graphic illustration of large ribosome subunits (top) and small ribosome subunits (bottom). Each subunit showing changes in translation abundance upon BCMA loss. Decreased expression (green), no change (gray) or increased expression (red).

**Supplemental Figure 6**

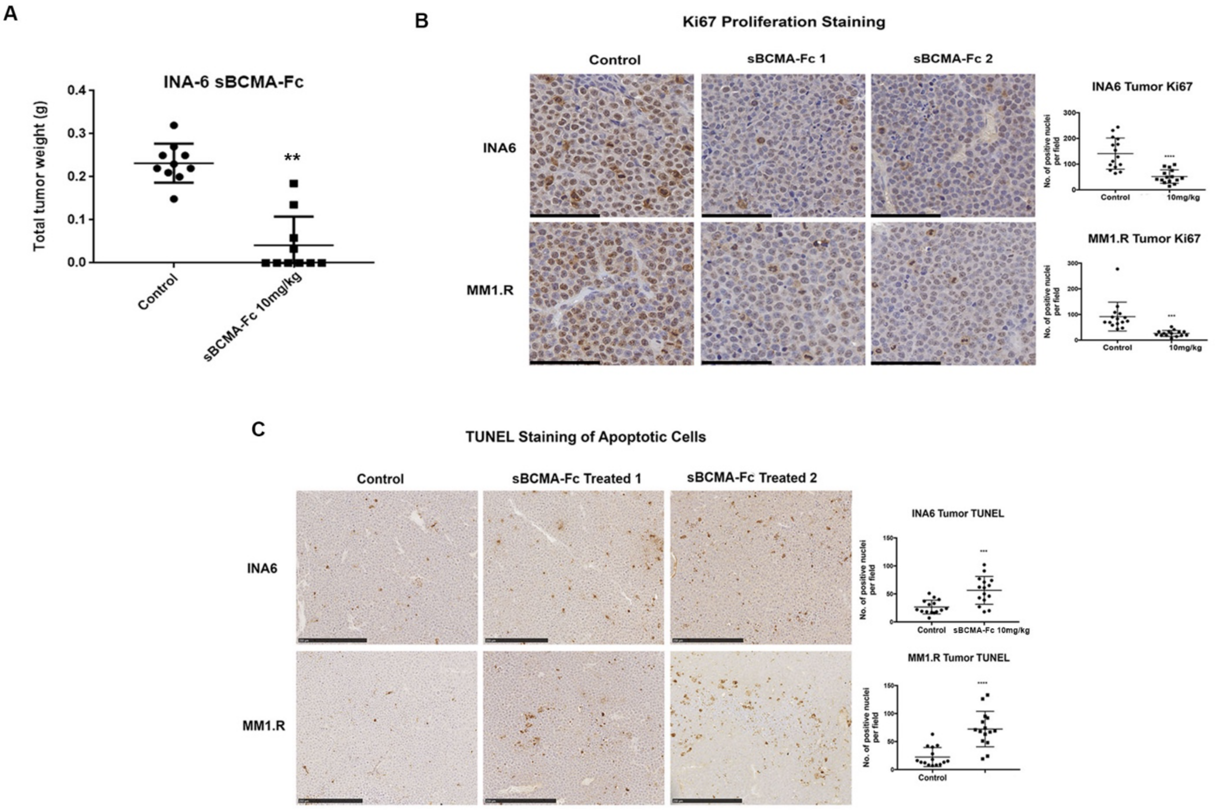


**Sup. Figure 6: (A)** Terminal tumor weight of mice inoculated with INA-6 MM tumors and treated with vehicle control or 10mg/kg of sBCMA-Fc (N=10). (B) Representative images of Ki67 positive cells in the vehicle control and sBCMA-Fc treated INA-6 (top panels) and MM1.R (bottom panels) MM tumors analyzed by IHC staining. Scale bar 50μm. Quantification of Ki67 staining on right. (C) Representative images of TUNEL positive cells in the vehicle control and sBCMA-Fc treated INA-6 (top panels) and MM1.R (bottom panels) MM tumors analyzed by IHC staining. Scale bar 50μm. Quantification of TUNEL staining on right. Statistical analysis was conducted using T test and one way ANOVA for comparing between treatment groups. Repeated ANOVA used for changes in tumor growth over time. P-value *=<0.05, **=<0.01. ***=<0.001.

**Supplemental Figure 7**

A

| Patient No. | Sex | Collection Date | Age | Treatment Status | Myeloma Type |
|---|---|---|---|---|---|
| 1 | Male | 6/1/18 | 53 | Untreated | Unknown |
| 2 | Female | 6/11/18 | 59 | Previously Treated | IgGk |
| 3 | Male | 7/25/18 | 53 | Untreated | IgGk |
| 4 | Male | 7/25/18 | 56 | Previously Treated | IgGl |
| 5 | Male | 8/2/18 | 60 | Untreated | IgGk |
| 6 | Male | 8/3/18 | 55 | Previously Treated | Unknown |
| 7 | Male | 9/18/18 | 66 | Untreated | IgGk |
| 8 | Female | 10/23/18 | 64 | Untreated | IgGl |
| 9 | Male | 10/23/18 | 71 | Relapse/Refractory | IgGk |
| 10 | Female | 2/13/19 | 49 | Untreated | Unknown |
| 11 | Male | 7/9/19 | 52 | Previously Treated | IgGk |

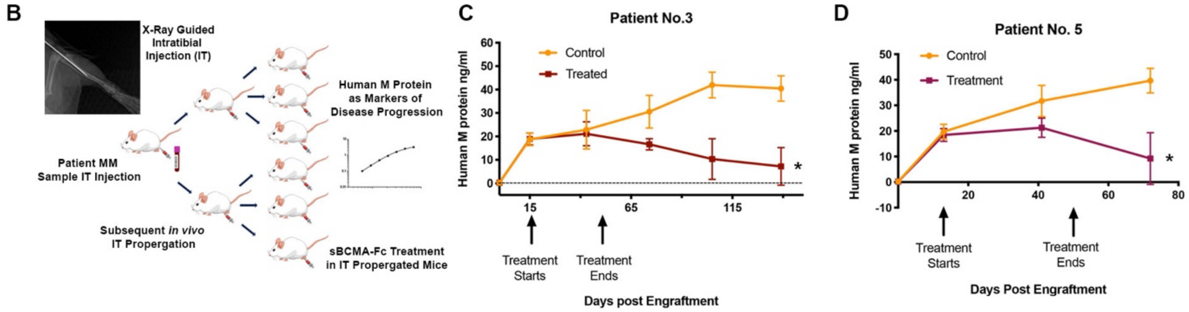


**Sup. Figure 7: (A)** Demographic information and treatment status of patient samples collected. **(B)** Schematic illustrations of *in vivo* MM PDX propagation. Patient tumor cells were isolated from patient bone marrow biopsies and injected into the tibia of the NSG mice. PDX were subsequently propagated *in vivo* using the same intratibial inoculation procedure and treated with vehicle control or sBCMA-Fc. Human IgG protein in mouse serum is continuously monitored over time as a marker of tumor progression. **(C)** Human IgG protein in mouse serum detected in animals successfully engrafted with MM cells from patient No. 3 showing reduction in Human M protein level post sBCMA-Fc treatment (N=8) compared to vehicle control (N=7). **(D)** Human IgG protein in mouse serum detected in animals successfully engrafted with MM cells from patient No. 5 showing reduction in Human IgG protein level post sBCMA-Fc treatment (N=10) compared to vehicle control (N=10). Statistical analysis was conducted using T test and one way ANOVA for comparing between treatment groups. Repeated ANOVA used for changes in tumor growth over time. P-value *=<0.05, **=<0.01. ***=<0.001.

**Supplemental Figure 8**

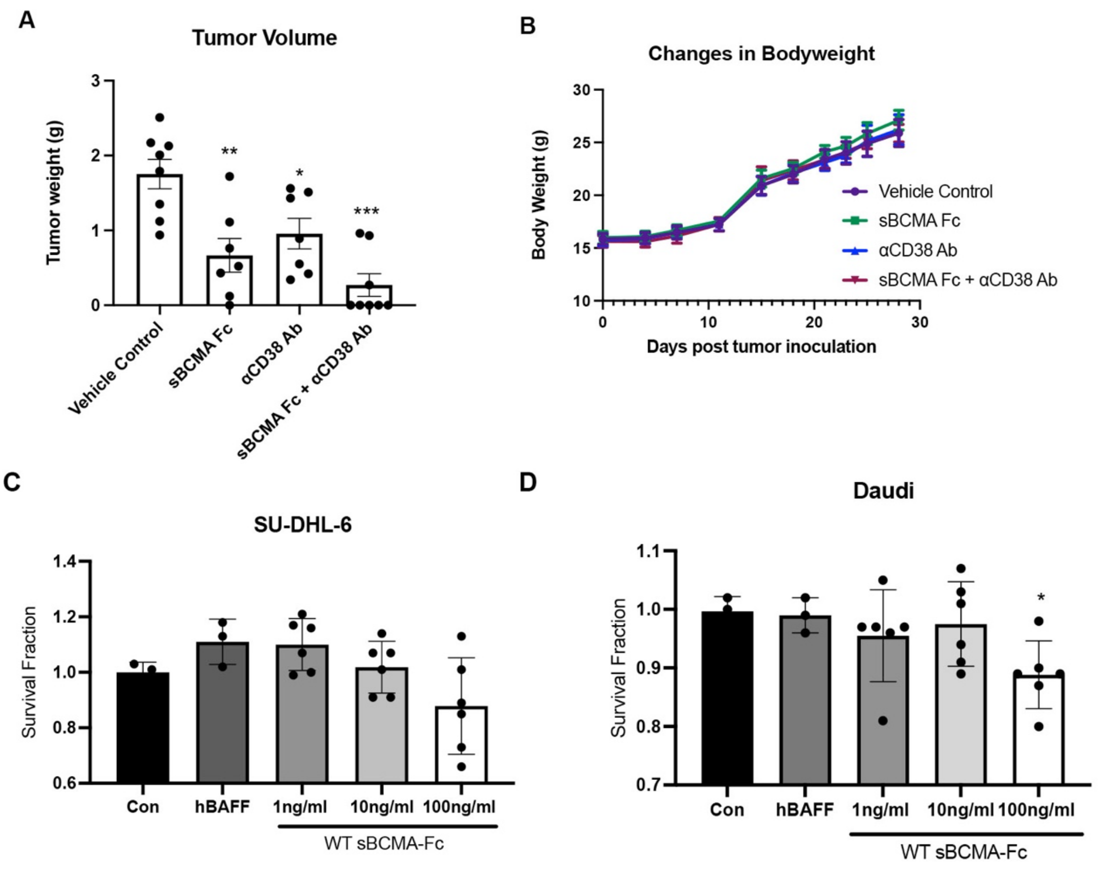


**Sup. Figure 8: (A)** Terminal tumor weight of MM1.R MM cells in mice dosed with sBCMA-Fc 10 mg/kg every 48 hours (N=7), $\alpha$CD38 10mg/kg weekly (N=7), sBCMA-Fc and $\alpha$CD38 combination (N=8) and vehicle control (N=8) in 6 weeks old female NSG mice. **(B)** Changes in bodyweight of animals from study described in (A). **(C)** sBCMA-Fc dose-dependent, cytotoxicity assay validating the *in vitro* cell survival in the presence of increasing doses of sBCMA-Fc and hBAFF (100ng/ml) in SU-DHL-6 DLBCL cells. **(D)** sBCMA-Fc dose-dependent, cytotoxicity assay validating the *in vitro* cell survival in the presence of increasing doses of sBCMA-Fc and hBAFF (100ng/ml) in Daudi DLBCL cells. Statistical analysis was conducted using T test and one way ANOVA for comparing between treatment groups. Repeated ANOVA used for changes in tumor growth over time. P value *=<0.05, **=<0.01. ***=<0.001.

Supplemental Figure 9

A

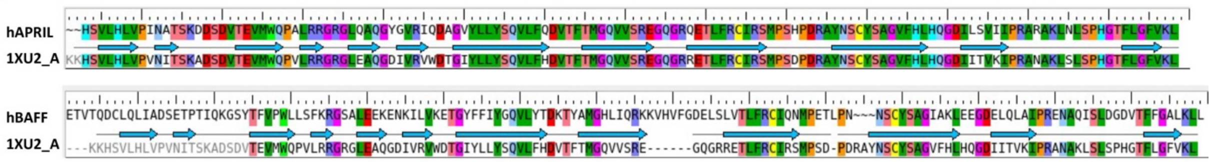


B

| Sequence Alignment | Identities | Positives |
|---|---|---|
| hAPRIL(115-250) vs. 1XU2_A | 86% | 91% |
| hBAFF(140-285) vs. 1XU2_A | 40% | 58% |

C

| Structrual Alignment | Score | RMSD(Å) |
|---|---|---|
| hAPRIL(4ZCH) vs. 1XU2_A | 0.032 | 0.893 |
| hBAFF(4ZCH) vs. 1XU2_A | 0.062 | 1.248 |
| hAPRIL(4ZCH) vs. hBAFF(4ZCH) | 0.057 | 1.189 |

D

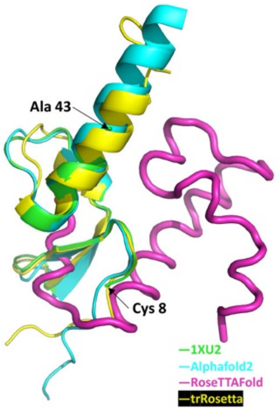


**Sup. Figure 9: (A)** Amino Acid sequence alignment between hAPRIL (top) or hBAFF (bottom) and Protein Data Bank structural ID 1XU2 (structure of mAPRIL and human sBCMA co-complex). PDB No. 4ZCH reported a single-chain human APRIL-BAFF-BAFF heterotrimer structure. **(B)** Quantification of sequence alignment between hAPRIL/hBAFF and published structure 1XU2. **(C)** Quantification of structural alignment between hAPRIL/hBAFF and published structure 1XU2 as well as between predicted hAPRIL and hBAFF structures. **(D)** Structure overlay of extracellular BCMA flexible region (aa no. 44-54) between 1XU2 and structures predicted using Alphafold2 (cyan), RoseTTAfold (magenta) and trRosetta (yellow).

Supplemental Figure 10

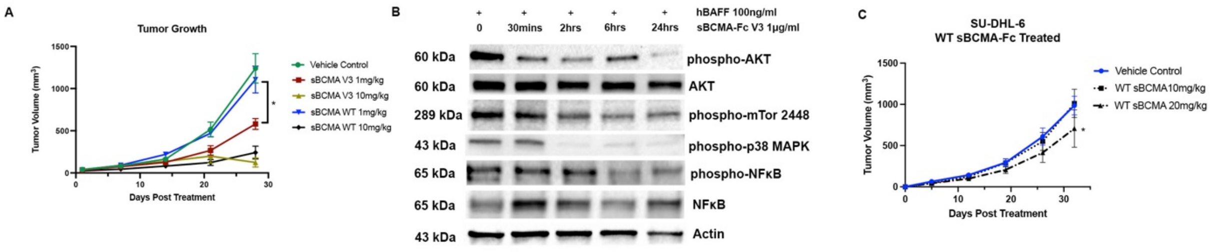


**Sup. Figure 10: (A)** Subcutaneous tumor growth of MM1.R MM tumors in 6 weeks old female NSG mice dosed with wildtype sBCMA-Fc at 1 and 10 mg/kg every 48 hours (N=5), sBCMA-Fc V3 at 1 and 10 mg/kg every 48 hours (N=5) and vehicle control (N=5). **(B)** Analysis of downstream protein expression in SU-DHL-6 DLBCL cells upon sBCMA-Fc V3 treatment at multiple time point. **(C)** Subcutaneous tumor growth of SU-DHL-6 DLBCL tumors in mice dosed with vehicle control, sBCMA-Fc 10 mg/kg and sBCMA-Fc 20 mg/kg every 48 hours (N=5). Statistical analysis was conducted using T test and one way ANOVA for comparing between treatment groups. Repeated ANOVA used for changes in tumor growth over time. P value *=<0.05, **=<0.01. ***=<0.001.
