## Supplemental tables for "Developing High-Affinity Decoy Receptor Optimized for Multiple Myeloma and Diffuse Large B Cell Lymphoma Treatment"

### Supplemental Table 1:

Clones selected from round 3

| Amino acid | | 1 | 2 | 3 | 4 | 5 | 6 | 7 | 8 | 9 | 10 | 11 | 12 | 13 | 14 | 15 | 16 | 17 | 18 | 19 | 20 | 21 | 22 | 23 | 24 | 25 | 26 | 27 | 28 | 29 | 30 | 31 | 32 | 33 | 34 | 35 | 36 | 37 | 38 | 39 | 40 | 41 | 42 | 43 | 44 | 45 | 46 | 47 | 48 | 49 | 50 | 51 | 52 | 53 | 54 | 55 | 56 |
|---|---|---|---|---|---|---|---|---|---|---|---|---|---|---|---|---|---|---|---|---|---|---|---|---|---|---|---|---|---|---|---|---|---|---|---|---|---|---|---|---|---|---|---|---|---|---|---|---|---|---|---|---|---|---|---|---|---|
| WT-BCMA | | M | L | Q | M | A | G | Q | C | S | Q | N | E | Y | F | D | S | L | L | H | A | C | I | P | C | Q | L | R | C | S | S | N | T | P | P | L | T | C | Q | R | Y | C | N | A | S | V | T | N | S | V | K | G | T | N | A | G | S |
| Clone # | 1 | M | S | Q | M | A | G | Q | C | P | Q | N | K | Y | F | D | S | L | L | H | A | C | I | P | C | Q | L | R | C | S | S | D | T | P | P | L | A | C | Q | R | Y | C | S | A | S | V | T | N | S | V | K | G | T | S | A | G | S |
| Clone # | 2 | V | L | Q | M | A | G | Q | C | S | Q | N | E | Y | F | D | S | L | L | H | A | C | I | P | C | Q | L | R | C | S | S | N | P | P | P | L | A | C | Q | R | Y | C | N | A | S | V | I | N | S | V | K | G | T | D | V | G | S |
| Clone # | 3 | M | L | R | M | A | G | Q | C | S | Q | N | E | Y | F | D | N | L | L | H | A | C | I | P | C | Q | L | R | C | S | S | N | T | P | P | L | A | C | Q | R | Y | C | N | T | S | V | T | N | S | V | K | G | T | N | A | G | S |
| Clone # | 4 | M | L | Q | M | A | G | Q | C | S | Q | N | E | Y | L | D | G | L | L | H | A | C | I | P | C | Q | L | R | C | S | S | N | T | P | P | L | A | C | Q | R | Y | C | N | A | S | A | T | D | S | V | K | G | T | N | A | G | S |
| Clone # | 5 | T | L | Q | V | A | G | Q | C | F | Q | N | E | Y | F | D | G | L | L | H | A | C | I | P | C | Q | L | R | C | S | S | N | A | P | P | L | T | C | R | R | Y | C | N | A | S | V | T | N | S | V | K | G | T | N | A | G | S |
| Clone # | 6 | A | L | Q | M | A | G | Q | C | A | Q | N | E | Y | F | D | S | L | L | H | A | C | I | P | C | Q | L | R | C | S | S | N | T | P | P | L | T | C | R | R | Y | C | N | A | S | V | T | N | S | V | K | G | T | N | A | E | S |
| Clone # | 7 | M | L | Q | M | A | E | Q | C | S | Q | N | E | Y | F | D | S | L | L | H | A | C | I | P | C | R | L | R | C | S | S | N | T | P | P | L | T | C | R | R | Y | C | N | A | S | V | T | N | S | V | K | G | T | N | A | G | S |
| Clone # | 8 | V | L | Q | I | A | E | Q | C | P | Q | D | E | Y | F | D | S | L | L | H | A | C | I | P | C | Q | L | R | C | S | S | N | T | P | P | L | T | C | Q | R | Y | C | N | A | S | V | T | N | S | M | K | G | M | N | V | G | S |
| Clone # | 9 | M | L | Q | M | A | G | Q | C | S | Q | D | E | Y | F | D | G | L | L | H | A | C | I | P | C | Q | L | R | C | S | S | S | T | P | P | L | T | C | Q | R | Y | C | N | A | S | V | T | N | S | V | K | G | T | N | A | G | S |
| Clone # | 10 | M | L | Q | M | A | G | Q | C | S | Q | D | E | Y | F | D | S | L | L | Y | A | C | M | P | C | Q | L | R | C | S | S | N | P | P | P | L | T | C | Q | R | Y | C | N | A | S | V | T | S | S | V | K | G | T | S | A | G | S |
| Clone # | 11 | M | L | Q | M | A | E | R | C | S | Q | N | E | Y | F | D | S | L | L | Y | A | C | I | P | C | Q | L | R | C | S | S | N | T | P | P | S | T | C | Q | R | Y | C | N | A | S | V | T | N | S | V | K | G | T | N | A | G | S |
| Clone # | 12 | M | L | Q | M | A | G | Q | C | S | Q | N | E | Y | F | D | S | L | L | Y | A | C | I | P | C | Q | L | R | C | S | S | N | T | P | P | L | T | C | Q | R | Y | C | D | A | S | V | T | N | P | V | K | G | A | N | A | G | S |

**Supplemental Table 2:**

Clones selected from round 4

| Amino acid | | 1 | 2 | 3 | 4 | 5 | 6 | 7 | 8 | 9 | 10 | 11 | 12 | 13 | 14 | 15 | 16 | 17 | 18 | 19 | 20 | 21 | 22 | 23 | 24 | 25 | 26 | 27 | 28 | 29 | 30 | 31 | 32 | 33 | 34 | 35 | 36 | 37 | 38 | 39 | 40 | 41 | 42 | 43 | 44 | 45 | 46 | 47 | 48 | 49 | 50 | 51 | 52 | 53 | 54 | 55 | 56 |
|---|---|---|---|---|---|---|---|---|---|---|---|---|---|---|---|---|---|---|---|---|---|---|---|---|---|---|---|---|---|---|---|---|---|---|---|---|---|---|---|---|---|---|---|---|---|---|---|---|---|---|---|---|---|---|---|---|---|
| **WT -BCMA** | | **M** | **L** | **Q** | **M** | **A** | **G** | **Q** | **C** | **S** | **Q** | **N** | **E** | **Y** | **F** | **D** | **S** | **L** | **L** | **H** | **A** | **C** | **I** | **P** | **C** | **Q** | **L** | **R** | **C** | **S** | **S** | **N** | **T** | **P** | **P** | **L** | **T** | **C** | **Q** | **R** | **Y** | **C** | **N** | **A** | **S** | **V** | **T** | **N** | **S** | **V** | **K** | **G** | **T** | **N** | **A** | **G** | **S** |
| Clone # | **13** | V | L | Q | M | A | G | Q | C | S | Q | N | E | Y | F | D | S | L | L | H | A | C | I | P | C | Q | L | R | C | S | S | D | I | P | P | L | A | C | Q | R | Y | C | N | A | S | V | T | N | S | V | K | G | T | N | A | G | S |
| Clone # | **14** | V | L | Q | M | T | G | Q | C | S | Q | N | E | Y | F | D | S | L | L | L | A | C | I | P | C | Q | L | R | C | S | S | N | T | P | P | L | A | C | Q | R | Y | C | N | A | S | V | T | N | S | V | K | G | T | N | A | G | S |
| Clone # | **15** | T | L | Q | M | A | G | Q | C | S | Q | N | E | Y | F | D | S | L | L | H | A | C | I | P | C | Q | L | R | C | S | S | D | A | P | P | L | A | C | R | R | Y | C | N | A | D | V | T | N | S | A | E | G | T | N | A | G | S |
| Clone # | **16** | V | L | Q | M | A | G | Q | C | S | Q | N | E | Y | F | D | S | L | L | H | A | C | I | P | C | Q | L | R | C | S | S | N | T | P | P | L | A | C | R | R | Y | C | N | V | S | V | T | N | S | V | K | G | T | N | A | G | S |
| Clone # | **17** | V | S | Q | M | A | G | Q | C | P | H | N | E | Y | F | D | S | L | L | H | A | C | I | P | C | Q | L | R | C | S | S | N | T | P | P | L | A | C | R | R | Y | C | N | A | S | V | T | N | S | V | G | G | T | N | A | G | S |
| Clone # | **18** | M | L | Q | M | A | G | Q | C | S | Q | N | E | Y | F | D | S | L | L | H | A | C | I | P | C | Q | L | R | C | S | S | N | T | P | P | L | A | C | R | R | Y | C | N | A | S | V | T | N | S | V | K | G | T | S | A | G | S |
| Clone # | **19** | T | S | Q | M | A | G | Q | C | S | Q | N | E | Y | F | D | S | L | L | H | A | C | I | P | C | Q | L | R | C | S | S | N | T | P | P | P | A | C | R | R | Y | C | N | A | S | V | A | N | S | V | R | G | T | N | A | G | S |
| Clone # | **20** | M | L | Q | M | T | G | Q | C | S | Q | N | E | Y | F | D | S | L | L | H | V | C | I | P | C | Q | L | R | C | S | S | N | T | P | P | L | A | C | R | R | Y | C | N | A | S | V | T | N | S | V | K | G | T | N | A | G | S |
| Clone # | **21** | T | L | Q | M | A | G | Q | C | S | Q | N | E | Y | F | D | G | L | L | H | A | C | V | P | C | Q | L | R | C | S | S | N | T | P | P | L | A | C | Q | R | Y | C | N | A | G | V | A | N | S | A | K | G | T | N | A | G | S |
| Clone # | **22** | M | L | Q | M | A | G | Q | C | S | Q | N | E | Y | F | D | G | L | L | H | A | C | I | P | C | Q | L | R | C | S | S | N | T | P | P | L | A | C | Q | R | Y | C | N | A | S | V | T | N | S | V | K | G | T | N | A | G | S |
| Clone # | **23** | I | L | Q | M | A | G | Q | C | S | Q | D | E | Y | F | D | G | L | L | H | A | C | M | P | C | Q | L | R | C | A | S | N | T | P | P | L | A | C | Q | R | Y | C | N | A | G | V | T | N | S | V | R | G | T | N | A | G | S |
| Clone # | **24** | C | C | R | E | A | G | Q | C | S | Q | D | E | Y | F | D | G | L | L | H | A | C | I | P | C | Q | L | R | C | S | S | N | T | P | P | L | P | C | Q | R | Y | C | N | A | S | V | T | N | S | V | K | G | T | N | A | R | S |
| Clone # | **25** | I | L | Q | M | A | G | Q | C | S | Q | D | E | Y | F | D | G | L | L | H | A | C | M | P | C | Q | L | R | C | A | S | N | T | P | P | L | A | C | Q | R | Y | C | N | A | G | V | T | N | S | V | R | G | T | N | A | G | S |
| Clone # | **26** | M | L | Q | M | A | G | Q | C | S | Q | D | E | Y | F | D | S | L | L | H | A | C | I | P | C | Q | L | R | C | S | S | D | I | P | P | L | A | C | Q | R | Y | C | N | A | N | V | T | D | S | V | K | G | T | D | A | G | S |
| Clone # | **27** | R | C | R | * | A | G | Q | C | S | Q | D | E | Y | F | D | S | L | L | Y | A | C | I | P | C | Q | L | R | C | S | S | N | T | P | P | L | A | C | Q | R | Y | C | S | A | S | A | T | N | S | V | K | G | T | S | A | G | S |
| Clone # | **28** | M | L | Q | M | A | G | Q | C | S | Q | N | E | Y | F | D | S | L | L | Y | A | C | I | P | C | Q | L | R | C | S | S | N | T | P | P | L | A | C | Q | R | Y | C | N | A | G | V | T | N | S | V | K | G | T | N | A | G | S |
| Clone # | **29** | M | L | Q | M | A | G | Q | C | S | Q | N | E | Y | F | D | S | L | L | Y | A | C | I | P | C | Q | L | R | C | S | S | N | I | P | P | L | A | C | Q | R | Y | C | N | A | S | V | T | N | S | A | K | G | T | N | A | G | S |
| Clone # | **30** | M | L | Q | M | A | G | Q | C | S | Q | N | E | Y | F | D | S | L | L | Y | A | C | I | P | C | Q | L | R | C | S | S | S | T | P | P | L | A | C | Q | R | Y | C | N | A | S | A | T | N | S | V | K | G | T | N | A | G | S |
| Clone # | **31** | M | L | Q | M | A | G | Q | C | S | Q | N | E | Y | F | D | S | L | L | Y | A | C | I | P | C | Q | L | R | C | S | S | S | T | P | P | L | A | C | Q | R | Y | C | N | A | S | V | T | N | S | V | K | G | T | N | A | G | S |
| Clone # | **32** | M | L | Q | M | A | G | Q | C | S | Q | N | E | Y | F | D | S | L | L | Y | A | C | I | P | C | Q | L | R | C | S | S | N | T | P | P | L | P | C | Q | R | Y | C | N | A | S | V | T | N | S | V | K | G | A | N | A | G | S |
| Clone # | **33** | M | L | Q | M | A | G | Q | C | S | Q | N | E | Y | F | D | S | L | L | Y | A | C | I | P | C | Q | L | R | C | S | S | D | T | P | P | L | T | C | Q | R | Y | C | N | A | S | V | T | N | S | V | K | G | M | N | A | G | S |
| Clone # | **34** | V | L | Q | M | A | G | Q | C | S | Q | N | E | Y | F | D | S | L | L | Y | A | C | I | P | C | Q | L | R | C | S | S | N | T | P | P | L | T | C | Q | R | Y | C | N | A | S | M | T | N | S | V | K | G | T | N | A | G | S |
| Clone # | **35** | M | L | Q | M | A | G | Q | C | S | Q | N | E | Y | F | D | G | L | L | Y | A | C | I | P | C | Q | L | R | C | S | S | N | T | P | P | L | T | C | Q | R | Y | C | N | A | S | V | T | D | S | V | K | G | T | N | A | G | S |
| Clone # | **36** | M | L | Q | M | A | G | Q | C | S | Q | N | E | Y | F | D | G | L | L | Y | A | C | I | P | C | Q | L | R | C | S | S | N | T | P | P | L | T | C | Q | R | Y | C | N | A | S | V | T | N | S | V | T | G | T | N | A | G | S |
| Clone # | **37** | M | L | Q | M | A | G | Q | C | S | Q | N | E | Y | F | D | G | L | L | Y | A | C | I | P | C | Q | L | R | C | S | S | N | T | P | P | L | T | C | Q | R | Y | C | N | A | N | V | T | N | S | V | R | G | T | N | A | G | S |
| Clone # | **38** | M | L | Q | M | A | G | Q | C | S | Q | D | E | Y | F | D | S | L | L | Y | A | C | I | P | C | Q | L | R | C | S | S | N | T | P | P | L | T | C | Q | R | Y | C | N | A | S | V | T | N | T | V | K | G | T | N | A | G | S |
| Clone # | **39** | M | L | Q | M | A | G | Q | C | P | Q | D | E | Y | F | D | R | L | L | H | A | C | I | P | C | Q | L | R | C | S | S | N | A | P | P | L | T | C | R | R | Y | C | N | A | G | V | I | N | S | V | K | G | A | D | T | G | S |
| Clone # | **40** | M | L | Q | M | A | G | Q | C | S | Q | D | E | Y | F | D | G | L | L | H | A | C | I | P | C | Q | L | R | C | S | S | N | T | P | P | L | T | C | Q | R | Y | C | N | A | R | V | T | N | S | V | K | G | T | N | A | G | S |
| Clone # | **41** | M | L | Q | M | A | G | Q | C | S | Q | N | E | Y | F | D | S | L | L | L | A | C | I | P | C | Q | L | R | C | S | S | N | A | P | P | L | T | C | Q | R | Y | C | N | A | G | V | T | N | S | V | K | E | A | N | A | G | S |

**Supplemental Table 3:**

Clones selected from round 5

| Amino acid | | 1 | 2 | 3 | 4 | 5 | 6 | 7 | 8 | 9 | 10 | 11 | 12 | 13 | 14 | 15 | 16 | 17 | 18 | 19 | 20 | 21 | 22 | 23 | 24 | 25 | 26 | 27 | 28 | 29 | 30 | 31 | 32 | 33 | 34 | 35 | 36 | 37 | 38 | 39 | 40 | 41 | 42 | 43 | 44 | 45 | 46 | 47 | 48 | 49 | 50 | 51 | 52 | 53 | 54 | 55 | 56 |
|---|---|---|---|---|---|---|---|---|---|---|---|---|---|---|---|---|---|---|---|---|---|---|---|---|---|---|---|---|---|---|---|---|---|---|---|---|---|---|---|---|---|---|---|---|---|---|---|---|---|---|---|---|---|---|---|---|---|
| WT-BCMA | | M | L | Q | M | A | G | Q | C | S | Q | N | E | Y | F | D | S | L | L | H | A | C | I | P | C | Q | L | R | C | S | S | N | T | P | P | L | T | C | Q | R | Y | C | N | A | S | V | T | N | S | V | K | G | T | N | A | G | S |
| Clone # | 42 | M | L | Q | M | A | G | Q | C | S | Q | N | E | Y | F | D | N | L | L | Y | A | C | I | P | C | Q | L | R | C | S | S | N | T | P | P | L | A | C | Q | R | Y | C | N | A | S | V | T | N | S | V | R | G | T | N | A | R | S |
| Clone # | 43 | V | L | Q | M | A | G | Q | C | S | Q | N | E | Y | F | D | S | L | L | Y | A | C | I | P | C | Q | L | R | C | S | S | N | T | P | P | L | A | C | Q | H | Y | C | N | A | S | V | A | N | S | V | K | G | T | N | A | G | S |
| Clone # | 44 | V | L | Q | M | A | G | Q | C | S | Q | N | E | Y | F | D | S | L | L | Y | A | C | I | P | C | Q | L | R | C | S | S | N | T | P | P | L | A | C | Q | R | Y | C | N | A | S | V | T | N | S | V | K | G | T | N | A | G | S |
| Clone # | 45 | M | L | Q | M | A | G | Q | C | S | Q | N | E | Y | F | D | S | L | L | Y | A | C | I | P | C | Q | L | R | C | S | S | N | T | P | P | L | A | C | Q | R | Y | C | D | A | S | V | T | S | P | V | K | G | T | N | A | R | S |
| Clone # | 46 | V | L | Q | M | A | G | Q | C | S | Q | N | E | Y | F | D | S | L | L | Y | A | C | I | P | C | Q | L | R | C | S | S | N | T | P | P | L | A | C | Q | R | Y | C | N | A | G | V | T | D | S | V | K | G | T | N | A | G | S |
| Clone # | 47 | V | L | Q | M | A | G | Q | C | S | Q | N | E | Y | F | D | S | L | L | Y | A | C | I | P | C | Q | L | R | C | S | S | N | T | P | P | L | A | C | Q | R | Y | C | R | A | S | V | T | N | S | V | K | G | T | S | A | G | S |
| Clone # | 48 | M | L | Q | M | A | G | Q | C | S | Q | N | E | Y | F | D | S | L | L | Y | A | C | I | P | C | Q | L | R | C | S | S | N | T | P | P | P | A | C | Q | R | Y | C | D | A | S | V | I | N | S | A | K | G | T | N | A | G | S |
| Clone # | 49 | M | L | P | M | A | G | Q | C | P | Q | N | E | Y | F | D | S | L | L | Y | A | C | I | P | C | Q | L | R | C | S | S | S | T | P | P | L | A | C | Q | H | Y | C | N | A | S | V | T | R | S | V | E | G | T | N | A | G | S |
| Clone # | 50 | V | L | Q | M | A | G | Q | C | S | Q | N | E | Y | F | D | S | L | L | Y | A | C | I | P | C | Q | L | R | C | S | S | N | T | P | P | L | A | C | Q | R | Y | C | R | A | S | V | T | N | S | V | K | G | T | S | A | G | S |
| Clone # | 51 | T | L | Q | M | A | G | Q | C | S | Q | N | E | Y | F | D | S | L | L | Y | A | C | I | P | C | Q | L | R | C | S | S | N | T | P | P | L | A | C | Q | R | Y | C | N | A | S | V | T | N | S | V | K | G | T | N | A | G | S |
| Clone # | 52 | V | L | Q | M | A | G | Q | C | S | Q | N | E | Y | F | D | N | L | L | Y | A | C | M | P | C | Q | L | R | C | S | S | N | T | P | P | L | A | C | Q | R | Y | C | N | A | S | V | T | N | S | V | K | G | T | N | A | G | S |
| Clone # | 53 | T | L | Q | M | A | G | Q | C | S | Q | D | E | Y | F | D | S | L | L | Y | A | C | I | P | C | Q | L | R | C | S | S | N | T | P | P | L | A | C | Q | R | Y | C | S | A | S | A | T | N | S | V | K | G | T | S | A | G | S |
| Clone # | 54 | T | L | Q | M | A | G | Q | C | S | Q | D | E | Y | F | D | S | L | L | Y | A | C | I | P | C | Q | L | R | C | S | S | N | T | P | P | L | A | C | Q | R | Y | C | S | A | S | A | T | N | S | V | K | G | T | S | A | G | S |
| Clone # | 55 | M | L | Q | M | A | G | Q | C | S | Q | D | E | Y | F | D | G | L | L | Y | A | C | I | P | C | Q | L | R | C | S | S | N | T | P | P | L | A | C | Q | R | Y | C | N | A | S | V | T | S | S | V | K | G | T | D | A | G | S |
| Clone # | 56 | M | L | Q | M | A | G | Q | C | S | Q | D | E | Y | F | D | G | L | L | Y | A | C | I | P | C | Q | L | R | C | S | S | N | T | P | P | L | A | C | Q | R | Y | C | N | A | S | V | T | S | S | V | K | G | T | D | A | G | S |
| Clone # | 57 | V | L | Q | M | A | G | Q | C | P | P | N | E | Y | F | D | G | L | L | Y | A | C | I | P | C | Q | F | R | C | S | S | N | T | P | P | L | A | C | Q | R | Y | C | N | V | S | V | T | N | S | V | K | G | T | D | A | G | S |
| Clone # | 58 | M | L | Q | M | A | G | Q | C | S | Q | N | E | Y | F | D | G | L | L | Y | A | C | I | P | C | Q | L | R | C | S | S | N | T | P | P | L | A | C | Q | R | Y | C | N | A | S | V | T | N | S | A | K | G | T | D | A | G | S |
| Clone # | 59 | M | L | Q | M | A | G | Q | C | S | Q | N | E | Y | F | D | G | L | L | H | A | C | I | P | C | Q | L | R | C | S | S | N | T | P | P | L | A | C | Q | R | Y | C | N | T | G | M | T | N | S | V | K | G | T | N | A | G | S |
| Clone # | 60 | M | L | Q | V | A | G | Q | C | P | Q | N | E | Y | F | D | G | L | L | H | A | C | I | P | C | Q | L | R | C | S | S | N | T | P | P | L | A | C | R | R | Y | C | N | A | S | V | T | N | S | V | K | G | T | N | A | G | S |
| Clone # | 61 | M | L | Q | M | A | G | Q | C | P | Q | S | E | Y | F | D | G | L | L | H | A | C | I | P | C | Q | L | R | C | S | S | N | T | P | P | L | A | C | R | R | Y | C | N | A | S | V | T | N | S | V | K | G | T | N | A | G | S |
| Clone # | 62 | M | L | Q | M | A | G | Q | C | S | Q | D | K | Y | F | D | R | L | L | H | A | C | I | P | C | Q | L | R | C | S | S | N | T | P | P | L | A | C | Q | R | Y | C | N | A | S | V | T | N | S | V | K | G | M | N | A | G | S |
| Clone # | 63 | M | L | Q | V | A | G | Q | C | S | Q | N | E | Y | F | D | S | L | L | H | A | C | I | P | C | Q | L | R | C | S | S | N | I | P | P | L | A | C | R | R | Y | C | N | T | S | A | T | N | P | V | K | G | T | N | A | G | S |
| Clone # | 64 | M | L | Q | M | A | G | Q | C | P | Q | D | E | Y | F | D | G | L | L | H | A | C | I | P | C | R | L | R | C | S | S | N | T | P | P | L | T | C | Q | R | Y | C | N | A | S | V | T | N | S | V | K | G | T | N | A | G | S |
| Clone # | 65 | T | L | Q | M | T | G | Q | C | P | Q | N | E | Y | F | D | G | L | L | H | A | C | I | P | C | R | L | R | C | S | S | D | T | P | P | L | T | C | Q | R | Y | C | N | A | S | V | T | N | S | M | K | G | T | N | A | G | S |
| Clone # | 66 | M | S | Q | M | A | G | Q | C | P | Q | N | E | Y | F | D | G | L | L | H | T | C | I | P | C | Q | L | R | C | S | S | N | I | P | P | L | T | C | R | R | Y | C | D | A | S | V | A | N | L | V | K | G | T | N | A | G | S |
| Clone # | 67 | M | L | Q | M | A | G | Q | C | S | Q | N | E | Y | F | D | G | L | L | H | A | C | I | P | C | R | L | R | C | S | S | N | T | P | P | L | T | C | Q | R | Y | C | N | A | S | V | A | N | S | V | K | G | T | N | A | G | S |
| Clone # | 68 | M | L | Q | M | A | E | Q | C | A | Q | N | E | Y | F | D | G | L | L | H | A | C | I | P | C | R | L | R | C | S | S | D | T | P | P | L | T | C | Q | R | Y | C | N | A | S | V | T | S | S | V | K | G | M | N | A | G | S |
| Clone # | 69 | M | L | Q | M | A | G | Q | C | S | Q | N | E | Y | F | D | S | L | L | Y | A | C | I | P | C | Q | L | R | C | S | S | N | T | P | P | L | T | C | R | R | Y | C | N | A | S | V | T | N | S | V | K | G | M | N | A | G | S |
| Clone # | 70 | M | L | Q | M | A | G | Q | C | S | Q | D | E | Y | F | D | S | L | L | Y | A | C | M | P | C | Q | L | R | C | S | S | N | P | P | P | L | T | C | Q | R | Y | C | N | A | S | V | T | S | S | V | K | G | T | S | A | G | S |
| Clone # | 71 | M | L | Q | M | A | G | Q | C | S | Q | N | E | Y | F | D | G | L | L | Y | A | C | I | P | C | Q | L | R | C | S | S | N | T | P | P | L | A | C | Q | R | Y | C | N | A | S | V | T | N | S | V | K | G | T | N | A | G | S |
| Clone # | 72 | M | L | Q | M | A | G | Q | C | S | Q | N | E | Y | F | D | G | L | L | Y | A | C | I | P | C | Q | L | R | C | S | S | N | T | P | P | L | A | C | Q | R | Y | C | N | A | S | V | T | N | S | V | K | G | T | D | A | G | S |
| Clone # | 73 | M | L | Q | M | A | G | Q | C | S | Q | N | E | Y | F | D | G | L | L | H | A | C | I | P | C | Q | L | R | C | S | S | N | T | P | P | L | A | C | Q | R | Y | C | N | A | S | V | T | N | S | V | K | G | T | N | A | G | S |
| Clone # | 74 | M | L | Q | M | A | G | Q | C | P | Q | D | E | Y | F | D | G | L | L | Y | A | C | I | P | C | Q | L | R | C | S | S | N | T | P | P | L | A | C | Q | R | Y | C | N | A | S | V | T | S | S | V | K | G | T | D | A | G | S |

**Supplemental Table 4:**

Clones selected from round 6

| Amino acid | | 1 | 2 | 3 | 4 | 5 | 6 | 7 | 8 | 9 | 10 | 11 | 12 | 13 | 14 | 15 | 16 | 17 | 18 | 19 | 20 | 21 | 22 | 23 | 24 | 25 | 26 | 27 | 28 | 29 | 30 | 31 | 32 | 33 | 34 | 35 | 36 | 37 | 38 | 39 | 40 | 41 | 42 | 43 | 44 | 45 | 46 | 47 | 48 | 49 | 50 | 51 | 52 | 53 | 54 | 55 | 56 |
|---|---|---|---|---|---|---|---|---|---|---|---|---|---|---|---|---|---|---|---|---|---|---|---|---|---|---|---|---|---|---|---|---|---|---|---|---|---|---|---|---|---|---|---|---|---|---|---|---|---|---|---|---|---|---|---|---|---|
| WT-BCMA | | M | L | Q | M | A | G | Q | C | S | Q | N | E | Y | F | D | S | L | L | H | A | C | I | P | C | Q | L | R | C | S | S | N | T | P | P | L | T | C | Q | R | Y | C | N | A | S | V | T | N | S | V | K | G | T | N | A | G | S |
| Clone # | 75 | M | L | P | M | A | G | Q | C | P | Q | N | E | Y | F | D | S | L | L | Y | A | C | I | P | C | Q | L | R | C | S | S | S | T | P | P | L | A | C | Q | H | Y | C | N | A | S | V | T | R | S | V | E | G | T | N | A | G | S |
| Clone # | 76 | M | L | P | M | A | G | Q | C | P | Q | N | E | Y | F | D | S | L | L | Y | A | C | I | P | C | Q | L | R | C | S | S | S | T | P | P | L | A | C | Q | H | Y | C | N | A | S | V | T | R | S | V | E | G | T | N | A | G | S |
| Clone # | 77 | M | L | P | M | A | G | Q | C | P | Q | N | E | Y | F | D | S | L | L | Y | A | C | I | P | C | Q | L | R | C | S | S | S | T | P | P | L | A | C | Q | H | Y | C | N | A | S | V | T | R | S | V | E | G | T | N | A | G | S |
| Clone # | 78 | M | L | P | M | A | G | Q | C | P | Q | N | E | Y | F | D | S | L | L | Y | A | C | I | P | C | Q | L | R | C | S | S | S | T | P | P | L | A | C | Q | H | Y | C | N | A | S | V | T | R | S | V | E | G | T | N | A | G | S |
| Clone # | 79 | M | L | P | M | A | G | Q | C | P | Q | N | E | Y | F | D | S | L | L | Y | A | C | I | P | C | Q | L | R | C | S | S | S | T | P | P | L | A | C | Q | H | Y | C | N | A | S | V | T | R | S | V | E | G | T | N | A | G | S |
| Clone # | 80 | M | L | P | M | A | G | Q | C | P | Q | N | E | Y | F | D | S | L | L | Y | A | C | I | P | C | Q | L | R | C | S | S | S | T | P | P | L | A | C | Q | H | Y | C | N | A | S | V | T | R | S | V | E | G | T | N | A | G | S |
| Clone # | 81 | M | L | P | M | A | G | Q | C | P | Q | N | E | Y | F | D | S | L | L | Y | A | C | I | P | C | Q | L | R | C | S | S | S | T | P | P | L | A | C | Q | H | Y | C | N | A | S | V | T | R | S | V | E | G | T | N | A | G | S |
| Clone # | 82 | M | L | P | M | A | G | Q | C | P | Q | N | E | Y | F | D | S | L | L | Y | A | C | I | P | C | Q | L | R | C | S | S | S | T | P | P | L | A | C | Q | H | Y | C | N | A | S | V | T | R | S | V | E | G | T | N | A | G | S |
| Clone # | 83 | M | L | P | M | A | G | Q | C | P | Q | N | E | Y | F | D | S | L | L | Y | A | C | I | P | C | Q | L | R | C | S | S | S | T | P | P | L | A | C | Q | H | Y | C | N | A | S | V | T | R | S | V | E | G | T | N | A | G | S |
| Clone # | 84 | M | L | P | M | A | G | Q | C | P | Q | N | E | Y | F | D | S | L | L | Y | A | C | I | P | C | Q | L | R | C | S | S | S | T | P | P | L | A | C | Q | H | Y | C | N | A | S | V | T | R | S | V | E | G | T | N | A | G | S |
| Clone # | 85 | M | L | P | M | A | G | Q | C | P | Q | N | E | Y | F | D | S | L | L | Y | A | C | I | P | C | Q | L | R | C | S | S | S | T | P | P | L | A | C | Q | H | Y | C | N | A | S | V | T | R | S | V | E | G | T | N | A | G | S |
| Clone # | 86 | M | L | P | M | A | G | Q | C | P | Q | N | E | Y | F | D | S | L | L | Y | A | C | I | P | C | Q | L | R | C | S | S | S | T | P | P | L | A | C | Q | H | Y | C | N | A | S | V | T | R | S | V | E | G | T | N | A | G | S |
| Clone # | **87** | M | L | P | M | A | G | Q | C | P | Q | N | E | Y | F | D | S | L | L | Y | A | C | I | P | C | Q | L | R | C | S | S | S | T | P | P | L | A | C | Q | H | Y | C | N | A | S | V | T | R | S | V | E | G | T | N | A | G | S |
| Clone # | **88** | M | L | P | M | A | G | Q | C | P | Q | N | E | Y | F | D | S | L | L | Y | A | C | I | P | C | Q | L | R | C | S | S | S | T | P | P | L | A | C | Q | H | Y | C | N | A | S | V | T | R | S | V | E | G | T | N | A | G | S |
| Clone # | **89** | M | L | P | M | A | G | Q | C | P | Q | N | E | Y | F | D | S | L | L | Y | A | C | I | P | C | Q | L | R | C | S | S | S | T | P | P | L | A | C | Q | H | Y | C | N | A | S | V | T | R | S | V | E | G | T | N | A | G | S |
| Clone # | **90** | M | L | P | M | A | G | Q | C | P | Q | N | E | Y | F | D | S | L | L | Y | A | C | I | P | C | Q | L | R | C | S | S | S | T | P | P | L | A | C | Q | H | Y | C | N | A | S | V | T | R | S | V | E | G | T | N | A | G | S |
| Clone # | **91** | M | L | P | M | A | G | Q | C | P | Q | N | E | Y | F | D | S | L | L | Y | A | C | I | P | C | Q | L | R | C | S | S | S | T | P | P | L | A | C | Q | H | Y | C | N | A | S | V | T | R | S | V | E | G | T | N | A | G | S |
| Clone # | **92** | M | L | P | M | A | G | Q | C | P | Q | N | E | Y | F | D | S | L | L | Y | A | C | I | P | C | Q | L | R | C | S | S | S | T | P | P | L | A | C | Q | H | Y | C | N | A | S | V | T | R | S | V | E | G | T | N | A | G | S |
| Clone # | **93** | M | L | P | M | A | G | Q | C | P | Q | N | E | Y | F | D | S | L | L | Y | A | C | I | P | C | Q | L | R | C | S | S | S | T | P | P | L | A | C | Q | H | Y | C | N | A | S | V | T | R | S | V | E | G | T | N | A | G | S |
| Clone # | **94** | M | L | P | M | A | G | Q | C | P | Q | N | E | Y | F | D | S | L | L | Y | A | C | I | P | C | Q | L | R | C | S | S | S | T | P | P | L | A | C | Q | H | Y | C | N | A | S | V | T | R | S | V | E | G | T | N | A | G | S |
| Clone # | **95** | V | S | Q | T | A | G | Q | C | S | Q | D | E | Y | F | D | S | L | L | Y | A | C | I | P | C | Q | L | R | C | S | S | N | T | P | P | L | A | C | Q | R | Y | C | N | A | S | V | T | N | S | V | K | G | T | N | A | G | S |
| Clone # | **96** | V | S | Q | T | A | G | Q | C | S | Q | D | E | Y | F | D | S | L | L | Y | A | C | I | P | C | Q | L | R | C | S | S | N | T | P | P | L | A | C | Q | R | Y | C | N | A | S | V | T | N | S | V | K | G | T | N | A | G | S |
| Clone # | **97** | V | S | Q | T | A | G | Q | C | S | Q | D | E | Y | F | D | S | L | L | Y | A | C | I | P | C | Q | L | R | C | S | S | N | T | P | P | L | A | C | Q | R | Y | C | N | A | S | V | T | N | S | V | K | G | T | N | A | G | S |
| Clone # | **98** | V | S | Q | T | A | G | Q | C | S | Q | D | E | Y | F | D | S | L | L | Y | A | C | I | P | C | Q | L | R | C | S | S | N | T | P | P | L | A | C | Q | R | Y | C | N | A | S | V | T | N | S | V | K | G | T | N | A | G | S |
| Clone # | **99** | V | S | Q | T | A | G | Q | C | S | Q | D | E | Y | F | D | S | L | L | Y | A | C | I | P | C | Q | L | R | C | S | S | N | T | P | P | L | A | C | Q | R | Y | C | N | A | S | V | T | N | S | V | K | G | T | N | A | G | S |
| Clone # | **100** | V | S | Q | T | A | G | Q | C | S | Q | N | E | Y | F | D | S | L | L | Y | A | C | I | P | C | Q | L | R | C | S | S | N | T | P | P | L | I | C | Q | R | Y | C | N | A | S | A | T | N | S | M | K | G | T | N | A | G | S |
| Clone # | **101** | V | S | Q | T | A | G | Q | C | S | Q | D | E | Y | F | D | S | L | L | Y | A | C | I | P | C | Q | L | R | C | S | S | N | T | P | P | L | A | C | Q | R | Y | C | N | A | S | V | T | N | S | V | K | G | T | N | A | G | S |
| Clone # | **102** | V | S | Q | T | A | G | Q | C | S | Q | D | E | Y | F | D | S | L | L | Y | A | C | I | P | C | Q | L | R | C | S | S | N | T | P | P | L | A | C | Q | R | Y | C | N | A | S | V | T | N | S | V | K | G | T | N | A | G | S |
| Clone # | **103** | V | S | Q | T | A | G | Q | C | S | Q | D | E | Y | F | D | S | L | L | Y | A | C | I | P | C | Q | L | R | C | S | S | N | T | P | P | L | A | C | Q | R | Y | C | N | A | S | V | T | N | S | V | K | G | T | N | A | G | S |
| Clone # | **104** | V | S | Q | T | A | G | Q | C | P | R | N | E | Y | F | D | S | L | L | Y | A | C | I | P | C | Q | L | R | C | S | S | N | T | P | P | L | A | C | Q | R | Y | C | N | A | S | V | A | S | S | V | K | G | T | N | A | G | S |
| Clone # | **105** | V | S | Q | T | A | G | Q | C | P | R | N | E | Y | F | D | S | L | L | Y | A | C | I | P | C | Q | L | R | C | S | S | N | T | P | P | L | A | C | Q | R | Y | C | N | A | S | V | A | S | S | V | K | G | T | N | A | G | S |
| Clone # | **106** | V | S | Q | T | A | G | Q | C | S | Q | N | E | Y | F | D | G | L | L | H | A | C | I | P | C | Q | L | R | C | S | S | D | I | P | P | L | A | C | Q | R | Y | C | N | A | S | V | T | N | S | V | K | G | T | N | A | G | S |
| Clone # | **107** | V | S | Q | T | A | G | Q | C | S | Q | N | E | Y | F | D | G | L | L | H | A | C | I | P | C | Q | L | R | C | S | S | D | I | P | P | L | A | C | Q | R | Y | C | N | A | S | V | T | N | S | V | K | G | T | N | A | G | S |
| Clone # | **108** | V | S | Q | T | A | G | Q | C | S | Q | N | E | Y | F | D | G | L | L | H | A | C | I | P | C | Q | L | R | C | S | S | D | I | P | P | L | A | C | Q | R | Y | C | N | A | S | V | T | N | S | V | K | G | T | N | A | G | S |
| Clone # | **109** | V | S | Q | T | A | G | Q | C | S | Q | N | E | Y | F | D | G | L | L | H | A | C | I | P | C | Q | L | R | C | S | S | D | I | P | P | L | A | C | Q | R | Y | C | N | A | S | V | T | N | S | V | K | G | T | N | A | G | S |
| Clone # | **110** | V | S | Q | T | A | G | Q | C | S | Q | N | E | Y | F | D | G | L | L | H | A | C | I | P | C | Q | L | R | C | S | S | D | I | P | P | L | A | C | Q | R | Y | C | N | A | S | V | T | N | S | V | K | G | T | N | A | G | S |
| Clone # | **111** | V | S | Q | T | A | G | Q | C | S | Q | N | E | Y | F | D | G | L | L | H | A | C | I | P | C | Q | L | R | C | S | S | D | I | P | P | L | A | C | Q | R | Y | C | N | A | S | V | T | N | S | V | K | G | T | N | A | G | S |
| Clone # | **112** | V | L | Q | T | A | G | Q | C | S | Q | N | E | Y | F | D | S | L | L | H | A | C | I | P | C | Q | L | R | C | S | S | N | T | P | P | L | A | C | R | R | Y | C | N | A | S | V | T | N | S | V | K | G | T | K | A | G | S |
| Clone # | **113** | T | L | Q | M | A | G | Q | C | S | Q | D | E | Y | F | D | S | L | L | Y | A | C | I | P | C | Q | L | R | C | S | S | N | T | P | P | L | A | C | Q | R | Y | C | S | A | S | A | T | N | S | V | K | G | T | S | A | G | S |
| Clone # | **114** | T | L | Q | M | A | G | Q | C | S | Q | N | E | Y | F | D | S | L | L | H | A | C | I | P | C | Q | L | R | C | S | S | D | A | P | P | L | A | C | R | R | Y | C | N | A | D | V | T | N | S | A | E | G | T | N | A | G | S |
| Clone # | **115** | T | L | Q | M | A | G | Q | C | P | Q | N | E | Y | F | D | S | L | L | H | A | C | I | S | C | Q | L | R | C | S | S | N | T | P | P | L | A | C | R | R | Y | C | S | A | S | V | T | N | P | A | K | G | T | N | V | G | S |
| Clone # | **116** | M | L | Q | M | A | G | Q | C | S | Q | N | E | Y | F | D | S | L | L | Y | A | C | I | P | C | Q | L | R | C | S | S | N | T | P | P | L | A | C | Q | R | Y | C | N | A | G | V | T | N | S | V | K | G | T | N | A | G | S |
| Clone # | **117** | M | L | Q | M | A | G | Q | C | S | Q | N | E | Y | F | D | S | L | L | Y | A | C | I | P | C | Q | L | R | C | S | S | N | T | P | P | L | A | C | Q | R | Y | C | N | A | S | V | T | N | S | V | K | G | T | N | A | G | S |
| Clone # | **118** | M | L | Q | T | A | G | Q | C | S | Q | N | E | Y | F | D | S | L | L | H | A | C | I | P | C | Q | L | R | C | S | S | N | T | P | P | L | A | C | R | R | Y | C | S | A | G | V | A | K | P | V | K | G | A | N | A | G | S |

**Supplemental Table 5:**

Male Hematology Results Part I

| Vehicle Control | Day(s) Relative to Start Date | RBC (10^12/L) | HGB (g/L) | HCT (%) | MCV (fL) | MCH (pg) | MCHC (g/L) | RDW (%) | RET (10^9/L) |
|---|---|---|---|---|---|---|---|---|---|
| 1101 | -13 | 6.21 | 139 | 49.4 | 79.6 | 22.4 | 281 | 13.3 | 192.0 |
| | -6 | 5.89 | 133 | 47.4 | 80.6 | 22.6 | 280 | 13.1 | 121.0 |
| | 1 | 5.47 | 124 | 41.7 | 76.3 | 22.7 | 298 | 13.3 | 154.9 |
| | 2 | 5.29 | 119 | 43.7 | 82.5 | 22.5 | 273 | 13.4 | 198.3 |
| | 7 | 4.99 | 112 | 38.9 | 77.9 | 22.4 | 287 | 13.6 | 235.8 |
| | 14 | 5.42 | 124 | 40.5 | 74.6 | 22.8 | 305 | 13.3 | 169.3 |
| | 42 | 5.81 | 129 | 42.6 | 73.4 | 22.3 | 303 | 13.4 | 109.6 |

| AB001 0.1 mg/kg | Day(s) Relative to Start Date | RBC (10^12/L) | HGB (g/L) | HCT (%) | MCV (fL) | MCH (pg) | MCHC (g/L) | RDW (%) | RET (10^9/L) |
|---|---|---|---|---|---|---|---|---|---|
| 1203 | -13 | 5.61 | 126 | 43.5 | 77.6 | 22.4 | 289 | 13.5 | 209.8 |
| | -6 | 6.23 | 140 | 47.9 | 76.8 | 22.5 | 293 | 12.9 | 140.6 |
| | 1 | 5.61 | 126 | 42.6 | 75.9 | 22.5 | 297 | 12.7 | 136.3 |
| | 2 | 5.12 | 114 | 39.0 | 76.2 | 22.3 | 292 | 12.9 | 157.0 |
| | 7 | 5.18 | 116 | 38.6 | 74.6 | 22.4 | 301 | 13.1 | 236.5 |
| | 14 | 5.55 | 122 | 40.2 | 72.4 | 22.0 | 304 | 12.9 | 178.4 |
| | 42 | 6.14 | 139 | 44.4 | 72.3 | 22.6 | 313 | 12.1 | 89.2 |

| AB001 1 mg/kg | Day(s) Relative to Start Date | RBC (10^12/L) | HGB (g/L) | HCT (%) | MCV (fL) | MCH (pg) | MCHC (g/L) | RDW (%) | RET (10^9/L) |
|---|---|---|---|---|---|---|---|---|---|
| 1305 | -13 | 6.18 | 137 | 45.6 | 73.8 | 22.1 | 300 | 12.9 | 250.0 |
| | -6 | 6.55 | 147 | 47.7 | 72.9 | 22.5 | 309 | 12.3 | 160.5 |
| | -3 | 6.69 | 151 | 47.9 | 71.6 | 22.5 | 315 | 12.2 | 137.3 |
| | 1 | 6.37 | 142 | 45.0 | 70.6 | 22.3 | 316 | 12.2 | 123.4 |
| | 2 | 5.67 | 127 | 41.3 | 72.8 | 22.4 | 307 | 12.4 | 144.8 |
| | 7 | 5.71 | 128 | 40.2 | 70.3 | 22.3 | 317 | 12.5 | 172.0 |
| | 14 | 6.08 | 132 | 42.7 | 70.2 | 21.8 | 310 | 12.2 | 188.3 |
| | 42 | 6.26 | 136 | 42.4 | 67.7 | 21.7 | 320 | 11.9 | 95.1 |

| AB001 10 mg/kg | Day(s) Relative to Start Date | RBC (10^12/L) | HGB (g/L) | HCT (%) | MCV (fL) | MCH (pg) | MCHC (g/L) | RDW (%) | RET (10^9/L) |
|---|---|---|---|---|---|---|---|---|---|
| 1407 | -13 | 5.00 | 115 | 38.1 | 76.1 | 22.9 | 301 | 13.8 | 232.6 |
| | -6 | 5.55 | 128 | 43.9 | 79.1 | 23.1 | 292 | 13.1 | 149.8 |
| | -3 | 5.27 | 121 | 39.1 | 74.3 | 22.9 | 308 | 12.7 | 170.4 |

| | | | | | | | | | |
|---|---|---|---|---|---|---|---|---|---|
| | 1 | 5.53 | 127 | 41.3 | 74.7 | 23.0 | 308 | 12.8 | 192.1 |
| | 2 | 5.02 | 114 | 37.9 | 75.5 | 22.7 | 301 | 13.1 | 193.8 |
| | 7 | 4.94 | 112 | 36.7 | 74.3 | 22.6 | 304 | 13.1 | 272.2 |
| | 14 | 5.22 | 118 | 38.9 | 74.5 | 22.5 | 302 | 12.9 | 250.5 |
| | 42 | 5.46 | 124 | 40.7 | 74.6 | 22.7 | 305 | 12.3 | 112.3 |

| AB001 100 mg/kg | Day(s) Relative to Start Date | RBC (10^12/L) | HGB (g/L) | HCT (%) | MCV (fL) | MCH (pg) | MCHC (g/L) | RDW (%) | RET (10^9/L) |
|---|---|---|---|---|---|---|---|---|---|
| 1509 | -13 | 6.12 | 137 | 46.9 | 76.5 | 22.3 | 291 | 12.6 | 100.6 |
| | -6 | 6.21 | 136 | 45.9 | 73.9 | 21.8 | 295 | 12.7 | 116.1 |
| | 1 | 5.89 | 131 | 44.0 | 74.8 | 22.3 | 298 | 12.9 | 152.1 |
| | 2 | 5.56 | 121 | 42.6 | 76.6 | 21.7 | 284 | 13.1 | 141.4 |
| | 7 | 5.32 | 115 | 39.3 | 73.8 | 21.6 | 292 | 13.2 | 258.1 |
| | 14 | 5.94 | 128 | 46.4 | 78.1 | 21.6 | 276 | 12.9 | 220.7 |
| | 42 | 6.71 | 147 | 51.4 | 76.7 | 21.9 | 285 | 12.6 | 88.2 |

Abbreviations:

| | |
|---|---|
| RBC | Red Blood Cells |
| HGB | Hemoglobin |
| HCT | Hematocrit |
| MCV | Mean Corpuscular Volume |
| MCH | Mean Corpuscular Hemoglobin |
| MCHC | Mean Corpuscular Hemoglobin Concentration |
| RDW | Red Cell Distribution Width |
| RET | Reticulocytes (Absolute) |

**Supplemental Table 6:**

Male Hematology Results Part II

| Vehicle Control | Day(s) Relative to Start Date | PLT (10^9/L) | WBC (10^9/L) | NEUT (10^9/L) | LYMP (10^9/L) | MONO (10^9/L) | EOS (10^9/L) | BASO (10^9/L) |
|---|---|---|---|---|---|---|---|---|
| 1101 | -13 | 257 | 16.07 | 8.51 | 6.91 | 0.24 | 0.17 | 0.11 |
| | -6 | 297 | 18.52 | 5.88 | 10.99 | 0.47 | 0.83 | 0.16 |
| | 1 | 295 | 21.84 | 8.67 | 11.78 | 0.45 | 0.60 | 0.14 |
| | 2 | 270 | 21.24 | 7.36 | 12.58 | 0.41 | 0.49 | 0.18 |
| | 7 | 327 | 18.12 | 5.08 | 11.85 | 0.43 | 0.50 | 0.11 |
| | 14 | 386 | 20.46 | 8.21 | 10.86 | 0.43 | 0.63 | 0.16 |
| | 42 | 280 | 15.70 | 4.30 | 10.26 | 0.38 | 0.47 | 0.12 |

| AB001 0.1 mg/kg | Day(s) Relative to Start Date | PLT (10^9/L) | WBC (10^9/L) | NEUT (10^9/L) | LYMP (10^9/L) | MONO (10^9/L) | EOS (10^9/L) | BASO (10^9/L) |
|---|---|---|---|---|---|---|---|---|
| 1203 | -13 | 347 | 9.14 | 4.36 | 4.32 | 0.30 | 0.07 | 0.04 |
| | -6 | 436 | 11.74 | 1.82 | 8.93 | 0.49 | 0.30 | 0.11 |
| | 1 | 387 | 10.69 | 1.43 | 8.52 | 0.26 | 0.34 | 0.07 |
| | 2 | 369 | 11.81 | 6.25 | 4.95 | 0.37 | 0.14 | 0.05 |
| | 7 | 410 | 9.25 | 3.78 | 4.95 | 0.24 | 0.19 | 0.04 |
| | 14 | 436 | 8.93 | 2.54 | 5.80 | 0.27 | 0.25 | 0.03 |
| | 42 | 364 | 12.90 | 6.83 | 5.37 | 0.38 | 0.19 | 0.07 |

| AB001 1 mg/kg | Day(s) Relative to Start Date | PLT (10^9/L) | WBC (10^9/L) | NEUT (10^9/L) | LYMP (10^9/L) | MONO (10^9/L) | EOS (10^9/L) | BASO (10^9/L) |
|---|---|---|---|---|---|---|---|---|
| 1305 | -13 | 334 | 22.46 | 6.80 | 14.83 | 0.32 | 0.05 | 0.20 |
| | -6 | 375 | 36.24 | 7.27 | 26.77 | 0.72 | 0.33 | 0.54 |
| | -3 | 335 | 30.61 | 4.56 | 24.24 | 0.44 | 0.37 | 0.47 |
| | 1 | 386 | 35.27 | 4.59 | 29.04 | 0.25 | 0.31 | 0.56 |
| | 2 | 330 | 32.46 | 7.99 | 23.17 | 0.29 | 0.26 | 0.38 |
| | 7 | 356 | 33.46 | 11.88 | 20.27 | 0.33 | 0.28 | 0.38 |
| | 14 | 343 | 34.98 | 8.03 | 25.17 | 0.45 | 0.47 | 0.43 |
| | 42 | 306 | 28.18 | 4.86 | 21.89 | 0.49 | 0.22 | 0.32 |

| AB001<br>10 mg/kg | Day(s) Relative to Start Date | PLT (10^9/L) | WBC (10^9/L) | NEUT (10^9/L) | LYMP (10^9/L) | MONO (10^9/L) | EOS (10^9/L) | BASO (10^9/L) |
|---|---|---|---|---|---|---|---|---|
| 1407 | -13 | 375 | 14.70 | 4.79 | 9.52 | 0.18 | 0.05 | 0.09 |
| | -6 | 389 | 30.01 | 13.65 | 14.59 | 1.07 | 0.25 | 0.26 |
| | -3 | 382 | 16.90 | 1.49 | 14.57 | 0.37 | 0.18 | 0.16 |
| | 1 | 339 | 19.36 | 1.85 | 16.55 | 0.36 | 0.25 | 0.19 |
| | 2 | 302 | 17.83 | 4.11 | 12.91 | 0.29 | 0.21 | 0.16 |
| | 7 | 375 | 16.53 | 4.47 | 11.34 | 0.31 | 0.22 | 0.11 |
| | 14 | 443 | 26.94 | 5.32 | 20.19 | 0.50 | 0.41 | 0.30 |
| | 42 | 331 | 22.21 | 2.61 | 18.37 | 0.46 | 0.29 | 0.25 |

| AB001<br>100 mg/kg | Day(s) Relative to Start Date | PLT (10^9/L) | WBC (10^9/L) | NEUT (10^9/L) | LYMP (10^9/L) | MONO (10^9/L) | EOS (10^9/L) | BASO (10^9/L) |
|---|---|---|---|---|---|---|---|---|
| 1509 | -13 | 452 | 10.38 | 5.67 | 3.82 | 0.70 | 0.03 | 0.09 |
| | -6 | 494 | 25.43 | 8.46 | 15.40 | 0.65 | 0.25 | 0.36 |
| | 1 | 515 | 20.92 | 5.45 | 14.11 | 0.58 | 0.37 | 0.21 |
| | 2 | 457 | 33.35 | 25.95 | 6.15 | 0.93 | 0.10 | 0.15 |
| | 7 | 409 | 18.22 | 8.40 | 8.97 | 0.47 | 0.13 | 0.14 |
| | 14 | 599 | 20.36 | 10.09 | 9.39 | 0.45 | 0.19 | 0.15 |
| | 42 | 510 | 16.62 | 7.42 | 8.33 | 0.52 | 0.14 | 0.12 |

Abbreviations:

| | |
|---|---|
| PLT | Platelets |
| WBC | White Blood Cells |
| NEUT | Neutrophils (Absolute) |
| LYMP | Lymphocytes (Absolute) |
| MONO | Monocytes (Absolute) |
| EOS | Eosinophils (Absolute) |
| BASO | Basophils (Absolute) |

**Supplemental Table 7:**

Female Hematology Results Part I

| Vehicle Control | Day(s) Relative to Start Date | RBC (10^12/L) | HGB (g/L) | HCT (%) | MCV (fL) | MCH (pg) | MCHC (g/L) | RDW (%) | RET (10^9/L) |
|---|---|---|---|---|---|---|---|---|---|
| 2102 | -13 | 6.31 | 141 | 46.3 | 73.4 | 22.4 | 305 | 12.4 | 122.8 |
| | -6 | 6.75 | 148 | 49.8 | 73.9 | 21.9 | 296 | 11.9 | 114.0 |
| | 1 | 6.00 | 131 | 44.2 | 73.7 | 21.9 | 297 | 12.0 | 121.5 |
| | 2 | 5.32 | 116 | 39.0 | 73.3 | 21.7 | 296 | 12.2 | 117.1 |
| | 7 | 5.45 | 119 | 41.0 | 75.2 | 21.8 | 290 | 12.5 | 255.5 |
| | 14 | 5.77 | 126 | 42.0 | 72.8 | 21.8 | 300 | 12.2 | 196.7 |
| | 42 | 6.18 | 136 | 44.3 | 71.7 | 22.1 | 308 | 11.5 | 72.4 |

| AB001 0.1 mg/kg | Day(s) Relative to Start Date | RBC (10^12/L) | HGB (g/L) | HCT (%) | MCV (fL) | MCH (pg) | MCHC (g/L) | RDW (%) | RET (10^9/L) |
|---|---|---|---|---|---|---|---|---|---|
| 2204 | -13 | 5.15 | 129 | 42.2 | 82.1 | 25.0 | 305 | 14.4 | 99.3 |
| | -6 | 5.31 | 136 | 43.1 | 81.1 | 25.5 | 315 | 14.3 | 101.3 |
| | 1 | 4.90 | 127 | 40.1 | 81.7 | 25.9 | 317 | 14.2 | 95.9 |
| | 2 | 4.60 | 113 | 37.0 | 80.4 | 24.6 | 306 | 14.4 | 107.8 |
| | 7 | 4.59 | 114 | 36.9 | 80.5 | 24.8 | 308 | 14.3 | 238.0 |
| | 14 | 4.95 | 123 | 39.6 | 80.0 | 24.8 | 310 | 14.3 | 180.1 |
| | 42 | 5.47 | 135 | 43.2 | 79.0 | 24.7 | 313 | 13.6 | 99.0 |

| AB001 1 mg/kg | Day(s) Relative to Start Date | RBC (10^12/L) | HGB (g/L) | HCT (%) | MCV (fL) | MCH (pg) | MCHC (g/L) | RDW (%) | RET (10^9/L) |
|---|---|---|---|---|---|---|---|---|---|
| 2306 | -13 | 5.20 | 125 | 40.9 | 78.6 | 24.0 | 306 | 13.1 | 70.8 |
| | -6 | 5.06 | 123 | 40.1 | 79.2 | 24.4 | 308 | 13.0 | 77.4 |
| | 1 | 4.98 | 121 | 39.8 | 79.8 | 24.3 | 305 | 13.1 | 80.4 |
| | 2 | 4.48 | 108 | 35.4 | 78.9 | 24.1 | 305 | 13.3 | 74.3 |
| | 7 | 4.55 | 111 | 35.8 | 78.6 | 24.3 | 309 | 13.6 | 148.8 |
| | 14 | 4.97 | 121 | 38.9 | 78.4 | 24.3 | 311 | 13.1 | 97.7 |
| | 42 | 5.25 | 130 | 40.1 | 76.4 | 24.7 | 323 | 12.2 | 40.1 |

| AB001 10 mg/kg | Day(s) Relative to Start Date | RBC (10^12/L) | HGB (g/L) | HCT (%) | MCV (fL) | MCH (pg) | MCHC (g/L) | RDW (%) | RET (10^9/L) |
|---|---|---|---|---|---|---|---|---|---|
| 2408 | -13 | 5.23 | 128 | 40.9 | 78.3 | 24.4 | 312 | 12.1 | 77.8 |
| | -6 | 5.30 | 129 | 42.4 | 80.1 | 24.3 | 303 | 11.8 | 90.7 |
| | 1 | 5.08 | 126 | 39.7 | 78.2 | 24.9 | 318 | 12.2 | 84.3 |
| | 2 | 4.70 | 115 | 37.1 | 79.0 | 24.5 | 310 | 12.2 | 97.1 |
| | 7 | 4.50 | 109 | 34.8 | 77.4 | 24.3 | 314 | 12.4 | 177.5 |
| | 14 | 4.85 | 117 | 37.0 | 76.2 | 24.1 | 316 | 11.9 | 91.4 |
| | 42 | 5.28 | 131 | 39.7 | 75.2 | 24.7 | 329 | 11.5 | 46.0 |

| AB001 100 mg/kg | Day(s) Relative to Start Date | RBC (10^12/L) | HGB (g/L) | HCT (%) | MCV (fL) | MCH (pg) | MCHC (g/L) | RDW (%) | RET (10^9/L) |
|---|---|---|---|---|---|---|---|---|---|
| 2510 | -13 | 5.25 | 112 | 37.9 | 72.3 | 21.3 | 295 | 14.6 | 152.9 |
| | -6 | 5.42 | 113 | 38.9 | 71.8 | 20.9 | 291 | 14.0 | 141.6 |
| | 1 | 5.18 | 109 | 37.0 | 71.6 | 21.1 | 295 | 14.1 | 143.2 |
| | 2 | 4.67 | 97 | 34.5 | 73.9 | 20.8 | 282 | 14.1 | 141.3 |
| | 7 | 4.84 | 103 | 34.0 | 70.2 | 21.2 | 302 | 14.5 | 194.5 |
| | 14 | 5.10 | 108 | 36.5 | 71.5 | 21.1 | 295 | 14.1 | 197.4 |
| | 42 | 5.40 | 114 | 36.9 | 68.3 | 21.1 | 309 | 12.9 | 100.6 |

Abbreviations:

| | |
|---|---|
| RBC | Red Blood Cells |
| HGB | Hemoglobin |
| HCT | Hematocrit |
| MCV | Mean Corpuscular Volume |
| MCH | Mean Corpuscular Hemoglobin |
| MCHC | Mean Corpuscular Hemoglobin Concentration |
| RDW | Red Cell Distribution Width |
| RET | Reticulocytes (Absolute) |

**Supplemental Table 8:**

Female Hematology Results Part II

| Vehicle Control | Day(s) Relative to Start Date | PLT (10^9/L) | WBC (10^9/L) | NEUT (10^9/L) | LYMP (10^9/L) | MONO (10^9/L) | EOS (10^9/L) | BASO (10^9/L) |
|---|---|---|---|---|---|---|---|---|
| 2102 | -13 | 409 | 12.30 | 8.25 | 3.64 | 0.28 | 0.01 | 0.06 |
| | -6 | 465 | 17.31 | 7.58 | 8.49 | 0.71 | 0.26 | 0.13 |
| | 1 | 463 | 17.06 | 6.68 | 9.41 | 0.47 | 0.21 | 0.13 |
| | 2 | 401 | 19.30 | 10.82 | 7.45 | 0.65 | 0.16 | 0.09 |
| | 7 | 466 | 18.84 | 10.43 | 7.52 | 0.53 | 0.14 | 0.10 |
| | 14 | 498 | 22.49 | 11.69 | 9.34 | 0.80 | 0.30 | 0.17 |
| | 42 | 406 | 15.61 | 6.94 | 7.72 | 0.55 | 0.18 | 0.11 |

| AB001 0.1 mg/kg | Day(s) Relative to Start Date | PLT (10^9/L) | WBC (10^9/L) | NEUT (10^9/L) | LYMP (10^9/L) | MONO (10^9/L) | EOS (10^9/L) | BASO (10^9/L) |
|---|---|---|---|---|---|---|---|---|
| 2204 | -13 | 291 | 10.33 | 5.08 | 4.84 | 0.29 | 0.02 | 0.05 |
| | -6 | 336 | 17.92 | 4.68 | 11.78 | 0.85 | 0.26 | 0.19 |
| | 1 | 314 | 15.43 | 4.61 | 9.67 | 0.66 | 0.24 | 0.13 |
| | 2 | 299 | 12.04 | 5.17 | 6.24 | 0.33 | 0.15 | 0.07 |
| | 7 | 371 | 12.08 | 5.61 | 5.78 | 0.43 | 0.11 | 0.08 |
| | 14 | 343 | 15.24 | 5.98 | 8.20 | 0.66 | 0.18 | 0.11 |
| | 42 | 292 | 13.39 | 4.75 | 7.79 | 0.47 | 0.19 | 0.08 |

| AB001 1 mg/kg | Day(s) Relative to Start Date | PLT (10^9/L) | WBC (10^9/L) | NEUT (10^9/L) | LYMP (10^9/L) | MONO (10^9/L) | EOS (10^9/L) | BASO (10^9/L) |
|---|---|---|---|---|---|---|---|---|
| 2306 | -13 | 511 | 18.73 | 12.88 | 5.48 | 0.23 | 0.04 | 0.06 |
| | -6 | 459 | 19.87 | 6.05 | 12.81 | 0.52 | 0.19 | 0.17 |
| | 1 | 551 | 16.10 | 3.61 | 11.82 | 0.26 | 0.21 | 0.10 |
| | 2 | 464 | 22.11 | 11.37 | 10.04 | 0.38 | 0.13 | 0.11 |
| | 7 | 468 | 16.55 | 7.28 | 8.71 | 0.32 | 0.10 | 0.08 |
| | 14 | 462 | 19.52 | 6.17 | 12.31 | 0.47 | 0.31 | 0.15 |
| | 42 | 426 | 19.46 | 6.04 | 12.49 | 0.39 | 0.28 | 0.16 |

| AB001<br>10 mg/kg | Day(s) Relative to Start Date | PLT (10^9/L) | WBC (10^9/L) | NEUT (10^9/L) | LYMP (10^9/L) | MONO (10^9/L) | EOS (10^9/L) | BASO (10^9/L) |
|---|---|---|---|---|---|---|---|---|
| 2408 | -13 | 283 | 11.59 | 4.41 | 6.73 | 0.30 | 0.02 | 0.05 |
| | -6 | 304 | 17.57 | 6.30 | 10.31 | 0.44 | 0.20 | 0.17 |
| | 1 | 324 | 14.88 | 3.99 | 9.90 | 0.42 | 0.33 | 0.10 |
| | 2 | 297 | 18.24 | 7.51 | 9.92 | 0.42 | 0.17 | 0.10 |
| | 7 | 294 | 17.57 | 10.20 | 6.80 | 0.32 | 0.10 | 0.08 |
| | 14 | 307 | 17.15 | 5.12 | 10.95 | 0.58 | 0.29 | 0.10 |
| | 42 | 300 | 21.70 | 9.11 | 11.29 | 0.74 | 0.22 | 0.15 |

| AB001<br>100 mg/kg | Day(s) Relative to Start Date | PLT (10^9/L) | WBC (10^9/L) | NEUT (10^9/L) | LYMP (10^9/L) | MONO (10^9/L) | EOS (10^9/L) | BASO (10^9/L) |
|---|---|---|---|---|---|---|---|---|
| 2510 | -13 | 359 | 13.64 | 7.64 | 5.45 | 0.36 | 0.07 | 0.06 |
| | -6 | 452 | 21.25 | 8.50 | 11.41 | 0.55 | 0.51 | 0.14 |
| | 1 | 418 | 18.34 | 6.48 | 10.88 | 0.40 | 0.38 | 0.09 |
| | 2 | 389 | 19.60 | 10.71 | 8.31 | 0.28 | 0.13 | 0.08 |
| | 7 | 434 | 18.73 | 13.53 | 4.70 | 0.22 | 0.17 | 0.04 |
| | 14 | 433 | 20.69 | 10.11 | 9.67 | 0.31 | 0.35 | 0.11 |
| | 42 | 412 | 20.23 | 12.52 | 6.91 | 0.35 | 0.28 | 0.11 |

Abbreviations:

| | |
|---|---|
| PLT | Platelets |
| WBC | White Blood Cells |
| NEUT | Neutrophils (Absolute) |
| LYMP | Lymphocytes (Absolute) |
| MONO | Monocytes (Absolute) |
| EOS | Eosinophils (Absolute) |
| BASO | Basophils (Absolute) |

**Supplemental Table 9:**

Male Coagulation Panel

| Vehicle Control | Day(s) Relative to Start Date | PT (Seconds) | APTT (Seconds) | FIB (g/L) |
|---|---|---|---|---|
| 1101 | -13 | 10.8 | 16.7 | 2.03 |
| | -6 | 9.8 | 16.6 | 1.61 |
| | 2 | 9.8 | 16.7 | 2.22 |
| | 42 | 9.8 | 16.3 | 1.80 |

| AB001 0.1 mg/kg | Day(s) Relative to Start Date | PT (Seconds) | APTT (Seconds) | FIB (g/L) |
|---|---|---|---|---|
| 1203 | -13 | 9.8 | 18.7 | 1.84 |
| | -6 | 9.0 | 20.1 | 1.80 |
| | 2 | 9.1 | 18.5 | 2.17 |
| | 42 | 8.6 | 19.7 | 2.07 |

| AB001 1 mg/kg | Day(s) Relative to Start Date | PT (Seconds) | APTT (Seconds) | FIB (g/L) |
|---|---|---|---|---|
| 1305 | -13 | 10.7 | 18.6 | 2.65 |
| | -6 | 9.5 | 19.0 | 2.49 |
| | 2 | 9.6 | 18.9 | 2.78 |
| | 42 | 9.4 | 20.0 | 3.54 |

| AB001 10 mg/kg | Day(s) Relative to Start Date | PT (Seconds) | APTT (Seconds) | FIB (g/L) |
|---|---|---|---|---|
| 1407 | -13 | 9.3 | 16.9 | 2.53 |
| | -6 | 9.1 | 17.7 | 2.65 |
| | 2 | 9.1 | 16.8 | 2.78 |
| | 42 | 9.0 | 18.1 | 2.61 |

| AB001<br>100 mg/kg | Day(s) Relative to Start Date | PT<br>(Seconds) | APTT<br>(Seconds) | FIB<br>(g/L) |
|---|---|---|---|---|
| 1509 | -13 | 11.5 | 20.4 | 2.74 |
| | -6 | 10.2 | 19.4 | 2.11 |
| | 2 | 11.1 | 19.6 | 4.02 |
| | 42 | 10.6 | 19.8 | 2.31 |

Abbreviations:

PT Prothrombin Time

APTT Activated Partial Thromboplastin Time

FIB Fibrinogen

**Supplemental Table 10:**

Female Coagulation Panel

| Vehicle Control | Day(s) Relative to Start Date | PT (Seconds) | APTT (Seconds) | FIB (g/L) |
|---|---|---|---|---|
| 2102 | -13 | 10.8 | 17.4 | 2.19 |
| | -6 | 9.9 | 16.7 | 2.03 |
| | 2 | 10.1 | 17.6 | 2.93 |
| | 42 | 9.8 | 18.3 | 2.20 |

| AB001 0.1 mg/kg | Day(s) Relative to Start Date | PT (Seconds) | APTT (Seconds) | FIB (g/L) |
|---|---|---|---|---|
| 2204 | -13 | 10.2 | 16.8 | 2.35 |
| | -6 | 9.3 | 16.9 | 2.39 |
| | 2 | 9.4 | 17.5 | 2.93 |
| | 42 | 9.2 | 18.1 | 2.54 |

| AB001 1 mg/kg | Day(s) Relative to Start Date | PT (Seconds) | APTT (Seconds) | FIB (g/L) |
|---|---|---|---|---|
| 2306 | -13 | 9.3 | 15.9 | 2.32 |
| | -6 | 8.8 | 14.6 | 2.03 |
| | 2 | 9.3 | 15.7 | 2.57 |
| | 42 | 8.5 | 14.8 | 2.15 |

| AB001 10 mg/kg | Day(s) Relative to Start Date | PT (Seconds) | APTT (Seconds) | FIB (g/L) |
|---|---|---|---|---|
| 2408 | -13 | 10.6 | 18.2 | 1.93 |
| | -6 | 9.0 | 17.3 | 1.95 |
| | 2 | 9.2 | 17.9 | 2.11 |
| | 42 | 8.8 | 18.6 | 2.34 |

| AB001<br>100 mg/kg | Day(s) Relative to Start Date | PT<br>(Seconds) | APTT<br>(Seconds) | FIB<br>(g/L) |
|---|---|---|---|---|
| 2510 | -13 | 10.2 | 19.9 | 1.86 |
| | -6 | 9.5 | 19.1 | 1.80 |
| | 2 | 9.2 | 20.0 | 2.11 |
| | 42 | 9.3 | 17.4 | 2.26 |

Abbreviations:

| | |
|---|---|
| PT | Prothrombin Time |
| APTT | Activated Partial Thromboplastin Time |
| FIB | Fibrinogen |

**Supplemental Table 11:**

Male Chemistry Panel Part I

| Vehicle Control | Day(s) Relative to Start Date | ALT (U/L) | AST (U/L) | ALP (U/L) | GGT (U/L) | CK (U/L) | TBIL (µmol/L) | GLU (mmol/L) |
|---|---|---|---|---|---|---|---|---|
| 1101 | -13 | 32.9 | 45.5 | 1122 | 61 | 175 | 5.9 | 3.63 |
| | -6 | 20.7 | 32.0 | 897 | 54 | 222 | 2.5 | 7.07 |
| | 2 | 34.3 | 72.3 | 766 | 53 | 221 | 2.5 | 6.21 |
| | 7 | 24.5 | 33.8 | 749 | 49 | 143 | 1.8 | 6.03 |
| | 14 | 19.2 | 27.6 | 783 | 49 | 195 | 0.7 | 6.96 |
| | 42 | 20.9 | 32.3 | 946 | 59 | 142 | 2.1 | 5.51 |

| AB001 0.1 mg/kg | Day(s) Relative to Start Date | ALT (U/L) | AST (U/L) | ALP (U/L) | GGT (U/L) | CK (U/L) | TBIL (µmol/L) | GLU (mmol/L) |
|---|---|---|---|---|---|---|---|---|
| 1203 | -13 | 49.9 | 72.3 | 799 | 95 | 302 | 3.1 | 5.77 |
| | -6 | 28.0 | 39.9 | 745 | 84 | 505 | 1.5 | 7.41 |
| | 2 | 54.8 | 93.5 | 702 | 77 | 715 | 1.6 | 5.92 |
| | 7 | 29.7 | 38.1 | 747 | 71 | 210 | 0.6 | 6.44 |
| | 14 | 26.0 | 37.2 | 803 | 74 | 318 | 0.6 | 5.12 |
| | 42 | 24.8 | 40.3 | 862 | 77 | 326 | 0.8 | 6.15 |

| AB001 1 mg/kg | Day(s) Relative to Start Date | ALT (U/L) | AST (U/L) | ALP (U/L) | GGT (U/L) | CK (U/L) | TBIL (µmol/L) | GLU (mmol/L) |
|---|---|---|---|---|---|---|---|---|
| 1305 | -13 | 42.2 | 55.5 | 681 | 140 | 472 | 5.4 | 5.68 |
| | -6 | 28.7 | 37.2 | 534 | 106 | 313 | 3.0 | 6.84 |
| | 2 | 43.9 | 75.8 | 513 | 97 | 753 | 2.7 | 6.00 |
| | 7 | 63.4 | 60.0 | 508 | 90 | 745 | 2.6 | 5.34 |
| | 14 | 31.8 | 32.4 | 531 | 93 | 213 | 1.1 | 5.77 |
| | 42 | 23.1 | 34.6 | 529 | 89 | 295 | 2.6 | 6.17 |

| AB001 10 mg/kg | Day(s) Relative to Start Date | ALT (U/L) | AST (U/L) | ALP (U/L) | GGT (U/L) | CK (U/L) | TBIL (µmol/L) | GLU (mmol/L) |
|---|---|---|---|---|---|---|---|---|
| 1407 | -13 | 64.6 | 71.1 | 601 | 87 | 664 | 3.2 | 6.84 |
| | -6 | 62.7 | 51.4 | 565 | 87 | 416 | 2.6 | 6.50 |
| | 2 | 61.0 | 108.5 | 525 | 76 | 471 | 2.0 | 6.83 |
| | 7 | 51.4 | 51.4 | 490 | 70 | 359 | 2.3 | 6.30 |
| | 14 | 42.8 | 38.4 | 501 | 72 | 234 | 1.2 | 5.83 |
| | 42 | 38.7 | 38.8 | 569 | 82 | 257 | 1.8 | 6.97 |

| AB001 100 mg/kg | Day(s) Relative to Start Date | ALT (U/L) | AST (U/L) | ALP (U/L) | GGT (U/L) | CK (U/L) | TBIL (µmol/L) | GLU (mmol/L) |
|---|---|---|---|---|---|---|---|---|
| 1509 | -13 | 92.7 | 71.8 | 613 | 80 | 188 | 3.2 | 5.62 |
| | -6 | 55.7 | 34.4 | 499 | 69 | 184 | 1.4 | 7.21 |
| | 2 | 257.8 | 866.5 | 581 | 82 | 12135 | 2.9 | 7.19 |
| | 7 | 272.2 | 117.9 | 593 | 71 | 614 | 1.4 | 6.76 |
| | 14 | 86.2 | 44.8 | 680 | 83 | 179 | 0.7 | 7.43 |
| | 42 | 60.3 | 46.1 | 827 | 98 | 281 | 1.5 | 7.82 |

Abbreviations:

| | |
|---|---|
| ALT | Alanine Aminotransferase |
| AST | Aspartate Aminotransferase |
| ALP | Alkaline Phosphatase |
| GGT | Gamma Glutamyl Transferase |
| CK | Creatine Kinase |
| TBIL | Total Bilirubin |
| GLU | Glucose |

**Supplemental Table 12:**

Male Chemistry Panel Part II

| Vehicle Control | Day(s) Relative to Start Date | UREA (mmol/L) | CREA (µmol/L) | TG (mmol/L) | CHOL (mmol/L) | TP (g/L) | ALB (g/L) | GLOB (g/L) |
|---|---|---|---|---|---|---|---|---|
| 1101 | -13 | 8.6 | 68 | 0.44 | 3.61 | 81.7 | 45.2 | 36.5 |
| | -6 | 8.4 | 62 | 0.36 | 3.58 | 74.0 | 42.1 | 31.9 |
| | 2 | 8.2 | 64 | 0.32 | 3.33 | 73.3 | 42.4 | 30.9 |
| | 7 | 7.9 | 64 | 0.50 | 3.73 | 73.6 | 41.4 | 32.2 |
| | 14 | 8.8 | 66 | 0.42 | 3.92 | 72.5 | 40.8 | 31.7 |
| | 42 | 7.6 | 61 | 0.56 | 3.35 | 75.9 | 43.2 | 32.7 |

| AB001 0.1 mg/kg | Day(s) Relative to Start Date | UREA (mmol/L) | CREA (µmol/L) | TG (mmol/L) | CHOL (mmol/L) | TP (g/L) | ALB (g/L) | GLOB (g/L) |
|---|---|---|---|---|---|---|---|---|
| 1203 | -13 | 8.3 | 69 | 0.63 | 2.95 | 74.1 | 46.5 | 27.6 |
| | -6 | 7.7 | 63 | 0.71 | 3.59 | 75.0 | 45.0 | 30.0 |
| | 2 | 5.5 | 63 | 0.53 | 2.75 | 73.2 | 45.8 | 27.4 |
| | 7 | 6.9 | 67 | 0.52 | 3.07 | 73.8 | 44.9 | 28.9 |
| | 14 | 7.5 | 71 | 0.59 | 3.20 | 70.4 | 42.9 | 27.5 |
| | 42 | 7.2 | 69 | 1.09 | 3.41 | 75.0 | 46.0 | 29.0 |

| AB001 1 mg/kg | Day(s) Relative to Start Date | UREA (mmol/L) | CREA (µmol/L) | TG (mmol/L) | CHOL (mmol/L) | TP (g/L) | ALB (g/L) | GLOB (g/L) |
|---|---|---|---|---|---|---|---|---|
| 1305 | -13 | 7.0 | 67 | 0.35 | 3.33 | 82.8 | 48.4 | 34.4 |
| | -6 | 5.5 | 56 | 0.22 | 4.11 | 78.4 | 45.6 | 32.8 |
| | 2 | 5.1 | 59 | 0.26 | 3.29 | 78.5 | 45.5 | 33.0 |
| | 7 | 5.7 | 61 | 0.30 | 3.78 | 79.4 | 47.3 | 32.1 |
| | 14 | 5.5 | 60 | 0.22 | 4.05 | 76.4 | 46.0 | 30.4 |
| | 42 | 5.8 | 59 | 0.39 | 4.42 | 81.0 | 45.6 | 35.4 |

| AB001<br>10 mg/kg | Day(s) Relative to Start Date | UREA (mmol/L) | CREA (μmol/L) | TG (mmol/L) | CHOL (mmol/L) | TP (g/L) | ALB (g/L) | GLOB (g/L) |
|---|---|---|---|---|---|---|---|---|
| 1407 | -13 | 7.4 | 69 | 0.46 | 3.50 | 76.6 | 44.1 | 32.5 |
| | -6 | 6.5 | 65 | 0.98 | 3.84 | 79.6 | 45.6 | 34.0 |
| | 2 | 5.4 | 70 | 0.49 | 3.66 | 77.6 | 43.8 | 33.8 |
| | 7 | 6.3 | 74 | 0.39 | 3.62 | 75.7 | 45.1 | 30.6 |
| | 14 | 6.0 | 67 | 0.43 | 4.12 | 73.7 | 44.2 | 29.5 |
| | 42 | 6.2 | 74 | 1.20 | 3.94 | 74.5 | 44.9 | 29.6 |

| AB001<br>100 mg/kg | Day(s) Relative to Start Date | UREA (mmol/L) | CREA (μmol/L) | TG (mmol/L) | CHOL (mmol/L) | TP (g/L) | ALB (g/L) | GLOB (g/L) |
|---|---|---|---|---|---|---|---|---|
| 1509 | -13 | 6.5 | 72 | 0.35 | 3.64 | 89.9 | 42.8 | 47.1 |
| | -6 | 6.3 | 73 | 0.65 | 3.67 | 86.5 | 40.0 | 46.5 |
| | 2 | 6.0 | 71 | 0.44 | 3.11 | 83.3 | 42.0 | 41.3 |
| | 7 | 6.0 | 73 | 0.59 | 3.26 | 78.9 | 43.3 | 35.6 |
| | 14 | 6.4 | 70 | 0.48 | 4.20 | 77.8 | 45.5 | 32.3 |
| | 42 | 6.2 | 76 | 0.87 | 4.38 | 75.2 | 47.8 | 27.4 |

Abbreviations:

| | |
|---|---|
| UREA | Urea |
| CREA | Creatinine |
| TG | Triglycerides |
| CHOL | Total Cholesterol |
| TP | Total Protein |
| ALB | Albumin |
| GLOB | Globulin |

**Supplemental Table 13:**

Male Chemistry Panel Part III

| Vehicle Control | Day(s) Relative to Start Date | A/G | Na (mmol/L) | K (mmol/L) | Cl (mmol/L) | IgA (g/L) | IgG (g/L) | IgM (g/L) |
|---|---|---|---|---|---|---|---|---|
| 1101 | -13 | 1.2 | 153 | 4.45 | 108.3 | 1.69 | 11.96 | 0.76 |
| | -6 | 1.3 | 153 | 3.74 | 104.0 | 1.54 | 10.08 | 0.66 |
| | 2 | 1.4 | 154 | 4.14 | 108.1 | 1.43 | 9.25 | 0.59 |
| | 7 | 1.3 | 150 | 3.59 | 105.5 | 1.50 | 10.29 | 0.54 |
| | 14 | 1.3 | 148 | 4.04 | 106.3 | 1.46 | 9.80 | 0.54 |
| | 42 | 1.3 | 146 | 3.41 | 104.0 | 1.57 | 11.28 | 0.66 |

| AB001 0.1 mg/kg | Day(s) Relative to Start Date | A/G | Na (mmol/L) | K (mmol/L) | Cl (mmol/L) | IgA (g/L) | IgG (g/L) | IgM (g/L) |
|---|---|---|---|---|---|---|---|---|
| 1203 | -13 | 1.7 | 156 | 4.20 | 112.4 | 0.81 | 8.44 | 0.51 |
| | -6 | 1.5 | 155 | 5.03 | 106.7 | 0.93 | 8.27 | 0.63 |
| | 2 | 1.7 | 150 | 3.87 | 106.0 | 0.84 | 7.48 | 0.56 |
| | 7 | 1.6 | 150 | 3.82 | 106.8 | 0.86 | 7.41 | 0.52 |
| | 14 | 1.6 | 149 | 3.87 | 108.0 | 0.93 | 7.88 | 0.57 |
| | 42 | 1.6 | 147 | 3.58 | 104.7 | 1.09 | 9.06 | 0.64 |

| AB001 1 mg/kg | Day(s) Relative to Start Date | A/G | Na (mmol/L) | K (mmol/L) | Cl (mmol/L) | IgA (g/L) | IgG (g/L) | IgM (g/L) |
|---|---|---|---|---|---|---|---|---|
| 1305 | -13 | 1.4 | 155 | 4.52 | 111.5 | 0.95 | 11.38 | 1.49 |
| | -6 | 1.4 | 149 | 4.15 | 104.7 | 1.00 | 10.83 | 1.83 |
| | 2 | 1.4 | 148 | 3.81 | 106.0 | 0.91 | 10.62 | 1.69 |
| | 7 | 1.5 | 149 | 4.26 | 106.5 | 0.81 | 9.67 | 1.39 |
| | 14 | 1.5 | 149 | 4.01 | 106.7 | 0.56 | 8.91 | 1.13 |
| | 42 | 1.3 | 144 | 3.67 | 102.8 | 0.88 | 11.94 | 1.57 |

| AB001 10 mg/kg | Day(s) Relative to Start Date | A/G | Na (mmol/L) | K (mmol/L) | Cl (mmol/L) | IgA (g/L) | IgG (g/L) | IgM (g/L) |
|---|---|---|---|---|---|---|---|---|
| 1407 | -13 | 1.4 | 152 | 4.08 | 108.0 | 0.75 | 10.20 | 0.74 |
| | -6 | 1.3 | 152 | 3.60 | 103.1 | 0.84 | 9.93 | 0.85 |
| | 2 | 1.3 | 148 | 3.41 | 106.0 | 0.84 | 9.47 | 0.83 |
| | 7 | 1.5 | 147 | 3.49 | 105.8 | 0.69 | 8.96 | 0.74 |
| | 14 | 1.5 | 145 | 3.75 | 102.3 | 0.58 | 8.21 | 0.64 |
| | 42 | 1.5 | 146 | 3.07 | 98.7 | 0.66 | 9.34 | 0.60 |

| AB001 100 mg/kg | Day(s) Relative to Start Date | A/G | Na (mmol/L) | K (mmol/L) | Cl (mmol/L) | IgA (g/L) | IgG (g/L) | IgM (g/L) |
|---|---|---|---|---|---|---|---|---|
| 1509 | -13 | 0.9 | 155 | 4.72 | 106.1 | 1.03 | 17.31 | 1.65 |
| | -6 | 0.9 | 148 | 4.46 | 102.1 | 0.95 | 16.38 | 2.75 |
| | 2 | 1.0 | 150 | 4.51 | 103.9 | 0.76 | 14.11 | 1.70 |
| | 7 | 1.2 | 152 | 4.68 | 106.9 | 0.60 | 10.86 | 1.20 |
| | 14 | 1.4 | 153 | 5.03 | 104.6 | 0.44 | 9.21 | 0.82 |
| | 42 | 1.7 | 155 | 5.04 | 104.2 | 0.24 | 7.24 | 0.49 |

Abbreviations:

| | |
|---|---|
| A/G | Albumin/Globulin Ratio |
| Na | Sodium |
| K | Potassium Chloride |
| Cl | Chloride |
| IgA | Immunoglobulin A |
| IgG | Immunoglobulin G |
| IgM | Immunoglobulin M |

**Supplemental Table 14:**

Female Chemistry Panel Part I

| Vehicle Control | Day(s) Relative to Start Date | ALT (U/L) | AST (U/L) | ALP (U/L) | GGT (U/L) | CK (U/L) | TBIL (µmol/L) | GLU (mmol/L) |
|---|---|---|---|---|---|---|---|---|
| 2102 | -13 | 17.9 | 53.3 | 865 | 64 | 225 | 3.1 | 3.92 |
| | -6 | 11.5 | 36.3 | 715 | 57 | 270 | 1.7 | 7.15 |
| | 2 | 29.2 | 208.0 | 688 | 50 | 3564 | 1.0 | 5.01 |
| | 7 | 58.2 | 55.1 | 623 | 50 | 218 | 0.8 | 6.95 |
| | 14 | 14.4 | 35.4 | 718 | 52 | 201 | 0.4 | 7.58 |
| | 42 | 10.7 | 41.9 | 708 | 56 | 144 | 1.2 | 5.53 |

| AB001 0.1 mg/kg | Day(s) Relative to Start Date | ALT (U/L) | AST (U/L) | ALP (U/L) | GGT (U/L) | CK (U/L) | TBIL (µmol/L) | GLU (mmol/L) |
|---|---|---|---|---|---|---|---|---|
| 2204 | -13 | 43.0 | 49.3 | 365 | 55 | 225 | 3.1 | 3.67 |
| | -6 | 28.5 | 28.3 | 344 | 57 | 153 | 1.5 | 5.76 |
| | 2 | 52.6 | 91.0 | 330 | 54 | 803 | 1.8 | 5.47 |
| | 7 | 82.2 | 62.8 | 320 | 54 | 508 | 1.6 | 5.54 |
| | 14 | 33.9 | 31.4 | 353 | 56 | 185 | 0.7 | 5.01 |
| | 42 | 30.1 | 38.1 | 406 | 61 | 203 | 1.6 | 6.41 |

| AB001 1 mg/kg | Day(s) Relative to Start Date | ALT (U/L) | AST (U/L) | ALP (U/L) | GGT (U/L) | CK (U/L) | TBIL (µmol/L) | GLU (mmol/L) |
|---|---|---|---|---|---|---|---|---|
| 2306 | -13 | 41.1 | 60.4 | 404 | 80 | 715 | 3.5 | 4.90 |
| | -6 | 28.5 | 30.5 | 326 | 79 | 158 | 2.2 | 5.32 |
| | 2 | 52.8 | 94.5 | 320 | 80 | 753 | 1.0 | 6.12 |
| | 7 | 39.5 | 37.3 | 319 | 77 | 217 | 1.5 | 4.76 |
| | 14 | 31.2 | 32.5 | 332 | 81 | 170 | 1.0 | 5.82 |
| | 42 | 25.7 | 35.9 | 403 | 92 | 164 | 1.2 | 5.14 |

| AB001 10 mg/kg | Day(s) Relative to Start Date | ALT (U/L) | AST (U/L) | ALP (U/L) | GGT (U/L) | CK (U/L) | TBIL (µmol/L) | GLU (mmol/L) |
|---|---|---|---|---|---|---|---|---|
| 2408 | -13 | 53.1 | 71.0 | 680 | 56 | 394 | 4.2 | 4.50 |
| | -6 | 44.7 | 39.3 | 581 | 54 | 257 | 2.8 | 6.27 |
| | 2 | 51.2 | 71.7 | 562 | 54 | 331 | 2.6 | 7.90 |
| | 7 | 36.6 | 34.9 | 527 | 53 | 158 | 2.3 | 5.80 |
| | 14 | 40.7 | 33.8 | 524 | 55 | 166 | 1.8 | 6.63 |
| | 42 | 33.7 | 39.6 | 706 | 57 | 233 | 2.1 | 4.41 |

| AB001 100 mg/kg | Day(s) Relative to Start Date | ALT (U/L) | AST (U/L) | ALP (U/L) | GGT (U/L) | CK (U/L) | TBIL (µmol/L) | GLU (mmol/L) |
|---|---|---|---|---|---|---|---|---|
| 2510 | -13 | 29.7 | 62.0 | 397 | 67 | 154 | 3.2 | 5.88 |
| | -6 | 16.4 | 31.0 | 328 | 62 | 106 | 1.8 | 7.48 |
| | 2 | 21.8 | 81.9 | 360 | 62 | 153 | 0.9 | 7.50 |
| | 7 | 22.9 | 38.6 | 331 | 62 | 109 | 1.0 | 6.71 |
| | 14 | 19.0 | 36.0 | 331 | 62 | 123 | 0.9 | 7.36 |
| | 42 | 15.8 | 33.2 | 372 | 61 | 132 | 1.5 | 6.38 |

Abbreviations:

| | |
|---|---|
| ALT | Alanine Aminotransferase |
| AST | Aspartate Aminotransferase |
| ALP | Alkaline Phosphatase |
| GGT | Gamma Glutamyl Transferase |
| CK | Creatine Kinase |
| TBIL | Total Bilirubin |
| GLU | Glucose |

**Supplemental Table 15:**

Female Chemistry Panel Part II

| Vehicle Control | Day(s) Relative to Start Date | UREA (mmol/L) | CREA (µmol/L) | TG (mmol/L) | CHOL (mmol/L) | TP (g/L) | ALB (g/L) | GLOB (g/L) |
|---|---|---|---|---|---|---|---|---|
| 2102 | -13 | 7.4 | 37 | 0.49 | 2.47 | 78.1 | 45.4 | 32.7 |
| | -6 | 7.2 | 44 | 0.40 | 2.30 | 73.1 | 43.2 | 29.9 |
| | 2 | 5.6 | 36 | 0.29 | 2.07 | 69.2 | 40.5 | 28.7 |
| | 7 | 5.2 | 37 | 0.22 | 2.34 | 69.3 | 41.2 | 28.1 |
| | 14 | 6.0 | 35 | 0.30 | 2.72 | 70.2 | 40.7 | 29.5 |
| | 42 | 5.3 | 36 | 0.59 | 2.50 | 71.6 | 43.5 | 28.1 |

| AB001 0.1 mg/kg | Day(s) Relative to Start Date | UREA (mmol/L) | CREA (µmol/L) | TG (mmol/L) | CHOL (mmol/L) | TP (g/L) | ALB (g/L) | GLOB (g/L) |
|---|---|---|---|---|---|---|---|---|
| 2204 | -13 | 6.3 | 63 | 0.47 | 3.23 | 79.8 | 42.4 | 37.4 |
| | -6 | 6.9 | 59 | 0.78 | 3.38 | 76.6 | 42.2 | 34.4 |
| | 2 | 5.2 | 63 | 0.44 | 3.18 | 73.1 | 38.8 | 34.3 |
| | 7 | 6.9 | 63 | 0.58 | 3.40 | 78.0 | 42.2 | 35.8 |
| | 14 | 6.6 | 61 | 0.46 | 3.71 | 74.6 | 39.9 | 34.7 |
| | 42 | 7.4 | 65 | 0.88 | 3.50 | 79.5 | 44.4 | 35.1 |

| AB001 1 mg/kg | Day(s) Relative to Start Date | UREA (mmol/L) | CREA (µmol/L) | TG (mmol/L) | CHOL (mmol/L) | TP (g/L) | ALB (g/L) | GLOB (g/L) |
|---|---|---|---|---|---|---|---|---|
| 2306 | -13 | 5.9 | 54 | 0.38 | 3.69 | 79.5 | 43.7 | 35.8 |
| | -6 | 5.6 | 52 | 0.29 | 3.82 | 75.2 | 41.5 | 33.7 |
| | 2 | 4.9 | 49 | 0.19 | 3.42 | 75.6 | 43.1 | 32.5 |
| | 7 | 5.0 | 55 | 0.21 | 3.96 | 74.1 | 44.6 | 29.5 |
| | 14 | 5.8 | 50 | 0.24 | 4.42 | 73.4 | 43.7 | 29.7 |
| | 42 | 5.6 | 48 | 0.32 | 4.28 | 75.2 | 44.1 | 31.1 |

| AB001 10 mg/kg | Day(s) Relative to Start Date | UREA (mmol/L) | CREA (µmol/L) | TG (mmol/L) | CHOL (mmol/L) | TP (g/L) | ALB (g/L) | GLOB (g/L) |
|---|---|---|---|---|---|---|---|---|
| 2408 | -13 | 11.8 | 60 | 0.33 | 3.52 | 73.2 | 44.8 | 28.4 |
| | -6 | 7.8 | 56 | 0.39 | 3.97 | 73.8 | 43.7 | 30.1 |
| | 2 | 7.6 | 59 | 0.31 | 3.72 | 72.5 | 44.6 | 27.9 |
| | 7 | 7.2 | 59 | 0.18 | 4.03 | 72.1 | 45.9 | 26.2 |
| | 14 | 7.3 | 59 | 0.21 | 4.50 | 69.8 | 44.5 | 25.3 |
| | 42 | 9.0 | 65 | 0.49 | 4.78 | 70.9 | 44.2 | 26.7 |

| AB001 100 mg/kg | Day(s) Relative to Start Date | UREA (mmol/L) | CREA (µmol/L) | TG (mmol/L) | CHOL (mmol/L) | TP (g/L) | ALB (g/L) | GLOB (g/L) |
|---|---|---|---|---|---|---|---|---|
| 2510 | -13 | 9.8 | 73 | 0.63 | 2.42 | 78.2 | 40.7 | 37.5 |
| | -6 | 7.7 | 66 | 0.64 | 2.36 | 73.1 | 38.9 | 34.2 |
| | 2 | 6.6 | 64 | 0.62 | 2.25 | 74.0 | 38.8 | 35.2 |
| | 7 | 7.9 | 81 | 0.70 | 2.67 | 73.7 | 41.8 | 31.9 |
| | 14 | 8.2 | 65 | 0.49 | 2.68 | 70.6 | 42.4 | 28.2 |
| | 42 | 7.3 | 72 | 0.64 | 2.35 | 66.7 | 40.4 | 26.3 |

Abbreviations:

| | |
|---|---|
| UREA | Urea |
| CREA | Creatinine |
| TG | Triglycerides |
| CHOL | Total Cholestrol |
| TP | Total Protein |
| ALB | Albumin |
| GLOB | Globulin |

**Supplemental Table 16:**

Female Chemistry Panel Part III

| Vehicle Control | Day(s) Relative to Start Date | A/G | Na (mmol/L) | K (mmol/L) | Cl (mmol/L) | IgA (g/L) | IgG (g/L) | IgM (g/L) |
|---|---|---|---|---|---|---|---|---|
| 2102 | -13 | 1.4 | 153 | 4.10 | 109.4 | 0.89 | 9.86 | 0.62 |
| | -6 | 1.4 | 152 | 4.52 | 106.6 | 0.96 | 8.70 | 0.68 |
| | 2 | 1.4 | 148 | 3.61 | 108.6 | 0.84 | 7.70 | 0.57 |
| | 7 | 1.5 | 148 | 3.72 | 108.6 | 0.92 | 7.93 | 0.58 |
| | 14 | 1.4 | 148 | 3.40 | 107.5 | 1.00 | 8.28 | 0.66 |
| | 42 | 1.5 | 145 | 3.41 | 105.2 | 1.02 | 9.33 | 0.67 |

| AB001 0.1 mg/kg | Day(s) Relative to Start Date | A/G | Na (mmol/L) | K (mmol/L) | Cl (mmol/L) | IgA (g/L) | IgG (g/L) | IgM (g/L) |
|---|---|---|---|---|---|---|---|---|
| 2204 | -13 | 1.1 | 155 | 4.49 | 114.2 | 1.96 | 11.39 | 1.60 |
| | -6 | 1.2 | 150 | 4.64 | 107.6 | 1.98 | 10.79 | 1.81 |
| | 2 | 1.1 | 147 | 3.94 | 109.5 | 1.65 | 9.50 | 1.40 |
| | 7 | 1.2 | 150 | 4.10 | 110.7 | 1.68 | 10.24 | 1.46 |
| | 14 | 1.1 | 150 | 4.27 | 110.4 | 1.72 | 10.01 | 1.53 |
| | 42 | 1.3 | 145 | 3.52 | 104.5 | 2.06 | 12.23 | 1.85 |

| AB001 1 mg/kg | Day(s) Relative to Start Date | A/G | Na (mmol/L) | K (mmol/L) | Cl (mmol/L) | IgA (g/L) | IgG (g/L) | IgM (g/L) |
|---|---|---|---|---|---|---|---|---|
| 2306 | -13 | 1.2 | 151 | 4.25 | 110.9 | 1.39 | 9.85 | 1.59 |
| | -6 | 1.2 | 146 | 4.25 | 104.7 | 1.33 | 9.22 | 1.49 |
| | 2 | 1.3 | 145 | 4.10 | 107.7 | 1.25 | 8.86 | 1.34 |
| | 7 | 1.5 | 146 | 3.71 | 106.5 | 1.07 | 8.53 | 1.13 |
| | 14 | 1.5 | 146 | 4.17 | 105.2 | 0.87 | 8.30 | 0.90 |
| | 42 | 1.4 | 144 | 3.46 | 103.8 | 1.05 | 9.95 | 0.99 |

| AB001 10 mg/kg | Day(s) Relative to Start Date | A/G | Na (mmol/L) | K (mmol/L) | Cl (mmol/L) | IgA (g/L) | IgG (g/L) | IgM (g/L) |
|---|---|---|---|---|---|---|---|---|
| 2408 | -13 | 1.6 | 152 | 3.36 | 111.7 | 0.87 | 9.36 | 0.91 |
| | -6 | 1.5 | 150 | 4.26 | 105.1 | 0.90 | 9.12 | 0.96 |
| | 2 | 1.6 | 145 | 3.54 | 103.1 | 0.82 | 9.01 | 0.87 |
| | 7 | 1.8 | 145 | 3.34 | 105.3 | 0.63 | 8.01 | 0.65 |
| | 14 | 1.8 | 146 | 3.62 | 104.5 | 0.45 | 7.15 | 0.55 |
| | 42 | 1.7 | 144 | 3.28 | 102.2 | 0.35 | 9.04 | 0.47 |

| AB001 100 mg/kg | Day(s) Relative to Start Date | A/G | Na (mmol/L) | K (mmol/L) | Cl (mmol/L) | IgA (g/L) | IgG (g/L) | IgM (g/L) |
|---|---|---|---|---|---|---|---|---|
| 2510 | -13 | 1.1 | 152 | 3.68 | 112.5 | 0.96 | 12.59 | 0.95 |
| | -6 | 1.1 | 148 | 3.63 | 107.0 | 0.98 | 12.24 | 0.93 |
| | 2 | 1.1 | 146 | 3.77 | 107.3 | 0.95 | 13.16 | 0.86 |
| | 7 | 1.3 | 146 | 3.92 | 110.2 | 0.72 | 10.76 | 0.74 |
| | 14 | 1.5 | 147 | 3.64 | 109.0 | 0.47 | 9.24 | 0.54 |
| | 42 | 1.5 | 145 | 3.23 | 107.3 | 0.13 | 7.71 | 0.24 |

Abbreviations:

| | |
|---|---|
| A/G | Albumin/Globulin Ratio |
| Na | Sodium |
| K | Potassium Chloride |
| Cl | Chloride |
| IgA | Immunoglobulin A |
| IgG | Immunoglobulin G |
| IgM | Immunoglobulin M |

**Supplemental Table 17:**

Male Cytokine Panel Part I

| Vehicle Control | Day(s) Relative to Start Date | IL-2 (pg/mL) | IL-4 (pg/mL) | IL-6 (pg/mL) | IL-10 (pg/mL) | IFN-γ (pg/mL) |
|---|---|---|---|---|---|---|
| 1101 | 1 (PrD0M) | 0.000 | 0.000 | 0.000 | 0.000 | 0.000 |
| | 1 (1HPD) | 0.000 | 0.000 | 61.490 | 0.000 | 0.000 |
| | 1 (3HPD) | 0.000 | 0.000 | 13.070 | 0.000 | 0.000 |
| | 1 (8HPD) | 0.000 | 2.000 | 10.280 | 0.000 | 0.000 |
| | 2 | 0.000 | 0.420 | 0.000 | 0.000 | 0.000 |
| | 3 | 0.000 | 0.000 | 0.000 | 0.000 | 0.000 |
| | 7 | 0.000 | 0.000 | 0.000 | 0.000 | 0.000 |

| AB001 0.1 mg/kg | Day(s) Relative to Start Date | IL-2 (pg/mL) | IL-4 (pg/mL) | IL-6 (pg/mL) | IL-10 (pg/mL) | IFN-γ (pg/mL) |
|---|---|---|---|---|---|---|
| 1203 | 1 (PrD0M) | 0.000 | 0.000 | 0.000 | 0.000 | 0.000 |
| | 1 (1HPD) | 0.000 | 0.000 | 22.110 | 0.000 | 0.000 |
| | 1 (3HPD) | 0.000 | 0.000 | 16.130 | 0.000 | 0.000 |
| | 1 (8HPD) | 0.000 | 0.000 | 19.480 | 0.000 | 0.000 |
| | 2 | 0.000 | 0.000 | 1.990 | 0.000 | 0.000 |
| | 3 | 0.000 | 0.000 | 0.000 | 0.000 | 0.000 |
| | 7 | 0.000 | 0.000 | 0.000 | 0.000 | 0.000 |

| AB001 1 mg/kg | Day(s) Relative to Start Date | IL-2 (pg/mL) | IL-4 (pg/mL) | IL-6 (pg/mL) | IL-10 (pg/mL) | IFN-γ (pg/mL) |
|---|---|---|---|---|---|---|
| 1305 | 1 (PrD0M) | 0.000 | 0.000 | 0.000 | 0.000 | 0.000 |
| | 1 (1HPD) | 0.000 | 0.000 | 88.650 | 1.780 | 0.000 |
| | 1 (3HPD) | 0.000 | 0.000 | 28.720 | 0.000 | 0.000 |
| | 1 (8HPD) | 0.000 | 0.000 | 6.200 | 0.000 | 0.000 |
| | 2 | 0.000 | 0.000 | 0.000 | 0.000 | 0.000 |
| | 3 | 0.000 | 0.000 | 0.420 | 0.000 | 0.000 |
| | 7 | 0.000 | 0.000 | 0.000 | 0.000 | 0.000 |

| AB001<br>10 mg/kg | Day(s) Relative to Start Date | IL-2 (pg/mL) | IL-4 (pg/mL) | IL-6 (pg/mL) | IL-10 (pg/mL) | IFN-γ (pg/mL) |
|---|---|---|---|---|---|---|
| 1407 | 1 (PrD0M) | 0.000 | 0.000 | 0.000 | 0.000 | 0.000 |
| | 1 (1HPD) | 0.000 | 0.000 | 23.140 | 0.000 | 0.000 |
| | 1 (3HPD) | 0.000 | 0.000 | 14.430 | 0.000 | 0.000 |
| | 1 (8HPD) | 0.000 | 0.000 | 0.000 | 0.000 | 0.000 |
| | 2 | 0.000 | 0.000 | 0.000 | 0.000 | 0.000 |
| | 3 | 0.000 | 0.000 | 0.000 | 0.000 | 0.000 |
| | 7 | 0.000 | 0.000 | 0.000 | 0.000 | 0.000 |

| AB001<br>100 mg/kg | Day(s) Relative to Start Date | IL-2 (pg/mL) | IL-4 (pg/mL) | IL-6 (pg/mL) | IL-10 (pg/mL) | IFN-γ (pg/mL) |
|---|---|---|---|---|---|---|
| 1509 | 1 (PrD0M) | 0.000 | 0.000 | 0.000 | 0.000 | 0.000 |
| | 1 (1HPD) | 0.000 | 3.240 | 108.130 | 86.190 | 10.180 |
| | 1 (3HPD) | 0.000 | 3.060 | 103.970 | 0.000 | 13.000 |
| | 1 (8HPD) | 0.000 | 0.870 | 52.660 | 25.630 | 8.950 |
| | 2 | 0.000 | 0.000 | 30.330 | 0.000 | 6.820 |
| | 3 | 0.000 | 0.000 | 3.150 | 20.440 | 0.000 |
| | 7 | 0.000 | 0.000 | 0.000 | 0.000 | 0.000 |

| Abbreviation | Description |
|---|---|
| PrD0M | Pre Dose 0 min |
| 1HPD | 1 Hr Post Dose |
| 3HPD | 3 Hr Post Dose |
| 8HPD | 8 Hr Post Dose |

**Supplemental Table 18:**

Male Cytokine Panel Part II

| Vehicle Control | Day(s) Relative to Start Date | TNF-α (pg/mL) | IL-17A (pg/mL) | IL-5 (pg/mL) | IL-13 (pg/mL) | IL-21 (pg/mL) |
|---|---|---|---|---|---|---|
| 1101 | 1 (PrD0M) | 0.000 | 0.000 | 0.00 | 0.00 | 0.00 |
| | 1 (1HPD) | 0.000 | 0.000 | 0.00 | 0.00 | 0.00 |
| | 1 (3HPD) | 0.000 | 0.000 | 0.00 | 0.00 | 0.00 |
| | 1 (8HPD) | 0.000 | 0.000 | 0.00 | 0.00 | 0.00 |
| | 2 | 0.000 | 0.000 | 0.00 | 0.00 | 0.00 |
| | 3 | 0.000 | 0.000 | 0.00 | 0.00 | 0.00 |
| | 7 | 0.000 | 0.000 | 0.00 | 0.00 | 0.00 |

| AB001 0.1 mg/kg | Day(s) Relative to Start Date | TNF-α (pg/mL) | IL-17A (pg/mL) | IL-5 (pg/mL) | IL-13 (pg/mL) | IL-21 (pg/mL) |
|---|---|---|---|---|---|---|
| 1203 | 1 (PrD0M) | 0.000 | 0.000 | 0.00 | 0.00 | 0.00 |
| | 1 (1HPD) | 0.000 | 0.000 | 0.00 | 0.00 | 0.00 |
| | 1 (3HPD) | 0.000 | 0.000 | 0.00 | 0.00 | 0.00 |
| | 1 (8HPD) | 0.000 | 0.000 | 0.00 | 0.00 | 0.00 |
| | 2 | 0.000 | 0.000 | 0.00 | 0.00 | 0.00 |
| | 3 | 0.000 | 0.000 | 0.00 | 0.00 | 0.00 |
| | 7 | 0.000 | 0.000 | 0.00 | 0.00 | 0.00 |

| AB001 1 mg/kg | Day(s) Relative to Start Date | TNF-α (pg/mL) | IL-17A (pg/mL) | IL-5 (pg/mL) | IL-13 (pg/mL) | IL-21 (pg/mL) |
|---|---|---|---|---|---|---|
| 1305 | 1 (PrD0M) | 0.000 | 0.000 | 0.00 | 0.00 | 0.00 |
| | 1 (1HPD) | 0.000 | 0.000 | 0.00 | 0.00 | 0.00 |
| | 1 (3HPD) | 0.000 | 0.000 | 0.00 | 0.00 | 0.00 |
| | 1 (8HPD) | 0.000 | 0.000 | 0.00 | 0.00 | 0.00 |
| | 2 | 0.000 | 0.000 | 0.00 | 0.00 | 0.00 |
| | 3 | 0.000 | 0.000 | 0.00 | 0.00 | 0.00 |
| | 7 | 0.000 | 0.000 | 0.00 | 0.00 | 0.00 |

| AB001<br>10 mg/kg<br>Day(s) Relative to Start Date | | TNF-α (pg/mL) | IL-17A (pg/mL) | IL-5 (pg/mL) | IL-13 (pg/mL) | IL-21 (pg/mL) |
|---|---|---|---|---|---|---|
| 1407 | 1 (PrD0M) | 0.000 | 0.000 | 0.00 | 0.00 | 0.00 |
| | 1 (1HPD) | 0.000 | 0.000 | 0.00 | 0.00 | 0.00 |
| | 1 (3HPD) | 0.000 | 0.000 | 0.00 | 0.00 | 0.00 |
| | 1 (8HPD) | 0.000 | 0.000 | 0.00 | 0.00 | 0.00 |
| | 2 | 0.000 | 0.000 | 0.00 | 0.00 | 0.00 |
| | 3 | 0.000 | 0.000 | 0.00 | 0.00 | 0.00 |
| | 7 | 0.000 | 0.000 | 0.00 | 0.00 | 0.00 |

| AB001<br>100 mg/kg<br>Day(s) Relative to Start Date | | TNF-α (pg/mL) | IL-17A (pg/mL) | IL-5 (pg/mL) | IL-13 (pg/mL) | IL-21 (pg/mL) |
|---|---|---|---|---|---|---|
| 1509 | 1 (PrD0M) | 0.000 | 0.000 | 0.00 | 0.00 | 0.00 |
| | 1 (1HPD) | 0.000 | 6.660 | 0.00 | 0.00 | 0.00 |
| | 1 (3HPD) | 0.000 | 3.330 | 0.00 | 0.00 | 0.00 |
| | 1 (8HPD) | 0.000 | 5.190 | 0.00 | 0.00 | 0.00 |
| | 2 | 0.000 | 0.000 | 0.00 | 0.00 | 0.00 |
| | 3 | 0.000 | 0.000 | 0.00 | 0.00 | 0.00 |
| | 7 | 0.000 | 0.000 | 0.00 | 0.00 | 0.00 |

| Abbreviation | Description |
|---|---|
| PrD0M | Pre Dose 0 min |
| 1HPD | 1 Hr Post Dose |
| 3HPD | 3 Hr Post Dose |
| 8HPD | 8 Hr Post Dose |

**Supplemental Table 19:**

Female Cytokine Panel Part I

| Vehicle Control | Day(s) Relative to Start Date | IL-2 (pg/mL) | IL-4 (pg/mL) | IL-6 (pg/mL) | IL-10 (pg/mL) | IFN-γ (pg/mL) |
|---|---|---|---|---|---|---|
| 2102 | 1 (PrD0M) | 0.000 | 0.420 | 0.000 | 0.000 | 0.000 |
| | 1 (1HPD) | 0.000 | 3.240 | 46.950 | 7.560 | 0.000 |
| | 1 (3HPD) | 0.000 | 4.820 | 34.150 | 7.560 | 8.110 |
| | 1 (8HPD) | 0.000 | 5.030 | 17.610 | 1.780 | 8.110 |
| | 2 | 0.000 | 0.720 | 0.000 | 0.000 | 0.000 |
| | 3 | 0.000 | 0.000 | 10.520 | 0.000 | 0.000 |
| | 7 | 0.000 | 1.340 | 0.000 | 20.440 | 0.000 |

| AB001 0.1 mg/kg | Day(s) Relative to Start Date | IL-2 (pg/mL) | IL-4 (pg/mL) | IL-6 (pg/mL) | IL-10 (pg/mL) | IFN-γ (pg/mL) |
|---|---|---|---|---|---|---|
| 2204 | 1 (PrD0M) | 0.000 | 0.000 | 0.000 | 0.000 | 0.000 |
| | 1 (1HPD) | 0.000 | 0.000 | 64.900 | 18.750 | 0.000 |
| | 1 (3HPD) | 0.000 | 0.000 | 27.940 | 0.000 | 0.000 |
| | 1 (8HPD) | 0.000 | 2.170 | 8.170 | 0.000 | 0.000 |
| | 2 | 0.000 | 2.340 | 0.000 | 0.000 | 0.000 |
| | 3 | 0.000 | 0.720 | 0.000 | 10.630 | 0.000 |
| | 7 | 0.000 | 0.000 | 0.000 | 0.000 | 0.000 |

| AB001 1 mg/kg | Day(s) Relative to Start Date | IL-2 (pg/mL) | IL-4 (pg/mL) | IL-6 (pg/mL) | IL-10 (pg/mL) | IFN-γ (pg/mL) |
|---|---|---|---|---|---|---|
| 2306 | 1 (PrD0M) | 0.000 | 2.170 | 0.000 | 0.000 | 5.460 |
| | 1 (1HPD) | 0.000 | 0.000 | 55.080 | 1.780 | 0.000 |
| | 1 (3HPD) | 0.000 | 0.000 | 29.120 | 0.000 | 0.000 |
| | 1 (8HPD) | 0.000 | 0.870 | 15.270 | 0.000 | 4.460 |
| | 2 | 0.000 | 0.280 | 0.000 | 0.000 | 3.350 |
| | 3 | 0.000 | 0.000 | 0.000 | 0.000 | 0.000 |
| | 7 | 0.000 | 2.520 | 0.000 | 0.000 | 8.530 |

| AB001<br>10 mg/kg | Day(s) Relative to Start Date | IL-2 (pg/mL) | IL-4 (pg/mL) | IL-6 (pg/mL) | IL-10 (pg/mL) | IFN-γ (pg/mL) |
|---|---|---|---|---|---|---|
| 2408 | 1 (PrD0M) | 0.000 | 0.000 | 0.000 | 0.000 | 0.000 |
| | 1 (1HPD) | 0.000 | 0.000 | 13.070 | 0.000 | 0.000 |
| | 1 (3HPD) | 0.000 | 0.000 | 0.000 | 0.000 | 0.000 |
| | 1 (8HPD) | 0.000 | 0.000 | 0.000 | 0.000 | 0.000 |
| | 2 | 0.000 | 0.000 | 0.000 | 0.000 | 0.000 |
| | 3 | 0.000 | 2.170 | 0.000 | 0.000 | 0.000 |
| | 7 | 0.000 | 0.000 | 0.000 | 0.000 | 0.000 |

| AB001<br>100 mg/kg | Day(s) Relative to Start Date | IL-2 (pg/mL) | IL-4 (pg/mL) | IL-6 (pg/mL) | IL-10 (pg/mL) | IFN-γ (pg/mL) |
|---|---|---|---|---|---|---|
| 2510 | 1 (PrD0M) | 0.000 | 0.000 | 0.000 | 0.000 | 0.000 |
| | 1 (1HPD) | 0.000 | 1.180 | 32.840 | 0.000 | 0.000 |
| | 1 (3HPD) | 0.000 | 2.170 | 7.280 | 0.000 | 8.110 |
| | 1 (8HPD) | 0.000 | 0.000 | 0.000 | 0.000 | 0.000 |
| | 2 | 0.000 | 0.000 | 0.000 | 0.000 | 0.000 |
| | 3 | 0.000 | 0.000 | 0.000 | 0.000 | 0.000 |
| | 7 | 0.000 | 2.170 | 0.000 | 4.610 | 10.180 |

| Abbreviation | Description |
|---|---|
| PrD0M | Pre Dose 0 min |
| 1HPD | 1 Hr Post Dose |
| 3HPD | 3 Hr Post Dose |
| 8HPD | 8 Hr Post Dose |

**Supplemental Table 20:**

Female Cytokine Panel Part II

| Vehicle Control | Day(s) Relative to Start Date | TNF-α (pg/mL) | IL-17A (pg/mL) | IL-5 (pg/mL) | IL-13 (pg/mL) | IL-21 (pg/mL) |
|---|---|---|---|---|---|---|
| 2102 | 1 (PrD0M) | 0.000 | 0.000 | 0.00 | 0.00 | 0.00 |
| | 1 (1HPD) | 0.000 | 5.960 | 0.00 | 0.00 | 0.00 |
| | 1 (3HPD) | 0.000 | 10.280 | 0.00 | 0.00 | 0.00 |
| | 1 (8HPD) | 0.000 | 9.720 | 0.00 | 0.00 | 0.00 |
| | 2 | 0.000 | 0.000 | 0.00 | 0.00 | 0.00 |
| | 3 | 0.000 | 0.000 | 0.00 | 0.00 | 0.00 |
| | 7 | 0.000 | 13.950 | 0.00 | 0.00 | 0.00 |

| AB001 0.1 mg/kg | Day(s) Relative to Start Date | TNF-α (pg/mL) | IL-17A (pg/mL) | IL-5 (pg/mL) | IL-13 (pg/mL) | IL-21 (pg/mL) |
|---|---|---|---|---|---|---|
| 2204 | 1 (PrD0M) | 0.000 | 0.000 | 0.00 | 0.00 | 0.00 |
| | 1 (1HPD) | 0.000 | 5.190 | 0.00 | 0.00 | 0.00 |
| | 1 (3HPD) | 0.000 | 0.000 | 0.00 | 0.00 | 0.00 |
| | 1 (8HPD) | 0.000 | 0.000 | 0.00 | 0.00 | 0.00 |
| | 2 | 0.000 | 0.000 | 0.00 | 0.00 | 0.00 |
| | 3 | 0.000 | 4.340 | 0.00 | 0.00 | 0.00 |
| | 7 | 0.000 | 0.000 | 0.00 | 0.00 | 0.00 |

| AB001 1 mg/kg | Day(s) Relative to Start Date | TNF-α (pg/mL) | IL-17A (pg/mL) | IL-5 (pg/mL) | IL-13 (pg/mL) | IL-21 (pg/mL) |
|---|---|---|---|---|---|---|
| 2306 | 1 (PrD0M) | 0.000 | 10.830 | 0.00 | 0.00 | 0.00 |
| | 1 (1HPD) | 0.000 | 0.000 | 0.00 | 0.00 | 0.00 |
| | 1 (3HPD) | 0.000 | 0.000 | 0.00 | 0.00 | 0.00 |
| | 1 (8HPD) | 0.000 | 8.560 | 0.00 | 0.00 | 0.00 |
| | 2 | 0.000 | 7.330 | 0.00 | 0.00 | 0.00 |
| | 3 | 0.000 | 0.000 | 0.00 | 0.00 | 0.00 |
| | 7 | 0.000 | 11.890 | 0.00 | 0.00 | 0.00 |

| AB001<br>10 mg/kg | Day(s) Relative to Start Date | TNF-α (pg/mL) | IL-17A (pg/mL) | IL-5 (pg/mL) | IL-13 (pg/mL) | IL-21 (pg/mL) |
|---|---|---|---|---|---|---|
| 2408 | 1 (PrD0M) | 0.000 | 0.000 | 0.00 | 0.00 | 0.00 |
| | 1 (1HPD) | 0.000 | 0.000 | 0.00 | 0.00 | 0.00 |
| | 1 (3HPD) | 0.000 | 0.000 | 0.00 | 0.00 | 0.00 |
| | 1 (8HPD) | 0.000 | 0.000 | 0.00 | 0.00 | 0.00 |
| | 2 | 0.000 | 0.000 | 0.00 | 0.00 | 0.00 |
| | 3 | 0.000 | 0.000 | 0.00 | 0.00 | 0.00 |
| | 7 | 0.000 | 0.000 | 0.00 | 0.00 | 0.00 |

| AB001<br>100 mg/kg | Day(s) Relative to Start Date | TNF-α (pg/mL) | IL-17A (pg/mL) | IL-5 (pg/mL) | IL-13 (pg/mL) | IL-21 (pg/mL) |
|---|---|---|---|---|---|---|
| 2510 | 1 (PrD0M) | 0.000 | 0.000 | 0.00 | 0.00 | 0.00 |
| | 1 (1HPD) | 0.000 | 5.960 | 0.00 | 0.00 | 0.00 |
| | 1 (3HPD) | 0.000 | 13.440 | 0.00 | 0.00 | 0.00 |
| | 1 (8HPD) | 0.000 | 0.000 | 0.00 | 0.00 | 0.00 |
| | 2 | 0.000 | 0.000 | 0.00 | 0.00 | 0.00 |
| | 3 | 0.000 | 0.000 | 0.00 | 0.00 | 0.00 |
| | 7 | 0.000 | 11.890 | 0.00 | 0.00 | 0.00 |

| Abbreviation | Description |
|---|---|
| PrD0M | Pre Dose 0 min |
| 1HPD | 1 Hr Post Dose |
| 3HPD | 3 Hr Post Dose |
| 8HPD | 8 Hr Post Dose |
